# HELZ is a RNA-DNA helicase that resolves R loops to facilitate homologous recombination repair

**DOI:** 10.1101/2023.12.14.571747

**Authors:** Ramona Haji-Seyed-Javadi, Allyson E. Koyen, Sandip K. Rath, Bo Wu, Matthew Z. Madden, Yingzi Hou, Priya Kapoor-Vazirani, Meili Aiello, Anthony R. Sanchez, Nho Cong Luong, Fatmata Sesay, John S. Kim, Tony Tan, Seohyun Kim, Boya Gao, Boying S. Song, Anna M. Kenney, Erin C. Connolly, Lily Yang, Blerta Xhemalce, Xiaoxian Li, Jeffrey M. Switchenko, Xiaofeng Yang, Zachary S. Buchwald, Xingming Deng, Kyle M. Miller, Bing Yao, Li Lan, Weixing Zhao, David S. Yu

**Affiliations:** Department of Radiation Oncology and Winship Cancer Institute, Emory University School of Medicine, Atlanta, GA 30322, USA; Department of Biochemistry and Structural Biology, University of Texas Health Science Center at San Antonio, San Antonio, TX 78229, USA; Department of Human Genetics and Winship Cancer Institute,, Atlanta, GA 30322, USA; Department of Molecular Genetics and Microbiology, Duke University School of Medicine, Durham, NC 27710, USA; Department of Pediatrics and Winship Cancer Institute, Emory University School of Medicine, Atlanta, GA 30322, USA; Department of Surgery and Winship Cancer Institute, Emory University School of Medicine, Atlanta, GA 30322, USA; Department of Biochemistry and Winship Cancer Institute, Emory University School of Medicine, Atlanta, GA 30322, USA; Department of Pathology and Laboratory Medicine and Winship Cancer Institute, Emory University School of Medicine, Atlanta, GA 30322, USA; Department of Biostatistics and Bioinformatics and Winship Cancer Institute, Emory University School of Medicine, Atlanta, GA 30322, USA

## Abstract

R loop homeostasis is critical for DNA double-strand break (DSB) repair; however, how R loops are resolved in this context is poorly understood. Here, we define HELZ as a unique RNA-DNA helicase that resolves R loops to facilitate homologous recombination (HR) repair. From a synthetic lethal etoposide resistance siRNA screen, we found that HELZ depletion causes R loop-mediated hypersensitivity to DSB-inducing agents, and HELZ localizes and binds to DSBs. HELZ preferentially binds to and unwinds RNA-DNA hybrids with 5’ssRNA overhangs to promote R loop resolution genome-wide and at DSBs. Interestingly, HELZ facilitates BRCA1 recruitment to DSBs by preventing R loop accumulation, thereby promoting DNA end resection and HR to prevent R loop mediated genomic instability. Our findings define a role for HELZ in resolving R loops critical for HR that promotes genome stability and governs DSB-inducing agent resistance.

## Introduction

R loops are nucleic acids structures consisting of a RNA-DNA hybrid with a displaced single-stranded DNA (ssDNA). R loops are prevalent, occupying about 5% of the human genome^1^, and can form in multiple contexts, most frequently when a nascent RNA transcript hybridizes with the template ssDNA^2,3^. R loops are involved in diverse cellular processes, including transcription, DNA replication, DNA repair, telomere homeostasis, histone modification, and immunoglobulin class switch recombination^2–8^. While R loops play an important role in many physiological functions, dysregulation of R loop homeostasis resulting in unresolved or excessive accumulation of R loops can lead to DNA damage and genome instability^2–8^. Dysregulation of R loop homeostasis is associated with a number of diseases, including autoimmune disease, neurological disorders, and cancer^9^. Thus, the formation, resolution, and prevention of R loops must be tightly regulated.

DNA double-strand breaks (DSB) are highly cytotoxic lesions which must be repaired in order to preserve genome integrity^10–12^. DSBs are repaired predominantly by two major pathways: homologous recombination (HR), which involves repair using a sister chromatid as a template and is error-free, and non-homologous end joining (NHEJ), which involves ligation of broken DNA ends and is error-prone. HR is initiated by resection of broken DNA ends to generate short 3’ ssDNA overhangs, which is facilitated by the BRCA1 breast tumor suppressor protein^13–18^. R loops and/or RNA-DNA hybrids have been reported at DSB sites^19–25^ and can both promote^20,22–24^ and impair DSB repair^21,26–29^ in a context-dependent manner. However, the precise mechanisms by which R loops are resolved to promote DSB repair are not well understood.

Helicase with Zinc Finger (HELZ) is a member of the superfamily I (SF1) class of RNA helicases that alter the conformation of RNA by unwinding double-stranded regions^30^. Despite *HELZ* being cloned over 30 years ago^40^, very little is known about its function with only two reported HELZ-focused manuscripts indicating a helicase-independent role in protein translation and mRNA stability^31,32^. HELZ promotes cell proliferation, translation initiation, and ribosomal protein 6 phosphorylation^32^ and interacts with the carbon catabolite repressor 4-negative on TATA box (CCR4-NOT) deadenylase complex to cause decay of bound mRNAs^31^. HELZ is evolutionarily conserved in metazoa and its paralogs include SETX, AQR, DNA2, and UPF1, which have been implicated in DNA repair and/or R loop regulation^26,33–36^. HELZ contains an amino-terminal C3H1-type zinc finger motif, a Walker A motif conferring ATP binding, a Walker B DEAA [Asp, Glu, Ala, Ala] box helicase motif, and an unstructured carboxyl-terminal region with a conserved polyA binding protein interacting motif 2 (PAM2); however, HELZ’s biochemical activity as a helicase has not been established. Moreover, HELZ has not previously been characterized in regulating R loops or DNA repair. In this study, we define a role for HELZ in resolving R loops critical for DSB repair by HR to promote genome stability and govern resistance to DSB-inducing agents.

## Results

### A synthetic lethal siRNA screen targeting nuclear enzymes identifies HELZ as a mediator of etoposide resistance

To identify genes critical for governing resistance to etoposide, a topoisomerase II inhibitor and chemotherapeutic drug, we performed an etoposide sensitivity screen with a siRNA library targeting 1,006 annotated nuclear enzymes in H128 small cell lung cancer (SCLC) cells, derived from a treatment-refractory tumor (Fig. 1a). Sensitivity results from the screen are shown as a plot of −Log10 (p-value) against -strictly standardized mean difference (SSMD) (Fig. 1b). 61 etoposide resistance genes were identified using the following criteria: an average cell viability < 0.7, an average SSMD < −2, and a two-tailed t-test p-value < 0.05 (Supplementary Table S1-2). The Z-factor of the screen, an indicator of screen quality, was 0.534, which is in the excellent range, indicating that our screen is robust. Importantly, the siATR and siATRIP positive controls behaved as expected. (Fig. 1b). To characterize common pathways and potential interactions in these etoposide resistance genes, we performed Reactome pathway analysis^37^ (Supplementary Fig. S1), Metacore pathway map and process network enrichment analysis (https://clarivate.com/products/metacore) (Supplementary Fig. S2), and Gene Oncology (GO) network analysis^38^ (Supplementary Fig. S3), which showed enrichment of genes involved in multiple pathways, including a number in the DNA damage response (DDR): *RAD9*^39^, *HUS1*^40^, *RNF8*^41,42^, *PKNP*^43^, *SMUG1*^44^, *ADAR*^45^, *SENP2*^46,47^, and *HELLS*^48^, demonstrating that our screen can yield DDR proteins expected to mediate etoposide resistance. *HELZ*, which has not previously been characterized in the DDR, was also identified as a novel etoposide resistance gene.

**Figure 1.**
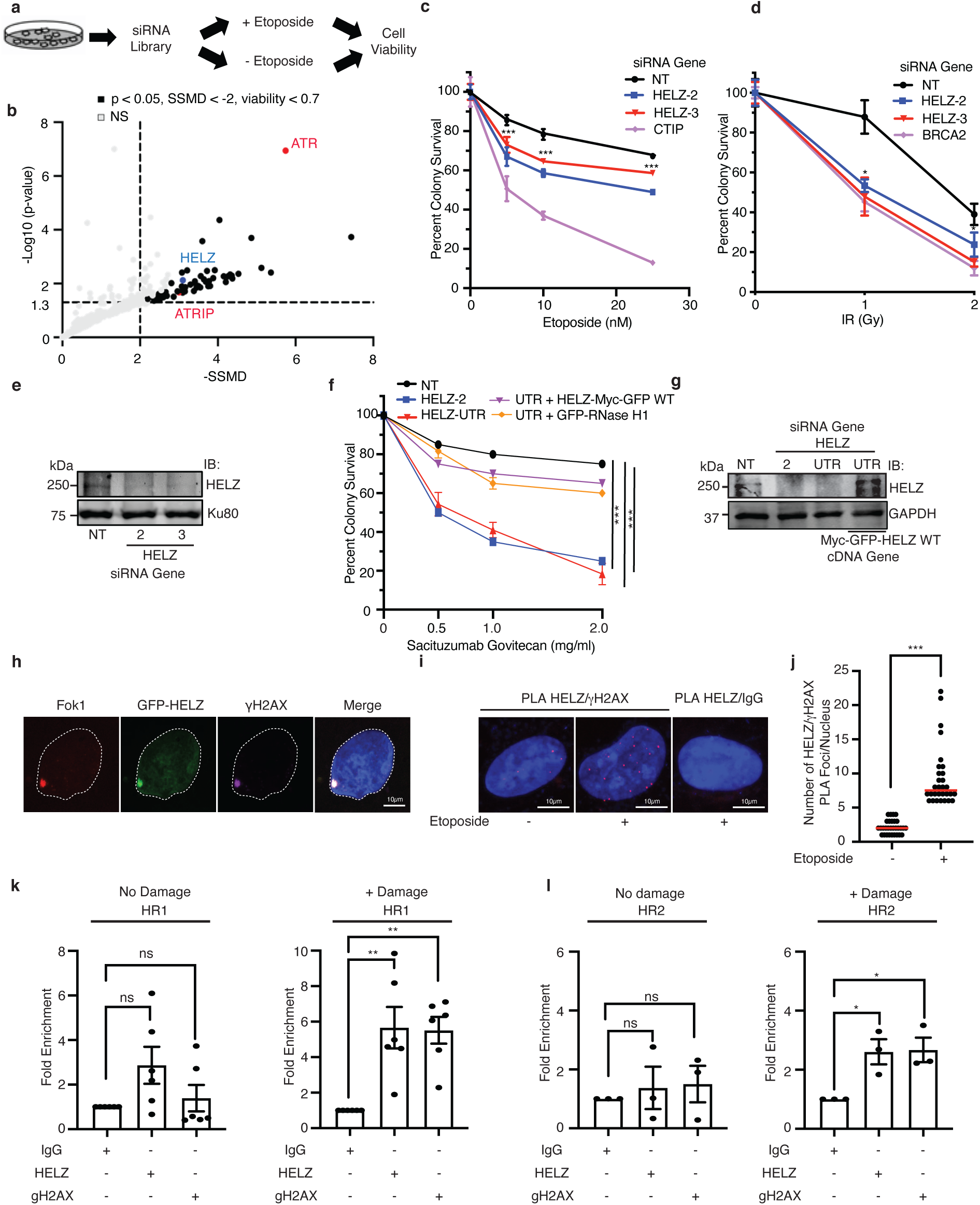
HELZ depletion causes hypersensitivity to DSB-inducing agents in a R loop dependent manner and HELZ localizes and binds to DSBs. **a** Primary etoposide hypersensitivity siRNA screen flow diagram. H128 cells were transfected with a siRNA library targeting 1006 annotated nuclear enzymes. The screen was conducted in triplicate in 96-well plates using ATR and ATRIP siRNA as positive controls and a non-targeting (NT) siRNA as a negative control. 48 hours (h) post-transfection, cells were treated with or without 10 µM etoposide for 72 h prior to measuring cell viability with rezasurin reagent. **b** All sensitized genes with normalized cell viability below 1 are displayed in a plot of −log10 (p-value) against the -strictly standardized mean difference (SSMD). Significant etoposide sensitization hits were identified using the following criteria: an average cell viability < 0.7, an average SSMD < −2, and a two-tailed t-test p-value < 0.05. **c-d** HELZ depletion causes hypersensitivity to etoposide (**c**) and IR (**d**). U2OS cells were transfected with indicated siRNA. 96 hours (h) after transfection, cells were treated with or without indicated doses of etoposide continuously or IR. Mean and standard deviation (SD) from three independent replicas is shown. * p < 0.05, *** p <0.001. **e** Western blot analysis showing HELZ knockdown in U2OS cells from (**c-d**). **f** MDA-MB-231 cells were transfected with indicated siRNA with or without HELZ-Myc-GFP or RNase H1. 72 h after transfection, cells were treated with or without indicated doses of sacituzmab govitecan (SG) continuously. Mean and standard error of the mean (SEM) from three independent replicas is shown. **g** Western blot analysis showing HELZ knockdown and rescue in MDA-MB-231 cells from (**f**). **h** U2OS-265 Fok1 cells were transfected with HELZ-Myc-GFP, and induced for Fok1 expression with 1 µM Shield-1 and 1 µM 4-OHT for 4 h. Samples were processed for direct immunofluorescence with mCherry-Fok1 and HELZ-Myc-GFP and indirect immunofluorescence with anti-γH2AX antibody. DNA was visualized by DAPI staining. **i** Representative images and **j** quantification from two independent replicas of PLA foci between HELZ and γH2AX before and after treatment with 20 µM etoposide for 4 h. The median is indicated by a horizontal line. *** p < 0.001. **k-l** CHIP-qPCR analysis of endogenous HELZ enrichment at HR1 (**k**) and HR2 (**l**) prone DNA repair sites in DIvA cells. DSBs were induced by treatment with 500 nM 4-OHT for 4 h. For **k**-**l,** mean and SEM from three independent replicas is shown. * indicates p < 0.05, ** p<0.01, *** p < 0.001, ns indicates not significant.

### HELZ depletion causes hypersensitivity to DSB-inducing agents in a R loop dependent manner

We found that HELZ depletion by multiple siRNA causes etoposide hypersensitivity in H128 cells as well as additional cell types, including U2OS osteosarcoma cells (Fig. 1b-c and Supplementary Fig. S4a), suggesting that the phenotype is not cell type specific and may be a general mechanism governing etoposide resistance. We generated a custom rabbit polyclonal anti-HELZ antibody and confirmed HELZ knockdown in these cells (Fig. 1e and Supplementary Fig. S4c). This antibody also recognizes overexpressed HELZ-Myc-GFP (Fig. 1g) thus confirming specificity of the antibody for both endogenous and overexpressed HELZ. HELZ depletion also caused hypersensitivity to additional DSB-inducing agents ionizing radiation (IR) and camptothecin (CPT), a topoisomerase I inhibitor, in these cells (Fig. 1d and Supplementary Fig.4b) and sacituzumab govitecan (SG), an antibody drug conjugate (ADC) consisting of the active irinotecan metabolite SN-38 conjugated to anti-Trop-2 monoclonal antibody, in MDA-MB-231 and BT549 triple-negative breast cancer (TNBC) cells (Fig. 1f-g and Supplementary Fig. S4d-e). Furthermore, the SG hypersensitivity of HELZ depletion was alleviated by expression of HELZ-Myc-GFP WT and GFP-RNase H1, which catalyzes the cleavage of RNA in RNA-DNA hybrids (Fig. 1f-g and Supplementary Fig. S4d-e), suggesting that HELZ responds generally to DSBs, the phenotype is not due to an off-target effect, and that HELZ governs resistance of cells to DSB-inducing agents in a R loop dependent manner.

### HELZ localizes and binds to DSBs

To determine if HELZ localizes to DSBs, we induced DSBs using mCherry-LacI-Fok1 endonuclease in U2OS reporter cells integrated with lac operator repeats^49^. HELZ-Myc-GFP co-localized with Fok1-induced DSBs that were also marked by γH2AX (Fig. 1h), suggesting that HELZ localizes to DSBs. A proximity ligation assay (PLA) validated the proximity of HELZ with γH2AX following etoposide treatment (Fig. 1i-j). To determine if HELZ binds to endogenous DSBs, we performed chromatin IP (ChIP) with quantitative PCR (qPCR) using U2OS DIvA (DSB Inducible via AsiSI) cells, in which 4-hydroxytamoxifen (4-OHT) treatment triggers nuclear localization of AsiSI to induce approximately 150 sequence-specific DSBs, dispersed across the genome^50^. HELZ showed enriched accumulation to HR1, HR2, and NHEJ prone DSB repair sites after 4-OHT treatment (Fig. 1k-l and Supplementary Fig. S4f), suggesting that HELZ binds to DSBs. We found a mild increase in chromatin localization but no evidence for a significant change in HELZ protein levels by biochemical fractionation following etoposide treatment (Supplementary Fig. S4g-i).

### HELZ binds and unwinds RNA-DNA hybrids and prevents the accumulation of R loops genome-wide and at DSBs

Several members of the DEAD box family of RNA helicases are involved in R loop homeostasis^51^. Given that HELZ contains a RNA helicase motif (Supplementary Fig. S5a-c), we determined whether HELZ depletion leads to an increase in R loops as determined by immunofluorescence (IF) microscopy using the S9.6 monoclonal antibody. An increase in S9.6 signal intensity was observed with HELZ depletion compared with a NT control in U2OS and MDA-MB-231 cells, which was alleviated by RNaseH treatment (Fig. 2a-b and Supplementary Fig S5d-f), demonstrating specificity of HELZ in preventing accumulation of endogenous R loops. We validated that HELZ depletion or etoposide treatment leads to an increase in S9.6 signal in HCT-116 colorectal cells by slot blot analysis, although no further increase in S9.6 signal was observed with combined HELZ depletion and etoposide treatment (Fig. 2c and Supplementary Fig. S5g), suggesting that HELZ may play a role in preventing etoposide-induced R loops. Overexpression of HELZ-Myc-GFP wild-type (WT) but not K674N, a mutation in the Walker A motif, which abolishes ATPase activity in DEAD box helicases^52^ (Supplementary Fig. S5a-c), alleviated the induction of S9.6 signal intensity of HELZ depletion (Fig. 2c) suggesting that HELZ prevents accumulation of R loops in cells through its ATPase activity.

**Figure 2.**
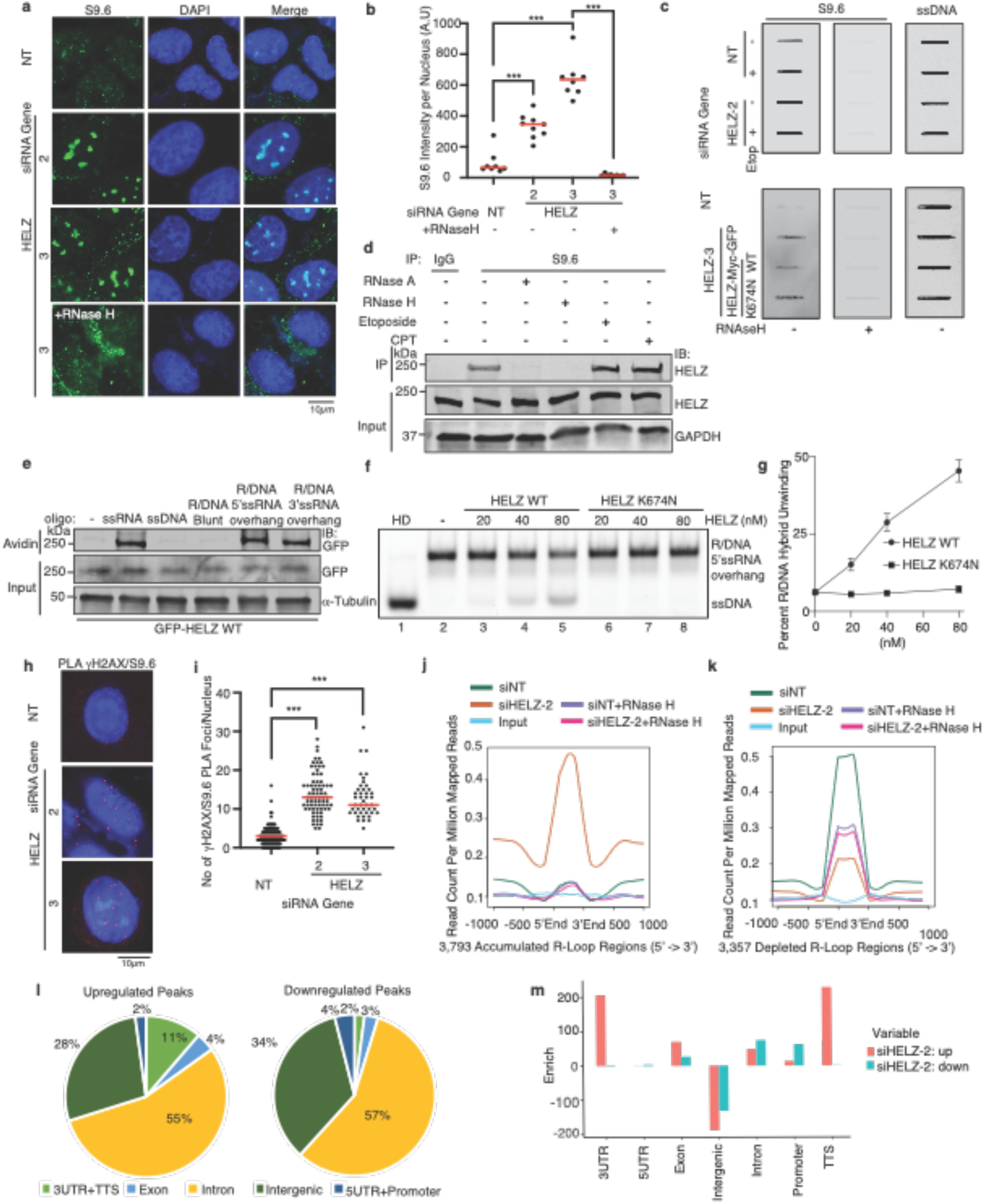
HELZ binds and unwinds RNA-DNA hybrids and prevents R loop accumulation genome-wide and at DSBs. **a-b** U2OS cells were transfected with HELZ siRNA and processed 72 h later for indirect immunofluorescence with anti-S9.6 antibody. Representative images and quantification are shown. The median is indicated by a horizontal line. *** p < 0.001. **c** Slot blot analysis of DNA-RNA hybrids from HCT116 cells transfected with indicated siRNA and cDNA with or without etoposide treatment. Equal amounts of DNA treated or untreated with RNase H (+) or denatured (ssDNA), were loaded as indicated. The membrane was immunoblotted with anti-S9.6 antibody or antibody recognizing ssDNA to verify equivalent DNA loading. **d** DNA-RNA hybrid immunoprecipitation western blot (DRIP-WB) analysis of lysate from U2OS cells demonstrating pull-down of endogenous HELZ with and without treatment with etoposiode, CPT, RNase H, and RNase A for 30 min. **e** Streptavidin pulldown of biotin-labeled nucleic acids incubated with HELZ-Myc-GFP WT purified from HEK293T cells. **f-g** Insect purified recombinant HELZ WT and K674N were tested in RNA/DNA hybrid unwinding assay with 5’ ssRNA overhangs. Representative blot (**f**) and quantification of the unwinding experiments showing the percentage of unwound ssDNA relative to the RNA/DNA hybrid substrate with mean and SD from three independent replicas is shown (**g**). **h-i** PLA foci suggesting an interaction between γH2AX and S9.6 in U2OS cells 72 h following HELZ knockdown. Representative images (**h**) and quantification (**i**) are shown. The median is indicated by a horizontal line. *** p < 0.001. **j** Ngsplot of R loop read counts across the accumulated R loop regions. **k** Ngsplot of R loop read counts across the depleted R loop regions. **l** Genomic distribution of significantly upregulated and downregulated R loop regions. **m** Enrichment analysis of the significantly upregulated R loops (red) and the significantly downregulated R loops (blue) in different genomic features. Comparison of differential R loop peak numbers in various gene features with expected R loop peak numbers based on the percentage of the length of each chromatin feature is shown as log2 Observed/Expected (log2 Obs/Exp).

To determine if HELZ interacts with R loops in cells, we performed DNA-RNA immunoprecipitation (DRIP) analysis using the S9.6 antibody, which showed an interaction of HELZ with R loops that was increased after treatment of cells with etoposide or CPT and abolished with RNase H or RNase A treatment (Fig. 2d), suggesting that HELZ binds to R loops and that this interaction increases in response to DSBs. To determine if HELZ binds to RNA-DNA hybrids *in vitro*, we performed streptavidin pulldown of biotin labeled RNA-DNA hybrid structures incubated with HELZ-Myc-GFP expressed in HEK293T cells. HELZ interacted with ssRNA and RNA-DNA hybrid structures with 5’ or 3’ ssRNA overhangs but not ssDNA or pure hybrids (Fig. 2e), indicating specificity of HELZ in binding to ssRNA and RNA-DNA hybrids with ssRNA overhangs. We confirmed comparable binding of insect purified recombinant HELZ WT and K674N to RNA-DNA hybrids with 5’ ssRNA overhangs to a similar extent but not to ssDNA with electrophoretic mobility shifty assay (EMSA) (Supplementary Fig. S5h-l). To determine if HELZ unwinds RNA-DNA hybrids, insect purified recombinant HELZ WT and K674N was incubated with RNA-DNA hybrids with 5’ ssRNA overhangs, and HELZ WT but not K674N showed a dose dependent unwinding of these structures (Fig. 2f-g), suggesting that HELZ has *in vitro* unwinding activity towards RNA-DNA hybrids with 5’ ssRNA overhangs and that ATPase dead K674N is also dead for RNA-DNA unwinding.

Consistent with our observation that HELZ binds to R loops in response to DSBs, both HELZ and S9.6 showed an enriched accumulation to HR1 prone DSBs in DIvA cells after induction of DSBs (Supplementary Fig. S5m). To determine if HELZ prevents accumulation of R loops at DSBs, we performed PLA with γH2AX and S9.6 following HELZ depletion. A significant increase in γH2AX and S9.6 PLA foci following HELZ depletion was observed (Fig. 2h-i), suggesting that HELZ prevents the accumulation of R loops at DSBs.

### HELZ depletion causes genome-wide accumulation of R loops

To determine the global impact of HELZ deficiency on R loops, we performed DRIP sequencing (DRIP-seq) analysis following IP of RNA-DNA hybrids with the S9.6 antibody in U2OS cells silenced for HELZ. DRIP-seq profiling revealed 84,618 R loop peaks that overlapped with 14,121 annotated genes (RefSeq, hg38) in siNT cells and 91,172 R loop peaks that overlapped with 14,231 annotated genes in siHELZ cells (Supplementary Fig. S6a-b). Most of the R loop-positive genes (12,845) were common to both cells (Supplementary Fig. S6b). Importantly, we validated that R loop enrichment at a subset of genes *APOE*, *RPL13A*, *EGR*1, and *HMGA*1 resulting from HELZ knockdown was not due to an increase in mRNA levels as detected by qRT-PCR at loci that have been previously validated for this purpose^53,54^ (Supplementary Fig. S6c), suggesting that R loop accumulation resulting from HELZ knockdown is not due to HELZ’s role in protein translation and mRNA stability^31,32^.

We plotted R loop read counts across all R loop peaks identified in both cells and found that R loops were upregulated upon HELZ depletion and was decreased in cells treated with RNase H (Supplementary Fig. S6d p = 1.36×10^−7^). A substantial portion of R loops were located in gene body regions (71%, including exons and introns) which may suggest an active transcription of the genes (Supplementary Fig. S6a). We then plotted R loop read counts across defined human RefSeq (hg38) genes and found that R loops in both cells were enriched across the genes compared to RNase H-treated samples (Supplementary Fig. S6e). HELZ depletion showed a global R loop accumulation in the genomic regions (Supplementary Fig. S6e, p = 1.97×10^−10^). Using DESeq2, we identified 3,793 regions that accumulated R loops significantly upon HELZ knockdown, and 3,357 regions that showed depleted R loops (Supplementary Fig. S6f, false discovery rate (FDR) < 0.05, log_2_(Fold change) > 1). The R loop read count plotting confirmed an increase of R loops across the accumulated R loop regions (Fig. 2j, p = 1.70×10^−18^) and a decrease of R loops across the depleted R loop regions (Fig. 2k, p = 7.91×10^−22^). Differential R loops were enriched particularly in 3’UTR and transcription termination sites (TTS), compared to expected values (Fig. 2l-m).

### HELZ depletion causes genomic instability via R loop accumulation

To determine if HELZ depletion leads to genomic instability, we examined for accumulation of spontaneous γH2AX. HELZ depletion led to an increase in γH2AX in U2OS, HeLa cervical cancer cells, and RPE-1 nontumorigenic hTERT immortalized retinal pigment epithelial cells (Supplementary Fig. S7a-e). HELZ depletion also led to an increased percentage of U2OS cells with micronuclei compared with a NT control (Supplementary Fig. S7f-g). Moreover, HELZ depletion or knockout (KO) by CRISPR/Cas9 in U2OS cells and HeLa cells led to an increase in 53BP1 nuclear bodies (Fig. 3a-b and Supplementary Fig. S7h-n) which was rescued by overexpression of mCherry-RNase H1 (Fig. 3a-b), suggesting that HELZ depletion causes genomic instability via R loop accumulation.

**Figure 3.**
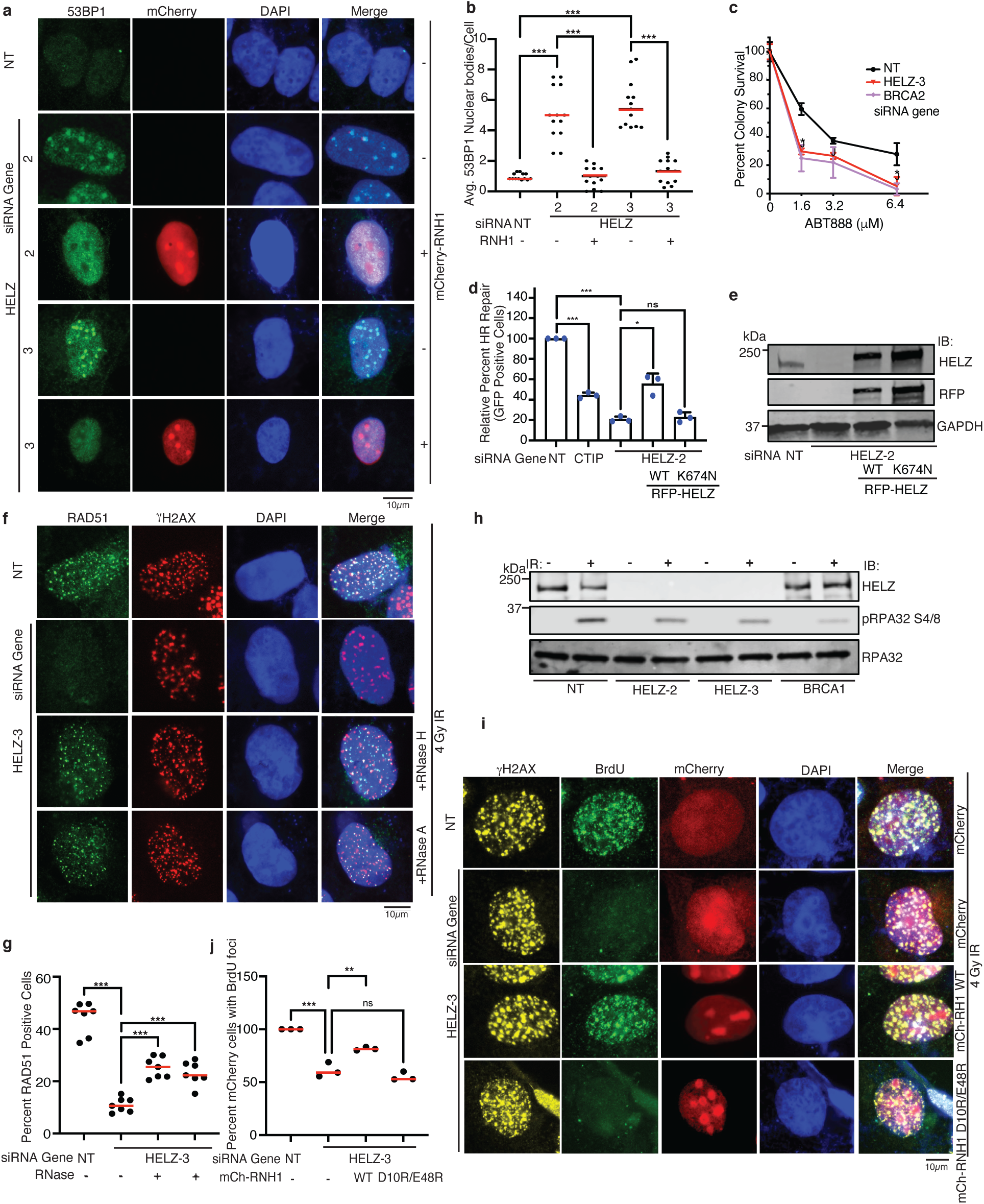
HELZ promotes homologous recombination and resection by preventing the accumulation of R loops and prevents R loop-mediated genomic instability. **a-b** Representative images and quantification for 53BP1 nuclear bodies in U2OS cells silenced for HELZ or a NT control and rescued for spontaneous 53BP1 nuclear bodies with overexpression of mCherry-RNaseH1. The median is indicated by a horizontal line. *** p < 0.001. **c** U2OS cells transfected with indicated siRNA were seeded for colony formation. 72 h post transfection, cells were treated with indicated doses of ABT888 every 2-3 days. Percent survival colonies is shown. Mean and SD from three independent replicas is shown. * p < 0.05. **d** HEK293 cells containing an integrated DR-GFP HR reporter were silenced with indicated siRNA, and transfected with I-SceI endonuclease and RFP (-) or RFP-HELZ (WT, K674N) as shown. Live cells collected were subjected to flow cytometry. RFP-expressing cell population was sorted and percent of GFP positive cells within this population was determined to analyze HR efficiency. Mean and SD from three independent replicas is shown. * p < 0.05, *** p < 0.001. **e** Western blot analysis showing HELZ knockdown and expression of RFP-HELZ WT and K674N from **d**. **f-g** U2OS cells were depleted for HELZ for 72h, treated with 4 Gy IR with or without RNase H or RNase A treatment in live cells for 1 h before fixation, and processed 4h after IR for indirect immunofluorescence with indicated antibodies. Representative images and quantitation are shown. The median is indicated by a horizontal line. *** p < 0.001. **h** 293T cells were transfected with NT, HELZ, or BRCA1 siRNA and 72h later, treated with 4 Gy IR and processed for western blot analysis with indicated antibodies. **i-j**. U2OS cells were transfected with HELZ or NT siRNA for 72h and mCherry-RNaseH1 WT or D10R/E48R for 48 h, treated with 4 Gy IR, and processed 4h later for indirect immunofluorescence with indicated antibodies. Representative images and quantitation are shown. The median is indicated by a horizontal line. *** p < 0.001.

### HELZ promotes homologous recombination and resection by preventing accumulation of R loops at DSBs

PARP inhibitor sensitivity is associated with impairment in HR^55,56^. Indeed, similar to BRCA2, HELZ depletion in U2OS cells caused hypersensitivity to ABT888 (Veliparib), a PARP inhibitor (Fig. 3c). To determine directly if HELZ functions in HR, we examined HELZ depletion in 293T and U2OS cells integrated with the direct repeat (DR)-GFP HR reporter substrate in which expression of the rare cutting endonuclease I-SceI generates a DSB which when repaired by HR restores GFP expression^57^. Similar to CtIP, HELZ depletion impaired HR (Fig. 3d-e and Supplementary Fig. S8a-b), which was alleviated by expression of RFP-HELZ WT but not K674N (Fig. 3d-e), suggesting that HELZ helicase activity promotes HR. Using the EJ7-GFP classical NHEJ (c-NHEJ) reporter integrated in HEK293 cells, we also observed a mild decrease in c-NHEJ with HELZ depletion (Supplementary Fig. S8c), suggesting that HELZ may also promote c-NHEJ. We observed no significant decrease in cells in S and G2 phase following HELZ depletion, suggesting that the impairment in HR is not an indirect effect of cell cycle change (Supplementary Fig. 8d). Consistent with our reporter data, HELZ depletion in U2OS cells caused an impairment in IR-induced RAD51 and 53BP1 foci (Fig. 3f-g and Supplementary Fig. S8e-h). Interestingly, the impairment in IR-induced RAD51 foci resulting from HELZ depletion was alleviated by treatment of cells with RNAse H and RNase A or overexpression of mCherry-RNase H1 (Fig. 3f-g and Supplementary Fig. S8e-f), suggesting that HELZ promotes HR by preventing accumulation of R loops at DSBs. DSB repair by HR is initiated by DNA end resection. We examined for IR-induced RPA70 foci, a marker for resection and found that similar to BRCA1, HELZ depletion in U2OS cells caused an impairment in IR-induced RPA70 foci (Supplementary Fig. S8i-j). Consistent with these findings, HELZ depletion also caused an impairment in IR-induced phosphorylation of RPA at Ser4/8 (Fig. 3h). To more directly determine if HELZ promotes DNA end resection, we labeled U2OS cells with BrdU, treated the cells with IR, and examined BrdU exposure under nondenaturing conditions, which labels ssDNA. HELZ depletion impaired BrdU foci, which was alleviated by expression of mCherry-RNaseH1 WT but not catalytically dead D10R/E48R^58^ or mCherry alone (Fig 3i-j and Supplementary Fig. S8k), suggesting directly that HELZ promotes DNA end resection by preventing accumulation of R loops at DSBs.

### HELZ interacts with BRCA1 in damage regulated and RNA-dependent manner

Given that BRCA1 promotes DNA end resection^13–18^ and HR^59^ and functions with several other DEAD box family members in resolving R loops and/or facilitating HR^59^, we examined the interaction of HELZ with BRCA1. First, we found that both endogenous HELZ and BRCA1 showed enriched accumulation to HR1 prone DSBs after 4-OHT treatment in DIvA cells (Supplementary Fig. 9a). Similarly, both HELZ-Myc-GFP and endogenous BRCA1 co-localized at Fok1 induced DSBs in U20S-265 cells (Fig. 4a). A PLA assay confirmed the close proximity of endogenous HELZ and BRCA1, which increased after etoposide treatment (Fig. 4b-c and Supplementary Fig. S9b-c). To determine if HELZ and BRCA1 physically interact, we performed co-immunoprecipitation (co-IP) studies. Co-IP of HELZ-Myc-GFP expressed in HEK293T cells pulled down endogenous BRCA1 (Supplementary Fig. S9d) and similarly co-IP of GFP-BRCA1 expressed in HEK293T cells pulled down endogenous HELZ (Supplementary Fig. S9e), suggesting that HELZ and BRCA1 interact in a complex. The co-IP of HELZ-Myc-GFP and endogenous BRCA1 was increased following treatment of cells with IR or etoposide and was preserved following benzonase nuclease or RNase H treatment (Fig. 4d-e), implying that the interaction of HELZ and BRCA1 is not nucleic acid mediated.

**Figure 4.**
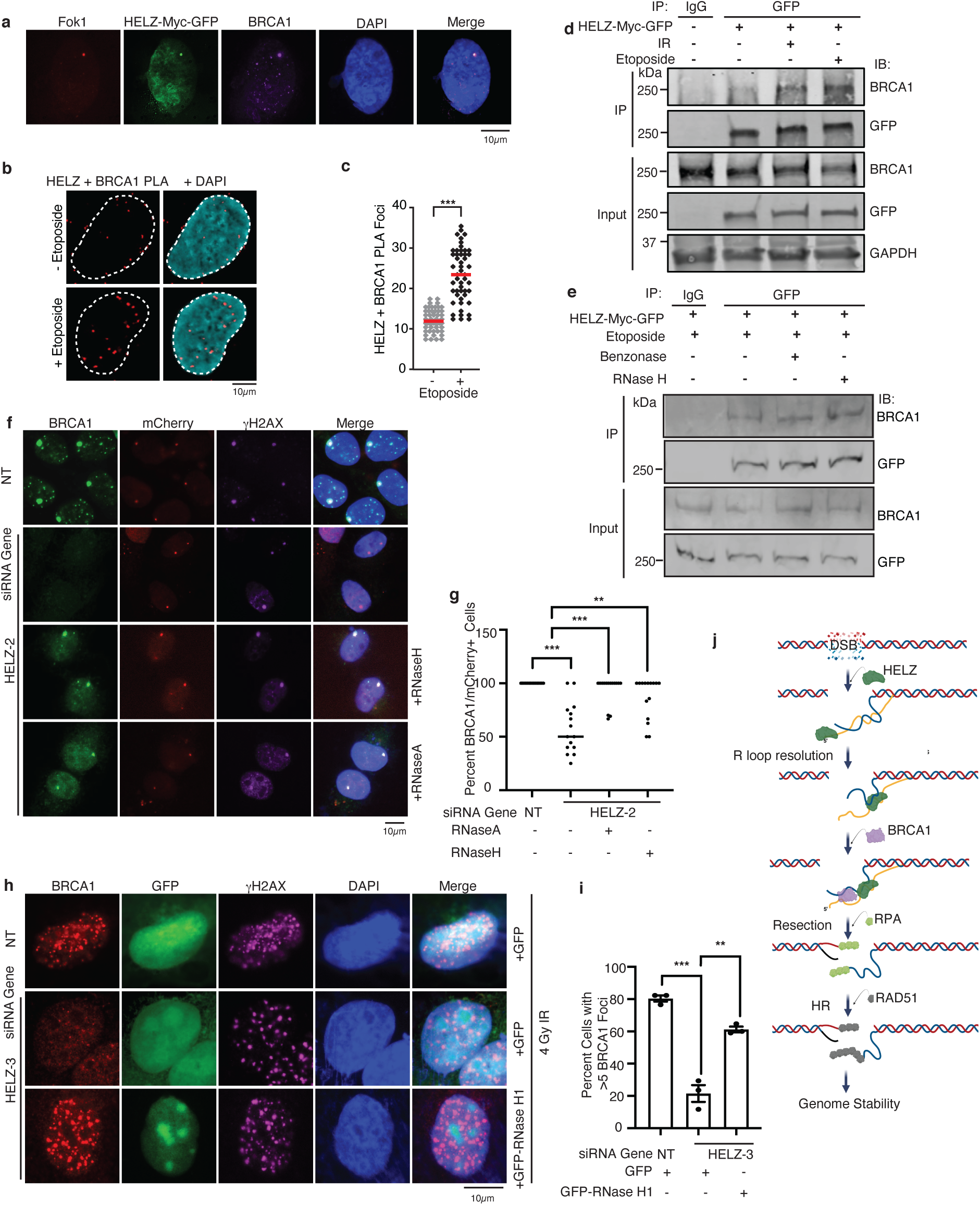
HELZ interacts with BRCA1 in a damage regulated and RNA-dependent manner and facilitates BRCA1 recruitment to DSBs by preventing accumulation of R loops. **a** U2OS-265 Fok1 cells were induced for Fok1 expression with 1 µM Shield-1 and 1 µM 4-OHT for 4 h. Localization of Fok1, HELZ-Myc-GFP and BRCA1 were visualized by direct immunofluorescence with mCherry and GFP and indirect immunofluorescence with anti-BRCA1 antibodies and stained with DAPI. **b-c** PLA foci suggestive of an interaction between endogenous HELZ and BRCA1 in U2OS cells before and after etoposide treatment (20 µM 4h). Representative images and quantification are shown. The median is indicated by a horizontal line. *** p < 0.001. **d** HEK293T cells were transfected with HELZ-Myc-GFP, treated with 10 Gy IR or 20 µM etoposide, harvested 4h later, IP’ed with anti-GFP antibodies, run on SDS-PAGE, and immunoblotted with indicated antibodies. **e** HEK293T cells were transfected with HELZ-Myc-GFP, treated with 20 µM etoposide and harvested after 4h. Benzonase nuclease or RNase H treatment was done on the same lysate for 30 min in 37°C water bath followed by 10 min centrifugation to remove digested debris. Samples were run on SDS-PAGE and blotted with indicated antibodies. **f-g** U2OS-265 Fok1 cells were depleted for HELZ. After 72 h, DSBs were induced by Shield-1 and 4-OHT. Cells were fixed after 4 h and stained with indicated antibodies. RNase H or RNase A treatment were done in live cells for 1 h before fixation. Percentage of cells with both a single red focus and green focus of BRCA1 were quantified. The median is indicated by a horizontal line. ** p < 0.01, *** p < 0.001. **h-i** U2OS control cells and U2OS HELZ KO cells were treated with 4 Gy IR, fixed after 4 h, and processed for indirect immunofluorescence with indicated antibodies. Representative images (**h**) and quantification (**i**) are shown. Mean and SD from three independent replicas is shown. ** p < 0.01. **j** XX Proposed model of HELZ activity. Regardless of DNA damage, HELZ prevents R loop accumulation to maintain genomic stability. It also interacts with BRCA1. DNA-RNA hybrid formation may be enhanced in the vicinity of DSBs due to the DNA rotation freedom provided by the break. In the absence of HELZ, R loop removal at DNA DSBs is hampered which results in impaired BRCA1 recruitment to the break site. HR cannot be completed successfully and that leads to genomic instability.

### HELZ facilitates BRCA1 recruitment to DSBs by preventing accumulation of R loops

To determine if HELZ functions upstream of BRCA1, we examined for the localization of BRCA1 to DSBs following HELZ depletion. HELZ depletion impaired BRCA1 localization to Fok1-induced DSBs in U2OS-265 cells (Fig. 4f-g Supplementary Fig. S9f-g) and to IR-induced DSBs (Fig.4h-i and Supplementary Fig. S9h-i). Given that HELZ depletion impairs HR in a RNA-mediated manner, we hypothesized that BRCA1 localization may be influenced by the accumulation of R loops. Indeed, treatment of U2OS-265 cells silenced for HELZ with RNase H or RNase A or GFP-RNase H1 overexpression in U2OS cells alleviated the impairment in BRCA1 localization to DSBs from HELZ depletion (Fig. 4f-i), suggesting that HELZ promotes BRCA1 recruitment to DSBs by preventing accumulation of R loops. Consistent with a role for HELZ in promoting HR by facilitating BRCA1 recruitment, 53BP1 depletion in U2OS cells rescued the impairment in IR-induced RAD51 foci from HELZ depletion (Supplementary Fig. S10a-d).

## Discussion

Our findings define HELZ as a unique RNA-DNA helicase that resolves R loops to facilitate BRCA1 recruitment to DSBs and thereby promote DNA end resection and HR. These findings identify HELZ as a critical regulator of R loop homeostasis by resolving R loops, establish HELZ as an important factor in promoting DNA end resection and DSB repair by HR, and provide a mechanistic link between the removal of R loops and promotion of BRCA1 recruitment to facilitate DNA end resection and HR via HELZ to promote genome integrity and govern resistance of cancer cells to sacituzumab govitecan and other DSB-inducing agents. In this regard, we completed a synthetic lethal etoposide resistance siRNA screen and found that HELZ depletion causes hypersensitivity to DSB-inducing agents etoposide, IR, PARP inhibitor, and sacituzumab govitecan in a R loop-dependent manner, and HELZ localizes and binds to DSBs. HELZ depletion further leads to the spontaneous accumulation of micronuclei, γH2AX, and 53BP1 nuclear bodies, which can be rescued by RNase H1 expression, suggesting that HELZ prevents genomic instability in a R loop-dependent manner. Indeed, we found that HELZ binds to R loops in response to etoposide and CPT treatment and prevents their accumulation in a helicase activity-dependent manner, genome-wide with enrichment at TTS and at DSBs, and in response to DSBs. Biochemically, HELZ preferentially binds to ssRNA and RNA-DNA hybrids with ssRNA overhangs and unwinds RNA-DNA hybrids with 5’ssRNA overhangs. HELZ helicase activity furthermore promotes HR and DNA end resection, and this impairment following HELZ depletion is alleviated by RNase H treatment and/or RNase H1 expression, suggesting that HELZ promotes HR and DNA end resection by preventing the accumulation of toxic R loops at DSBs. Interestingly, HELZ complexes with BRCA1 in a damage-regulated manner and facilitates its recruitment to DSBs by preventing R loop accumulation. Thus, our findings are consistent with a model in which in response to DSBs, HELZ localizes, binds, and resolves unscheduled harmful R loops at DSBs to facilitate BRCA1 recruitment to promote DNA end resection and HR (Fig. 4j) Dysregulation of this pathway leads to unscheduled R loop accumulation, which impairs HR leading to genomic instability. Given that HELZ depletion leads to accumulation of R loops at TTS and DSBs and that HELZ unwinds RNA-DNA hybrids with 5’ssRNA overhangs, our data suggest that HELZ may act at DSBs at transcribed regions where the nascent ssRNA anneals to DNA behind RNA polymerase II although our model does not exclude other possibilities such as HELZ resolving R loops at DSBs formed independent of transcription or co-transcriptionally independent of DSBs.

HELZ has previously been reported to have a helicase-independent role in promoting cell proliferation, translation initiation, and ribosomal protein 6 phosphorylation^32^ and regulating the stability and translation of mRNA via its interaction with mRNA decay factors in the cytoplasm^31^. We now show that HELZ has biochemical activity as a ssRNA-specific RNA-DNA helicase that is critical for promoting genome stability by resolving R loops to facilitate DSB repair by HR. Given that HELZ WT but not catalytically inactive K674N rescues the accumulation of R loops resulting from HELZ deficiency, HELZ’s role in resolving R loops and/or preventing accumulation of R loops is likely mediated through its helicase activity and not as a result of increased mRNA levels from its noncatalytic role in mRNA stability. Indeed, we found no evidence for an increase in mRNA levels at a set of genes with increased R loops resulting from HELZ depletion that has been validated for this purpose^53,54^. Furthermore, given that cellular hypersensitivity to sacituzumab govitecan, genomic instability, and impairment in HR, end resection, and BRCA1 recruitment resulting from HELZ depletion can all be alleviated by RNase treatment or RNase H1 overexpression, our findings suggest the HELZ’s role in promoting genome stability is mediated through its activity in resolving R loops.

It has been reported that the accumulation of R loops around DSBs competes with RAD51 localization and binding to DSBs^60^. Consistent with these data, we found that the impairment in IR-induced RAD51 foci from HELZ depletion was alleviated by RNase H and RNase A treatment, suggesting a role for R loops in impairing HR and that HELZ resolves R loops to facilitate HR. In addition, we found that impairment in DNA end resection resulting from HELZ depletion is rescued with RNase H1 WT but not catalytically inactive D10E/E48R, suggesting that HELZ also promotes DNA resection by resolving R loops. These data are consistent with a previous report that R loops can impair DNA end resection^23^. While R loops have also been reported to promote HR and DNA end resection^22^, we speculate that excessive amounts of unscheduled R loops may be harmful, which are resolved at least in part by HELZ, or that this may be due to spatial differences of R loops at different sites or temporal differences whereby R loops are first required to initiate repair and then must be removed to complete repair. Our data suggest a unique mechanism for promotion of DNA end resection and HR whereby HELZ facilitates the recruitment of BRCA1 to DSBs via resolution of toxic R loops. A similar impairment in BRCA1 recruitment to DSBs by R loop accumulation has been observed with depletion of the DIS3 ribonuclease^61^. It is possible that HELZ may unwind R loops, which are subsequently degraded by DIS3. Although our data and those of others suggest that BRCA1 also binds to RNA-DNA hybrids ^20,62,63^ and BRCA1 has previously been suggested to function with Senataxin, DHX9, and COBRA1 in promoting R loop resolution and/or preventing R loop accumulation^62,64,65^, we speculate that this may be context dependent perhaps based on levels or types of RNA-DNA hybrids.

A number of helicases, including Senataxin^66–68^, AQR^35^, UPF1^69^, PIF1^70^, WRN^71^, BLM^72^, RTEL1^73^, FANCM^74–76^; POLQ^77^, ATRX^78^, SMARCAL1^74^, ZRANB3^74^, DDX1^21,79,80^, DDX5^81^, DDX17^82^, DDX18^83^, DDX19^84^, DDX21^85,86^, DDX23, UAP56/DDX39B^53^, DDX41^87,88^, DDX47^89^ and DHX9^90–93^ have been reported to be involved in R loop homeostasis with Senataxin^26,94^, AQR^34^, BLM^95^, DHX9^64,91^, DDX1^21,79^, DDX5^29^, DDX17^82^, DDX18^83^, and UPF1^69^ involved in facilitating the repair of DSBs. How HELZ may function with these helicases in resolving R loop homeostasis to facilitate DSB repair is unclear. It is possible that HELZ may act on specific types of R loop structures or sites or act in a temporal manner potentially in collaboration with other helicases. In this regard, similar to Senataxin, HELZ unwinds RNA-DNA hybrids with 5’ ssRNA overhangs but not RNA-DNA hybrids with no overhangs or 3’ ssRNA overhangs (see accompanying manuscript); however, HELZ is unique in binding specifically to ssRNA but not ssDNA and unwinds only RNA-DNA hybrids in contrast to Senataxin which binds to ssDNA with higher affinity than ssRNA and unwinds both RNA-DNA and DNA-DNA hybrids^96^. BLM, DDX5, and DDX21 unwind RNA-DNA hybrids with no overhangs^72,81,86^, DHX9 and DDX17 unwind RNA-DNA with 3’ overhangs^90,97^, and DDX39B unwinds RNA-DNA hybrids with 5’, 3’ or no overhangs^53^. The unique ssRNA-specific RNA-DNA helicase activity of HELZ may help explain, at least in part, its critical role in the resolution of toxic R loops to promote BRCA1 recruitment to DSBs to facilitate DNA end resection and HR.

Given that HELZ depletion sensitizes cancer cells to sacituzumab govitecan, PARP inhibitor, and other DSB-inducing agents, which can be rescued by RNase H1, our findings suggest that HELZ-mediated resolution of R loops governs resistance of cancer cells to DSB-inducing agents and provides rationale for targeting the helicase activity of HELZ as an adjunct to DSB-inducing agents or using HELZ expression as a biomarker for selecting patients who might best respond to these agents. This may be particularly useful for triple-negative breast cancer (TNBC) patients treated with sacituzumab govitecan where despite gaining FDA approval and improving overall survival (OS)^98–103^, overall response rates are modest at 33% and no reliable biomarkers exist, including Trop-2 expression or *BRCA1/2* status^104^.

## Methods

### Cell lines

Human SCLC line NCI-H128 was provide by the laboratory of Dr. Taofeek Owonikoko and grown in RPMI 1640 (Gibco) with 7.5% FBS. HEK293T, HCT116, HeLa, U20S, and MDA-MB-231 mammalian cell lines were purchased from American Type Culture Collection (ATCC, Manassas, VA). U20S-235 mCherry-LacI-Fok1 cell line were provided by Dr. Roger Greenberg. and U20S-DR-GFP, HEK293-DR-GFP and HEK293-EJ7-GFP cell lines were obtained from Dr. Jeremy Stark. AsiSI-ER-U2OS (DIvA-DSB inducible via AsiSI) cells were provided by Dr. Gaëlle Legube. All cell lines were grown in DMEM (Gibco) antibiotic-free media supplemented with 7.5% FBS. U20S-235 was additionally cultured in 2 μg/mL puromycin and 100 μg/mL hygromycin. DIvA cells were additionally cultured in 1 μg/mL puromycin. All cell lines were grown at 37 °C under humidified conditions with 5% CO2 and 95% air.

### Primary etoposide siRNA screen

A custom siRNA library targeted 1,006 annotated nuclear enzymes in a 96-well plate SMARTpool format, with each well containing 4 siRNAs targeting unique sequences within the same gene (Dharmacon). Each plate had 9 negative control siNT wells, three positive sensitivity control siATR wells, and one blank well. We included siCHK1, siATRIP, mock transfected (no siRNA), and non-transfected (no transfection reagent) control wells on the final plate. Each well in the 96-well plates received 12,000 H128 cells and 0.3 µL of Lipofectamine RNAiMAX (Invitrogen), as well as a final siRNA concentration of 25 nM and a final volume of 100µL. After 24 hours, each transfection plate was split into 4 clear-bottomed plates with final well volumes of 100 µL. After another 24 hr, 50 µL of media was added to the two non-treated plates while 50 µL of etoposide-containing media was added to one plate each with a final concentration of 10 µM. After a further 72 hr, 10 µL of Resazurin reagent (R&D Systems) was added to each well for a final concentration of 1X in media. Fluorescence, corresponding to the number of live cells per well, was measured 8 hr later. 61 Etoposide resistance genes were identified using the following criteria: an average cell viability < 0.7, an average SSMD < −2, and a two-tailed t-test p-value < 0.05. To assess screen quality, Z-factor was calculated using the means and standard deviations of both positive (siATR) and negative (siNT) controls. The etoposide siRNA screen fell in the excellent range (between 0.5 and 1.0) with a calculated Z-factor of 0.534.

### Pathway and functional enrichment analysis

To characterize the biological processes and pathways associated with etoposide sensitization hits, enrichment and network analyses were performed using curated databases and bioinformatics platforms. Reactome pathway analysis was conducted using the Database for Annotation, Visualization and Integrated Discovery (DAVID 2021). MetaCore analysis (Clarivate Analytics) was used to map sensitization genes to pathway maps and process networks. Gene Oncology (GO) enrichment analysis was performed using the STRING database (v12.0) with GO terms related to biological process, molecular functions, and cellular component analyzed separately. Pathways with a p-value < 0.05 were considered significantly enriched.

### Cell viability assay

Cell viability assays were performed as previously described [6]. Cells were seeded in 96-well plates at the appropriate density (H128: 10,000 cells/well H146: 20,000 cells/well) and were treated with varying doses of etoposide (10-25 μM). For UV treatment, cells were irradiated with the indicated dose of UV prior to seeding. After treatment, cells were grown in 96-well plates for 72 hours and then treated with Resazurin reagent at a final concentration of 1X in media. Cells were then incubated at 37°C for 3 hours. The Resazurin signal was then read as fluorescence (excitation wavelength: 544 nm, emission wavelength:590 nm) on a Synergy H1 microplate reader (BioTek) in conjunction with Gen5 Microplate Reader Software (BioTek). Percent cell viability was quantified relative to the Resazurin signal in the untreated group, including subtracting the background the Resazurin signal for wells with no cells and media with Resazurin only.

### Colony formation assay

U2OS, MDA-MB-231, and BT-549 cells were seeded in 6 well plates, at 100 cells/well for low doses or 200 cells/well for high doses of damage to optimize colony numbers for counting and subsequently normalized. Cells were given 8-24 hours to adhere and then irradiated with IR or treated with continuous treatment with etoposide or sacituzumab govitecan (SG-catalog No. D4002, Batch: D40020). Cells were grown until the untreated group formed colonies of approximately 50 cells in density, which, depending on the doubling time of the cell line, took between 10 and 14 days. To visualize colonies, cells were washed once with ice-cold PBS and then fixed in 3% crystal violet in methanol for 15 minutes. Crystal violet was removed by gently submerging plates in water. Visible colonies were then quantified with Bantex colony counter 920A.

### Cell and protein lysate treatments

Cells were treated with 10 Gy of IR for 4 hrs using X-Ray 320 irradiator (Precision X-Ray Inc., N. Branford, CT). The following drug treatments were used to treat cells: 5 mM Camptothecin (CPT; Sigma # C9911) for 4 hours; 1 µM Shield-1 (Takara # 632189) for 4 hours; 1 µM or 500 nM 4-Hydroxytamoxifen (4-OHT; Sima # H7904) for 4 hours to induce Fok1- or AsiSI-dependent DSB, respectively. Protein lysates for DNA:RNA hybrid pull-down assays were treated with 10 µL RnaseA (Sigma # R6148) or with 10 µL RnaseH (NEB # M0297S) for 30 min at 37°C.

### Antibodies

Primary antibodies used for western blotting, immunoprecipitation (IP) and immunofluorescence (IF) are as follows: HELZ (ThermoFisher customized 1:50 for Western; Proteintech # 26635-1-AP; 1:1000 for Western, 1.2 mL per 1 mg DNA for CHIP, 1:1000 for PLA); S9.6 (Millipore #MABE1095; 1:600 for Slot Blot, 1:250 for IF, 0.5 mL per one million cell nuclei for DRIP-WB, 5U per 1 mg DNA for DRIP); ssDNA (Millipore #MAB3868; 1:1000 for Slot Blot); IgG (Invitrogen # 10500C and Sigma # N103 for CHIP); GFP (Santa Cruz Tech # SC996; 1:1000 for Western; and abcam # ab290; 1:5000 for Western, 1 mg per 2 mg lysate for IP); γH2AX (Cell Signaling Technology # 2577S; 1:500 for IF; Millipore # 05-636; 1:6000 for IF; and Active Motif # 39117; 1 mL per 1 mg DNA for CHIP); KU80 (abcam # 80592; 1:1000 for Western); GAPDH (Santa Cruz tech # sc-47724; 1:1000 for Western); 53BP1 (Bethyl # A300-273A; 1:1000 for IF); RPA32 (Santa Cruz Tech # sc-14692; 1:400 for Western); pRPA32 S4/8 (Bethyl # A700-009; 1:1000 for Western); RPA70 (Cell Signaling # 2267S; 1:120 for IF); BRCA1 (Millipore # 07-434; 1:1000 for Western); CtIP (Millipore # MABE1060; 1:1000 for Western); Senataxin (Novus biological # NB100-57542; 1:1000 for IF) Rad51 (abcam # ab176458; 1:1000 for IF); Lamin A/C (Cell Signaling #2032; 1:1000 for Western) RFP (abcam # ab125244; 1:1000 for Western); α-Tubulin (Sigma # T6074; 1:10,000 for Western); and BrdU (BD Biosciences # 347580; 1:200 for IF). Secondary antibodies used for Western (at 1:1000) are: donkey anti-rabbit IR Dye 800 (Licor Biosciences #926-32213); donkey anti-rabbit IR Dye 680 (Licor Biosciences # 926-68023); donkey anti-mouse IR Dye 800 (Licor Biosciences # 926-32213); donkey anti-mouse IR Dye 680 (Licor Biosciences # 926-68022). Secondary antibodies used for IF (at 1:1000) are: goat anti-mouse Alexa Fluor 555 (Invitrogen # A21424); goat anti-rabbit Alexa Fluor 488 (Invitrogen # A11034); goat anti-mouse Alexa Fluor 647 (Invitrogen # A21235).

### Plasmids, siRNA and sgRNA

Plasmids expressing full length HELZ-Myc-GFP were generated by cloning HELZ-Myc into the BamH1 and EcoR1 restriction sites of pcDNA3.1-GFP (Addgene # 70219) and RFP (Addgene # 13032), respectively (done by Emory Integrated Genomics Core). The plasmid was used to purify HELZ and as template for the construction of HELZ-Myc-GFP K674N by QuickChange site-directed mutagenesis (Agilent) for rescue assays. RNaseH1-mCherry WT and D10R/E48R were purchased were purchased (Addgene #60365 and #60367). GFP-BRCA1 was kindly provided by Dr. Xiaochun Yu, RNaseH1-mCherry was purchased (Addgene #60365 and #60367).

The following siRNAs were used: HELZ-2 (CAGCACACCUUGUUAAAUC; Dharmacon # D-021180-02); HELZ-3 (GAUAUCACGUGGAAGACUU; Dharmacon # D-021180-03); HELZ-4 (CUACAGAAGAUCUCGAAUA; Dharmacon # D-021180-04); HELZ-UTR (GGAAAUAGAACGCAUCAAA; Dharmacon #D-021180-01) ATR (GAACAACACUGCUGGUUUG; Dharmacon # D-003202-05), BRCA1 (Dharmacon # (J-003461-09-0010); BRCA2 (GAGACACAAUUACAACUAAA); CTIP (GAGCAGACCUUUCUCAGUA; Dharmacon # D-011376-01); sgHELZ (GATACTTTGGCTGGACGACTGG; Sigma # HS0000373232).

### Transfections

For plasmid transfections, 1-4 million cells were seeded on 60 mm plates and 24 hours post plating, cells were transfected with 2-5 µg of the indicated plasmid using Lipofectamine 3000 as per manufacturer’s instructions (ThermoFisher). Transfection medium was exchanged to fresh medium and cells were moved to 10 cm plates 4-6 hours post-transfection. Cells were harvested 2-4 days post-transfection for downstream assays.

### RNAi silencing

HELZ was silenced using RNAi Max reagent using the protocol outlined by the manufacturer (Invitrogen). In brief, 30-60 nmol siRNA was transfected into one million cells in 60 mm plates. Medium was exchanged to fresh medium 24 hours post transfection and grown for additional 24-48 hours as needed for subsequent assays.

### Protein lysate, fractionation and western blotting

Harvested cells were washed in PBS buffer and then lysed for 30 min on ice in Nonidet P-40 buffer (200 mM NaCl, 1% Nonidet P-40, 50 mM Tris·HCl pH 8.0) freshly supplemented with protease inhibitors. Lysates were clarified by centrifugation (15,700×g, 10 min at 4 °C), and the supernatants were then collected. Protein concentration was determined by Bradford and resolved by SDS/PAGE, transferred onto PVDF, and probed using the desired primary antibodies. Signal was detected using the Li-Cor Odyssey system. Separation of U2OS cellular compartments was performed according to the manufacturer’s instructions with a Standard Cell Fractionation kit (abcam # ab109719). Two different markers were used to assess the purity of each fraction: cytoplasm (GAPDH) and nucleus (Lamin A/C).

### Immunoprecipitation

Protein lysate prepared as described above (750 µg for an over-expressed protein). Benzonase nuclease (Pierce 88701, 500U) or RNase H (NEB # M0297S, 100U) treatment was done on the same lysate for 30 min in 37°C water bath followed by 10 min centrifugation to remove digested debris. Lysate was precleared with 30 µL of protein G or protein A agarose beads (Roche # 10042392 and Invitrogen # 15918-014, respectively) for one hour. Precleared lysate was added to 30 µL protein G or A agarose beads that were prebound to the IP antibody and incubated overnight on a rotator at 4°C. Beads were washed three time with 0.375% CHAPS buffer on a rotator at room temperature and then resuspended in 50/50 0.375% CHAPS buffer and 5X SDS buffer. Resuspended beads were boiled for 10 min at 100°C, centrifuged and supernatant was processed for Western blotting as indicated.

### DNA-RNA hybrid Immunoprecipitation-Western Blot (DRIP-WB)

DRIP-WB assay was performed using Magna Nuclear RIP^TM^ (Native) RNA-Binding Protein Immunoprecipitation Kit (Millipore/Sigma # 17-10522) following the manufacturer’s recommended procedure with modifications. Briefly, same number of cells were counted from each condition to prepare nuclear lysate. The nuclei resuspended in RIP lysis buffer was passed 2X through an insulin syringe, snap-frozen on dry ice. Next, the nuclear lysate was thawed at 37°C and centrifuged at 10,000 x g for 10 min at 4°C. 100 µL from this final nuclear lysate was incubated and rotated overnight at 4°C with 10 µl of magnetic beads resuspended with either S9.6 or IgG antibody. For RNase A or RNase H digestion, the lysate was incubated 30 min at 37°C prior to addition of magnetic beads. Next day, beads were separated placing on magnetic separator for 1 min. Supernatant was discarded and beads were washed twice for 20 min on rotor at 4°C with ice cold Nuclear RIP Wash buffer. The DNA-RNA hybrid was eluted by re-suspending the magnetic beads in 5X SDS-PAGE loading buffer followed by 10 min boiling at 100°C. Supernatant was processed for Western blotting as described above.

### DR-GFP assay

U20S DR-GFP cells were transfected with the siRNA of interest. Twenty-four hours later, medium was removed, and cells were transfected with 3 µg I-SceI and 2 µg of either the empty RFP plasmid or the indicated SAMHD1-RFP rescue plasmid. Seventy-two hours later, cells were harvested, washed twice with PBS, resuspended in PBS and subjected to flow cytometry (Aurora Cytek) for GFP and RFP fluorescence. To measure HR efficiency, percentage of GFP positive cells (HR positive) within the RFP positive cells (i.e. Those transfected with empty or rescue plasmids) was analyzed using the FlowJo software.

### EJ7-GFP assay

HEK293 EJ7-GFP cells were transfected with the indicated siRNA. 24h post transfection, fresh medium was added, and cells were transfected with cas9a and cas9b guide RNA. 72 h post transfection, cells were harvested, washed twice with ice cold PBS. Cell pellet was then resuspended in 500 μl PBS and subjected to flow cytometry (Aurora Cytek) to measure GFP positive cells. Acquired FACS data was analyzed using the FlowJo software.

### BrdU foci DNA end resection assay

U20S cells were transfected with the siRNA of interest and twenty-four hours later, transfected with empty mCherry or mCherry-RNAseH1 WT or mutant plasmid, where indicated. Forty-eight hours after plasmid transfection, cells were incubated with 30 μM BrdU for 36 hours and then incubated for an additional 4 hours in the presence of IR. Cells were incubated in extraction buffer 1 (10 mM PIPES pH 7.0, 300 mM Sucrose, 100 mM NaCl, 3 mM MgCl_2_, 1mM EGTA, 0.5% Triton X-100) for 20 minutes on ice, washed with PBS and then incubated in extraction buffer 2 (10 mM Tris-HCl pH 7.5, 10 mM NaCl, 3 mM MgCl_2_, 1% Tween 40, 0.5% sodium deoxycholate) for 15 minutes on ice. Cells were washed in PBS and subsequently fixed with PFA and permeabilized with Triton X-100 for immunofluorescence with the appropriate antibody. DNA was visualized with DAPI (Southern Biotech # 0100-20) and mCherry-RNaseH1 was observed by mCherry fluorescence.

### Immunofluorescence

Appropriate number of U2OS cells were transfected with the indicated siRNA and seeded on coverslips. Overexpression with plasmids was performed 24 h later if mentioned for the particular experiment. Treatment with IR or drug was done 4 h prior to fixation. For U20S-265 mCherry-LacI-Fok1 cells, treatment with 1 µM Shield-1 and 1 µM 4-OHT was done for 4 hours to induce recombinant Fok1 expression. 72 h after siRNA, the coverslips were washed with PBS, fixed in 4% paraformaldehyde for 20 minutes at room temperature and permeabilized with 0.5% Triton X-100 for 10 minutes on ice. Blocking with 5% BSA was done for 1 h following with overnight incubation with the indicated primary antibodies at 4°C. For RNase H (NEB # M0297S) or RNase A (Sigma #R6148) treatment, cells were treated at 10U in 500 ul PBS or 40 μg in 500 μl PBS respectively for 30-60 min at 37°C following permeabilization in live cells before fixation with PFA (40 µg in 500 µl phosphate buffer saline (PBS) as adapted^61^. Secondary antibody staining was performed for 1 h. DNA was stained with DAPI and coverslips were mounted on slides. Immunofluorescence signals were visualized using a 63x oil objective on a Zeiss Observer Z1 microscope equipped with Axiovision Rel 4.8 software. RNA-DNA hybrid immune-detection was conducted as described^105^. Briefly, coverslips of attached U2OS cells were washed with PBS and fixed with ice-cold 100% methanol for 10 min at −20°C, followed by 1 min incubation with cold acetone at room temperature. Coverslips were then washed three times with SSC 4X buffer and blocked in 4X SSC+3% BSA for 1 h. Primary antibody S9.6 (1:250) was incubated overnight at 4°C. The subsequent steps were as described above. Enzymatic treatments with RNase H or RNase A were performed after methanol/acetone fixation for 30 minutes at 37C in PBS and then rinsed in SCC 4X twice before blocking.

### Proximity Ligation Assay (PLA)

PLA experiments were performed using the DuoLink In Situ Red Starter Kit Mouse/Rabbit (Millipore/Sigma-Aldrich # DUO92101-1KT) and following manufacturer protocol. U2OS cells were knocked down for the gene of interest, seeded on coverslips and treated with etoposide after 72 h for 4 h if indicated. For HELZ or γH2AX/S9.6 PLA, coverslips containing U2OS cells were processed following the RNA-RNA hybrid immune-detection described above.

### RNA-DNA hybrid immunodetection by slot blot

Cells were harvested after 72 h siRNA knockdown with or without 4 h etoposide treatment and genomic DNA was isolated with DNeasy Blood and Tissue kit (Qiagen # 69504). Nucleic acid was measured by nanodrop, 0.5 µg DNA was used from each condition. DNA was treated with 0.5 N NaOH and 1.5 M NaCl for 10 min to denature the DNA and neutralized for another 10 min in 0.5 M Tris-HCl buffer (pH 7.0) containing 1 M NaCl. 5U RNase H (NEB #M0297S) per 1 μg DNA treatment was done on the same amount of DNA followed by 1 h incubation at 37°C. All samples were then brought up to a final volume of 200 µl with 20X SSC buffer and blotted onto presoaked Nitrocellulose membrane in 20X SSC buffer using a slot blot apparatus (Whatman Schleicher & Schuell Minifold I). After UV-cross linking (0.12 J/m2), we blocked in 5% BSA. Next, the non-treated and RNase H treated part of the membrane were subjected to blotting with S9.6 while the treated membrane serving as loading control was stained with single-stranded DNA antibody.

### Micronuclei assay

Cells were knock downed for HELZ and seeded on coverslips. In the next 48 h, cells were treated with 3 μg/ml cytochalasin B or left untreated for 12 hours, prior to fixation at 72 hours post knockdown. Cells were fixed in 4% paraformaldehyde, stained with DAPI, mounted on coverslips and imaged. Percent of binucleated cells with one or more DAPI positive micronucleus were quantified. 300 binucleated cells were counted per group.

### Cell cycle analysis

Cells were knocked down for the gene of interest. 72 h later, they were harvested, washed and resuspended in cold PBS followed by fixation with cold 70% ethanol on ice for a minimum of 1 h. After washing twice in PBS, cells were treated with RNase A and stained with propidium iodide at room temperature for a minimum of 30 min. Finally, at least 25,000 cells were analyzed for propidium iodide fluorescence on a flow cytometer (Cytek Aurora). Debris and aggregates were excluded throughout analysis by FlowJo software.

### ChIP assay

ChIP assay was performed using ChromaFlash^TM^ One-Step ChIP Kit (P-2025, Epigentek, Farmingdale, NY, USA) according to the manufacture’s instruction. Briefly, DivA cells were treated with Shield-1 and 4-OHT for 4 hours to induce DSBs, then fixed with 1% formaldehyde (Sigma-252549) for 15 mins at room temperature. After two washes with cold PBS, 0.125 M glycine was added for 5 minutes to quench formaldehyde reaction. Next, the cells were washed twice with PBS lysed by 0.75% CHAPS in lysis buffer (10% glycerol, 150 nM NaCl, 50 nM Tris pH-7.5). The lysed cells were sonicated to an average size of 500-800 base pairs using a Branson microtip SFX250 sonicator. Antibodies described above were used for immunoprecipitation for 4 hours at room temperature with IgG as the negative control. The obtained immunoprecipitated chromatin was subjected to quantitative real-time (PowerTrack™ SYBR Green Master Mix A46012, PCR 7500 Fast Real-Time PCR system, ThermoFisher) using primers specific for no-DSBs, HR1 and HR2 prone sites in DIvA cells.

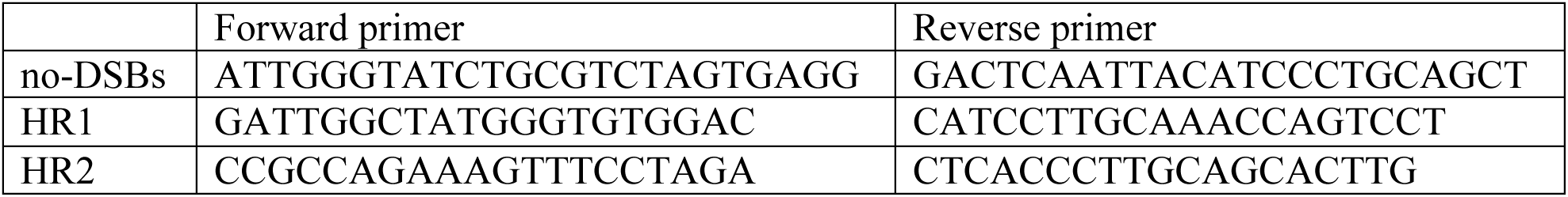

### DNA-RNA hybrid Immunoprecipitation-sequencing (DRIP-seq)

Genomic DNA was extracted from ∼2 million U2OS cells transfected with siNC or siHELZ and fragmented with restriction enzymes (BsrGI, EcoRI, HindIII, SspI, XbaI, 30 U/each, NEB) following a published protocol ^106^. Fragmented nucleic acids were recovered by phenol-chloroform extraction. Four ug of fragmented gDNA was digested with 20 U of RNase H (NEB #M0297S) at 37°C for 6 hr to serve as a negative control. Four μg of fragmented gDNA or RNase H digested DNA were immunoprecipitated with 8 μg of S9.6 antibody (Millipore #MABE1095) overnight at 4°C in DRIP binding buffer (10 mM sodium phosphate, 140 mM sodium chloride, 0.05% (v/v) Triton-X-100, pH 7.0). DNA-antibody complex was then incubated with 40 μL of Protein G beads (Invitrogen #10003D) for 2 hr at 4°C, and beads were washed three times with DRIP binding buffer. Antibody-captured DNA-RNA hybrids were then eluted in (50 mM Tris, 10 mM EDTA, 0.5% (v/v) SDS, pH 8.0) at 55°C for 45 min and subjected to phenol-chloroform extraction. The immunoprecipitated DNA-RNA hybrids were quantified by Qubit (ThermoFisher Scientific #Q32854). Ten ng of DNA-RNA hybrids were treated with 5 U of RNase H for 1 h and then were sonicated into small fragments (∼100-500 bp) using a Covaris Focused-Ultrasonicator Me220. Fragmented DNA was blunted, 5′ phosphorylated and 3’ A-tailed by NEBNext Ultra II End Prep Enzyme Mix following the manufacturer’s instruction (New England Biolabs #E7645L). An adaptor was ligated, and USER enzyme was used for U excision to yield adaptor ligated double-strand DNA. The DNA was then PCR-amplified using barcoded PCR primers (New England Biolabs #E7335L, #E7500L). After AMPure XP bead purification (Beckman Coulter #A63881) and Qubit quantification, the libraries were sent to Admera Health for quality analysis and sequencing. Equimolar pooling of libraries was performed, and 8-12 samples were pooled in one lane. Libraries were sequenced on a HiSeq with a read length configuration of 150 PE, targeting 740 M total reads per lane (370 M in each direction).

### Bioinformatic analysis for DRIP-seq

DRIP-seq reads were aligned to the human genome sequence (hg38) by Bowtie2 version 2.4.4 with default parameter ^107^. Aligned reads in the bam file were sorted by genomic coordinate by SAMtools ^108^. Peaks for each library were identified by MACS2 version 2.2.6 compared to their corresponding input bam file ^109^. DRIP-seq peaks identified from siNT and siHELZ were merged and reads were re-counted in the merged peak regions. Differential analysis was conducted by DESeq2 based on the reads count matrix in the merged peak regions and significantly changed regions were defined by FDR<0.05 ^110^. The differential regions were annotated to genes by annotatePeaks.pl from HOMER version 4.11 ^111^.

### RNA/DNA hybrid with 5’ overhangs, EMSA binding assay and HELZ unwinding assay RNA/DNA hybrid substrate

RNA/DNA hybrids with 5’ overhangs were assembled from the oligonucleotides Cy5-UCGUAGCUCGGGAGUGCACCAGAUUCAGCAAUUAAGCUCUAAGCC and GTCACTTGATAAGAGGTCATTTGAATTCATGGCTTAGAGCTTAATTGCTGAATCTGGTGCTGGGATCCAACATGTTTTAAATATGCAATG, as previously described [REF Zhao Steinfeld Nature2017].

### Biotinylated nucleic acid pull-down assay

HEK293T or HEK293T cells expressing HELZ-Myc-GFP were washed with PBS and cell lysates were prepared by resuspending in Buffer B (10mM Tris-HCl pH 7.5, 100mM NaCl, 10% Glycerol, 10ug/mL BSA, 0.05% NP40, CHAPS 0.35%, protease inhibitors) and incubating on ice for 20 minutes. Lysates were clarified by centrifugation and protein concentration in the supernatant was determined by Bradford assay. Whole cell lysates (600 μg – 1 mg) were precleared with 30 μL streptavidin-conjugated agarose beads (Millipore # 69203) for one hour at 4°C, where the beads had been pre-washed in Buffer A (10 mM Tris-HCl pH 7.5, 100 mM NaCl). Precleared lysates were incubated with 30 μL streptavidin-conjugated beads pre-bound to 40 pmol biotinylated RNA/DNA overnight at 4°C. As a negative control, precleared lysate was also incubated with beads not prebound to biotinylated RNA/DNA. Biotinylated RNA/DNA used were single-stranded RNA or DNA (ss), double-stranded (ds), 5’ RNA overhang or 3’ RNA overhang and sequences are as previously described above. Lysate-beads-biotinylated samples were washed with Buffer B four times, boiled in 5X SDS sample buffer and western blotted with anti-GFP or anti-α-tubulin antibody.

### EMSA binding assay

A 5 nM RNA/DNA hybrid substrate was mixed with HELZ protein at the concentrations indicated in Figure S5, in a buffer containing 10 mM Tris-HCl (pH 7.5), 60 mM KCl, 100 ng/μl BSA, 2 U/μl RNaseIN (Promega), and 1 mM DTT, to a final volume of 12.5 μl. This mixture was incubated on ice for 30 minutes, followed by the addition of 2.5 μl of loading buffer (50% glycerol, 20 mM Tris-HCl (pH 7.4), 0.5 mM EDTA, and 0.05% Orange G). Electrophoretic separation of the protein-bound substrates was performed on 5% native TAE gels at 70 V for 60 minutes at 4°C. The gels were directly imaged using the Cy5 channel of the ChemiDoc MP Imaging System, and the data were analyzed with Image Lab software.

### HELZ unwinding assay

A 5 nM concentration of the RNA/DNA hybrid with 5’ overhangs was mixed with HELZ protein at the concentrations indicated in Figure #. The reaction buffer consisted of 10 mM Tris-HCl (pH 7.5), 60 mM KCl, 100 ng/μl BSA, 1 mM MgCl2, 2 mM ATP, and 1 mM DTT, with a final volume of 12.5 μl. The mixture was incubated at 37°C for 15 minutes. The reaction was terminated by adding a 200-fold molar excess of competitor oligonucleotide (trap DNA: GGCTTAGAGCTTAATTGCTGAATCTGGTGC) to trap the unwound oligonucleotides and prevent reannealing. Additionally, 2.5 μl of loading buffer (composed of 50% glycerol, 20 mM Tris-HCl (pH 7.4), 0.5 mM EDTA, 0.05% Orange G, 0.5 μl 10% SDS, and 1 μl 1 mg/ml proteinase K) was added, followed by a 15-minute incubation at 37°C. Electrophoretic separation of the unwound substrates was performed on 5% native TAE gels at 70 V for 60 minutes at 4°C. The gels were directly imaged using the Cy5 channel of the ChemiDoc MP Imaging System. Data analysis was conducted using Image Lab software.

### Data Availability

All data are available in the main text or the supplementary materials. Uncropped blots and source data are provided with this paper. DRIPseq GEO accession number GSE242527.

## Funding

National Institutes of Health grant R01CA178999 (DSY)

National Institutes of Health grant R01CA254403 (DSY)

National Institutes of Health grant U54CA274513 (DSY)

National Institutes of Health grant R01CA301614 (DSY)

National Institutes of Health grant P30CA138292 Winship Cancer Institute Shared Resource Core in flow cytometry, cloning, and biostatistics (SSR)

Winship Cancer Institute Cell and Molecular Biology Pilot Award (DSY)

## Author contributions

R.H., A.E.K., and D.S.Y. conceived and designed the study. R.H., A.E.K., S.R., B.W., M.Z.M., Y.H., P.K., M.A., A.S., N.C.L., F.S., J.S.K., T.T., S.K., B.G., and B.S.S. performed and analyzed the experiments. R.H., A.E.K., S.K.R., B.W., M.Z.M., Y.H., P.K., M.A., A.S., N.C.L., F.S., J.S.K., T.T., S.K., B.G., B.S.S., A.M.K., E.C.C., L.Y., B.X., X.L., J.M.S., X.Y., Z.S.B., X.D., K.M.M., B.Y., L.L., W.Z., and D.S.Y. interpreted the findings. R.H. and D.S.Y. wrote the manuscript with input from all authors.

## Competing interests

Authors declare that they have no competing interests.

## Supplementary Materials

Materials and Methods

Figs. S1 to S10

Tables S1 to S2

## Notes

### Competing Interest Statement

The authors have declared no competing interest.

### Summary of Updates

This version of the manuscript has been revised to update the Results and Figures.

## References

1 Sanz, L. A. et al. Prevalent, Dynamic, and Conserved R-Loop Structures Associate with Specific Epigenomic Signatures in Mammals. Molecular cell 63, 167–178, doi:10.1016/j.molcel.2016.05.032 (2016).

2 Garcia-Muse, T. & Aguilera, A. R Loops: From Physiological to Pathological Roles. Cell 179, 604–618, doi:10.1016/j.cell.2019.08.055 (2019).

3 Petermann, E., Lan, L. & Zou, L. Sources, resolution and physiological relevance of R-loops and RNA-DNA hybrids. Nature reviews. Molecular cell biology 23, 521–540, doi:10.1038/s41580-022-00474-x (2022).

4 Aguilera, A. & Gomez-Gonzalez, B. DNA-RNA hybrids: the risks of DNA breakage during transcription. Nat Struct Mol Biol 24, 439–443, doi:10.1038/nsmb.3395 (2017).

5 Brickner, J. R., Garzon, J. L. & Cimprich, K. A. Walking a tightrope: The complex balancing act of R-loops in genome stability. Molecular cell 82, 2267–2297, doi:10.1016/j.molcel.2022.04.014 (2022).

6 Crossley, M. P., Bocek, M. & Cimprich, K. A. R-Loops as Cellular Regulators and Genomic Threats. Molecular cell 73, 398–411, doi:10.1016/j.molcel.2019.01.024 (2019).

7 Li, F. et al. R-Loops in Genome Instability and Cancer. Cancers (Basel) 15, doi:10.3390/cancers15204986 (2023).

8 Niehrs, C. & Luke, B. Regulatory R-loops as facilitators of gene expression and genome stability. Nature reviews. Molecular cell biology 21, 167–178, doi:10.1038/s41580-019-0206-3 (2020).

9 Richard, P. & Manley, J. L. R Loops and Links to Human Disease. J Mol Biol 429, 3168–3180, doi:10.1016/j.jmb.2016.08.031 (2017).

10 Ciccia, A. & Elledge, S. J. The DNA damage response: making it safe to play with knives. Molecular cell 40, 179–204, doi:10.1016/j.molcel.2010.09.019 (2010).

11 Jackson, S. P. & Bartek, J. The DNA-damage response in human biology and disease. Nature 461, 1071–1078, doi:10.1038/nature08467 (2009).

12 Scully, R., Panday, A., Elango, R. & Willis, N. A. DNA double-strand break repair-pathway choice in somatic mammalian cells. Nature reviews. Molecular cell biology 20, 698–714, doi:10.1038/s41580-019-0152-0 (2019).

13 Ceppi, I. et al. Mechanism of BRCA1-BARD1 function in DNA end resection and DNA protection. Nature 634, 492–500, doi:10.1038/s41586-024-07909-9 (2024).

14 Chen, L., Nievera, C. J., Lee, A. Y. & Wu, X. Cell cycle-dependent complex formation of BRCA1.CtIP.MRN is important for DNA double-strand break repair. The Journal of biological chemistry 283, 7713–7720, doi:10.1074/jbc.M710245200 (2008).

15 Cruz-Garcia, A., Lopez-Saavedra, A. & Huertas, P. BRCA1 accelerates CtIP-mediated DNA-end resection. Cell reports 9, 451–459, doi:10.1016/j.celrep.2014.08.076 (2014).

16 Salunkhe, S. et al. Promotion of DNA end resection by BRCA1-BARD1 in homologous recombination. Nature 634, 482–491, doi:10.1038/s41586-024-07910-2 (2024).

17 Schlegel, B. P., Jodelka, F. M. & Nunez, R. BRCA1 promotes induction of ssDNA by ionizing radiation. Cancer research 66, 5181–5189, doi:10.1158/0008-5472.CAN-05-3209 (2006).

18 Yun, M. H. & Hiom, K. CtIP-BRCA1 modulates the choice of DNA double-strand-break repair pathway throughout the cell cycle. Nature 459, 460–463, doi:10.1038/nature07955 (2009).

19 Britton, S. et al. DNA damage triggers SAF-A and RNA biogenesis factors exclusion from chromatin coupled to R-loops removal. Nucleic acids research 42, 9047–9062, doi:10.1093/nar/gku601 (2014).

20 D’Alessandro, G. et al. BRCA2 controls DNA:RNA hybrid level at DSBs by mediating RNase H2 recruitment. Nature communications 9, 5376, doi:10.1038/s41467-018-07799-2 (2018).

21 Li, L. et al. DEAD Box 1 Facilitates Removal of RNA and Homologous Recombination at DNA Double-Strand Breaks. Molecular and cellular biology 36, 2794–2810, doi:10.1128/MCB.00415-16 (2016).

22 Lu, W. T. et al. Drosha drives the formation of DNA:RNA hybrids around DNA break sites to facilitate DNA repair. Nature communications 9, 532, doi:10.1038/s41467-018-02893-x (2018).

23 Ohle, C. et al. Transient RNA-DNA Hybrids Are Required for Efficient Double-Strand Break Repair. Cell 167, 1001–1013 e1007, doi:10.1016/j.cell.2016.10.001 (2016).

24 Ouyang, J. et al. RNA transcripts stimulate homologous recombination by forming DR-loops. Nature 594, 283–288, doi:10.1038/s41586-021-03538-8 (2021).

25 Wei, W. et al. A role for small RNAs in DNA double-strand break repair. Cell 149, 101–112, doi:10.1016/j.cell.2012.03.002 (2012).

26 Cohen, S. et al. Senataxin resolves RNA:DNA hybrids forming at DNA double-strand breaks to prevent translocations. Nature communications 9, 533, doi:10.1038/s41467-018-02894-w (2018).

27 Marnef, A. & Legube, G. R-loops as Janus-faced modulators of DNA repair. Nat Cell Biol 23, 305–313, doi:10.1038/s41556-021-00663-4 (2021).

28 Ortega, P., Merida-Cerro, J. A., Rondon, A. G., Gomez-Gonzalez, B. & Aguilera, A. DNA-RNA hybrids at DSBs interfere with repair by homologous recombination. Elife 10, doi:10.7554/eLife.69881 (2021).

29 Sessa, G. et al. BRCA2 promotes DNA-RNA hybrid resolution by DDX5 helicase at DNA breaks to facilitate their repairdouble dagger. EMBO J 40, e106018, doi:10.15252/embj.2020106018 (2021).

30 Fairman-Williams, M. E., Guenther, U. P. & Jankowsky, E. SF1 and SF2 helicases: family matters. Curr Opin Struct Biol 20, 313–324, doi:10.1016/j.sbi.2010.03.011 (2010).

31 Hanet, A. et al. HELZ directly interacts with CCR4-NOT and causes decay of bound mRNAs. Life Sci Alliance 2, doi:10.26508/lsa.201900405 (2019).

32 Hasgall, P. A. et al. The putative RNA helicase HELZ promotes cell proliferation, translation initiation and ribosomal protein S6 phosphorylation. PLoS One 6, e22107, doi:10.1371/journal.pone.0022107 (2011).

33 Azzalin, C. M. & Lingner, J. The human RNA surveillance factor UPF1 is required for S phase progression and genome stability. Current biology : CB 16, 433–439, doi:10.1016/j.cub.2006.01.018 (2006).

34 Sakasai, R. et al. Aquarius is required for proper CtIP expression and homologous recombination repair. Sci Rep 7, 13808, doi:10.1038/s41598-017-13695-4 (2017).

35 Sollier, J. et al. Transcription-coupled nucleotide excision repair factors promote R-loop-induced genome instability. Molecular cell 56, 777–785, doi:10.1016/j.molcel.2014.10.020 (2014).

36 Zheng, L. et al. Human DNA2 is a mitochondrial nuclease/helicase for efficient processing of DNA replication and repair intermediates. Molecular cell 32, 325–336, doi:10.1016/j.molcel.2008.09.024 (2008).

37 Milacic, M. et al. The Reactome Pathway Knowledgebase 2024. Nucleic acids research 52, D672–D678, doi:10.1093/nar/gkad1025 (2024).

38 The Gene Ontology, C. The Gene Ontology Resource: 20 years and still GOing strong. Nucleic acids research 47, D330–D338, doi:10.1093/nar/gky1055 (2019).

39 Prakash, L. Lack of chemically induced mutation in repair-deficient mutants of yeast. Genetics 78, 1101–1118, doi:10.1093/genetics/78.4.1101 (1974).

40 al-Khodairy, F., et al. Identification and characterization of new elements involved in checkpoint and feedback controls in fission yeast. Molecular biology of the cell 5, 147–160, doi:10.1091/mbc.5.2.147 (1994).

41 Huen, M. S. et al. RNF8 transduces the DNA-damage signal via histone ubiquitylation and checkpoint protein assembly. Cell 131, 901–914, doi:10.1016/j.cell.2007.09.041 (2007).

42 Mailand, N. et al. RNF8 ubiquitylates histones at DNA double-strand breaks and promotes assembly of repair proteins. Cell 131, 887–900, doi:10.1016/j.cell.2007.09.040 (2007).

43 Jilani, A. et al. Molecular cloning of the human gene, PNKP, encoding a polynucleotide kinase 3’-phosphatase and evidence for its role in repair of DNA strand breaks caused by oxidative damage. The Journal of biological chemistry 274, 24176–24186, doi:10.1074/jbc.274.34.24176 (1999).

44 Haushalter, K. A., Todd Stukenberg, M. W., Kirschner, M. W. & Verdine, G. L. Identification of a new uracil-DNA glycosylase family by expression cloning using synthetic inhibitors. Current biology : CB 9, 174–185, doi:10.1016/s0960-9822(99)80087-6 (1999).

45 Avkin, S., Adar, S., Blander, G. & Livneh, Z. Quantitative measurement of translesion replication in human cells: evidence for bypass of abasic sites by a replicative DNA polymerase. Proceedings of the National Academy of Sciences of the United States of America 99, 3764–3769, doi:10.1073/pnas.062038699 (2002).

46 Garvin, A. J. et al. The deSUMOylase SENP2 coordinates homologous recombination and nonhomologous end joining by independent mechanisms. Genes & development 33, 333–347, doi:10.1101/gad.321125.118 (2019).

47 Lee, M. H., Mabb, A. M., Gill, G. B., Yeh, E. T. & Miyamoto, S. NF-kappaB induction of the SUMO protease SENP2: A negative feedback loop to attenuate cell survival response to genotoxic stress. Molecular cell 43, 180–191, doi:10.1016/j.molcel.2011.06.017 (2011).

48 Burrage, J. et al. The SNF2 family ATPase LSH promotes phosphorylation of H2AX and efficient repair of DNA double-strand breaks in mammalian cells. J Cell Sci 125, 5524–5534, doi:10.1242/jcs.111252 (2012).

49 Shanbhag, N. M., Rafalska-Metcalf, I. U., Balane-Bolivar, C., Janicki, S. M. & Greenberg, R. A. ATM-dependent chromatin changes silence transcription in cis to DNA double-strand breaks. Cell 141, 970–981, doi:10.1016/j.cell.2010.04.038 (2010).

50 Aymard, F. et al. Transcriptionally active chromatin recruits homologous recombination at DNA double-strand breaks. Nat Struct Mol Biol 21, 366–374, doi:10.1038/nsmb.2796 (2014).

51 Wang, I. X. et al. Human proteins that interact with RNA/DNA hybrids. Genome Res. 28, 1405–1414 (2018).

52 Pause, A. & Sonenberg, N. Mutational analysis of a DEAD box RNA helicase: the mammalian translation initiation factor eIF-4A. EMBO J 11, 2643–2654, doi:10.1002/j.1460-2075.1992.tb05330.x (1992).

53 Perez-Calero, C. et al. UAP56/DDX39B is a major cotranscriptional RNA-DNA helicase that unwinds harmful R loops genome-wide. Genes & development 34, 898–912, doi:10.1101/gad.336024.119 (2020).

54 Salas-Armenteros, I. et al. Human THO-Sin3A interaction reveals new mechanisms to prevent R-loops that cause genome instability. EMBO J 36, 3532–3547, doi:10.15252/embj.201797208 (2017).

55 Bryant, H. E. et al. Specific killing of BRCA2-deficient tumours with inhibitors of poly(ADP-ribose) polymerase. Nature 434, 913–917, doi:10.1038/nature03443 (2005).

56 Farmer, H. et al. Targeting the DNA repair defect in BRCA mutant cells as a therapeutic strategy. Nature 434, 917–921, doi:10.1038/nature03445 (2005).

57 Pierce, A. J., Johnson, R. D., Thompson, L. H. & Jasin, M. XRCC3 promotes homology-directed repair of DNA damage in mammalian cells. Genes & development 13, 2633–2638 (1999).

58 Tsunaka, Y., Haruki, M., Morikawa, M. & Kanaya, S. Strong nucleic acid binding to the Escherichia coli RNase HI mutant with two arginine residues at the active site. Biochim Biophys Acta 1547, 135–142, doi:10.1016/s0167-4838(01)00180-7 (2001).

59 Moynahan, M. E., Chiu, J. W., Koller, B. H. & Jasin, M. Brca1 controls homology-directed DNA repair. Molecular cell 4, 511–518, doi:10.1016/s1097-2765(00)80202-6 (1999).

60 Yang, X. et al. RNA-DNA hybrids regulate meiotic recombination. Cell reports 37, 110097, doi:10.1016/j.celrep.2021.110097 (2021).

61 Gritti, I. et al. Loss of ribonuclease DIS3 hampers genome integrity in myeloma by disrupting DNA:RNA hybrid metabolism. EMBO J 41, e108040, doi:10.15252/embj.2021108040 (2022).

62 Hatchi, E. et al. BRCA1 recruitment to transcriptional pause sites is required for R-loop-driven DNA damage repair. Molecular cell 57, 636–647, doi:10.1016/j.molcel.2015.01.011 (2015).

63 Yasuhara, T. et al. Human Rad52 Promotes XPG-Mediated R-loop Processing to Initiate Transcription-Associated Homologous Recombination Repair. Cell 175, 558–570 e511, doi:10.1016/j.cell.2018.08.056 (2018).

64 Matsui, M. et al. USP42 enhances homologous recombination repair by promoting R-loop resolution with a DNA-RNA helicase DHX9. Oncogenesis 9, 60, doi:10.1038/s41389-020-00244-4 (2020).

65 Zhang, X. et al. Attenuation of RNA polymerase II pausing mitigates BRCA1-associated R-loop accumulation and tumorigenesis. Nature communications 8, 15908, doi:10.1038/ncomms15908 (2017).

66 Alzu, A. et al. Senataxin associates with replication forks to protect fork integrity across RNA-polymerase-II-transcribed genes. Cell 151, 835–846, doi:10.1016/j.cell.2012.09.041 (2012).

67 Mischo, H. E. et al. Yeast Sen1 helicase protects the genome from transcription-associated instability. Molecular cell 41, 21–32, doi:10.1016/j.molcel.2010.12.007 (2011).

68 Skourti-Stathaki, K., Proudfoot, N. J. & Gromak, N. Human senataxin resolves RNA/DNA hybrids formed at transcriptional pause sites to promote Xrn2-dependent termination. Molecular cell 42, 794–805, doi:10.1016/j.molcel.2011.04.026 (2011).

69 Ngo, G. H. P., Grimstead, J. W. & Baird, D. M. UPF1 promotes the formation of R loops to stimulate DNA double-strand break repair. Nature communications 12, 3849, doi:10.1038/s41467-021-24201-w (2021).

70 Tran, P. L. T. et al. PIF1 family DNA helicases suppress R-loop mediated genome instability at tRNA genes. Nature communications 8, 15025, doi:10.1038/ncomms15025 (2017).

71 Marabitti, V. et al. ATM pathway activation limits R-loop-associated genomic instability in Werner syndrome cells. Nucleic acids research 47, 3485–3502, doi:10.1093/nar/gkz025 (2019).

72 Chang, E. Y. et al. RECQ-like helicases Sgs1 and BLM regulate R-loop-associated genome instability. J Cell Biol 216, 3991–4005, doi:10.1083/jcb.201703168 (2017).

73 Bjorkman, A. et al. Human RTEL1 associates with Poldip3 to facilitate responses to replication stress and R-loop resolution. Genes & development 34, 1065–1074, doi:10.1101/gad.330050.119 (2020).

74 Hodson, C. et al. Branchpoint translocation by fork remodelers as a general mechanism of R-loop removal. Cell reports 41, 111749, doi:10.1016/j.celrep.2022.111749 (2022).

75 Lafuente-Barquero, J. et al. The Smc5/6 complex regulates the yeast Mph1 helicase at RNA-DNA hybrid-mediated DNA damage. PLoS genetics 13, e1007136, doi:10.1371/journal.pgen.1007136 (2017).

76 Schwab, R. A. et al. The Fanconi Anemia Pathway Maintains Genome Stability by Coordinating Replication and Transcription. Molecular cell 60, 351–361, doi:10.1016/j.molcel.2015.09.012 (2015).

77 Ozdemir, A. Y., Rusanov, T., Kent, T., Siddique, L. A. & Pomerantz, R. T. Polymerase theta-helicase efficiently unwinds DNA and RNA-DNA hybrids. The Journal of biological chemistry 293, 5259–5269, doi:10.1074/jbc.RA117.000565 (2018).

78 Nguyen, D. T. et al. The chromatin remodelling factor ATRX suppresses R-loops in transcribed telomeric repeats. EMBO reports 18, 914–928, doi:10.15252/embr.201643078 (2017).

79 Li, L., Monckton, E. A. & Godbout, R. A role for DEAD box 1 at DNA double-strand breaks. Molecular and cellular biology 28, 6413–6425, doi:10.1128/MCB.01053-08 (2008).

80 Ribeiro de Almeida, C., et al. RNA Helicase DDX1 Converts RNA G-Quadruplex Structures into R-Loops to Promote IgH Class Switch Recombination. Molecular cell 70, 650–662 e658, doi:10.1016/j.molcel.2018.04.001 (2018).

81 Mersaoui, S. Y. et al. Arginine methylation of the DDX5 helicase RGG/RG motif by PRMT5 regulates resolution of RNA:DNA hybrids. EMBO J 38, e100986, doi:10.15252/embj.2018100986 (2019).

82 Bader, A. S. et al. DDX17 is required for efficient DSB repair at DNA:RNA hybrid deficient loci. Nucleic acids research 50, 10487–10502, doi:10.1093/nar/gkac843 (2022).

83 Lin, W. L. et al. DDX18 prevents R-loop-induced DNA damage and genome instability via PARP-1. Cell reports 40, 111089, doi:10.1016/j.celrep.2022.111089 (2022).

84 Hodroj, D. et al. An ATR-dependent function for the Ddx19 RNA helicase in nuclear R-loop metabolism. EMBO J 36, 1182–1198, doi:10.15252/embj.201695131 (2017).

85 Argaud, D., Boulanger, M. C., Chignon, A., Mkannez, G. & Mathieu, P. Enhancer-mediated enrichment of interacting JMJD3-DDX21 to ENPP2 locus prevents R-loop formation and promotes transcription. Nucleic acids research 47, 8424–8438, doi:10.1093/nar/gkz560 (2019).

86 Song, C., Hotz-Wagenblatt, A., Voit, R. & Grummt, I. SIRT7 and the DEAD-box helicase DDX21 cooperate to resolve genomic R loops and safeguard genome stability. Genes & development 31, 1370–1381, doi:10.1101/gad.300624.117 (2017).

87 Mosler, T. et al. R-loop proximity proteomics identifies a role of DDX41 in transcription-associated genomic instability. Nature communications 12, 7314, doi:10.1038/s41467-021-27530-y (2021).

88 Weinreb, J. T. et al. Excessive R-loops trigger an inflammatory cascade leading to increased HSPC production. Dev Cell 56, 627–640 e625, doi:10.1016/j.devcel.2021.02.006 (2021).

89 Marchena-Cruz, E. et al. DDX47, MeCP2, and other functionally heterogeneous factors protect cells from harmful R loops. Cell reports 42, 112148, doi:10.1016/j.celrep.2023.112148 (2023).

90 Chakraborty, P. & Grosse, F. Human DHX9 helicase preferentially unwinds RNA-containing displacement loops (R-loops) and G-quadruplexes. DNA repair 10, 654–665, doi:10.1016/j.dnarep.2011.04.013 (2011).

91 Chakraborty, P. & Hiom, K. DHX9-dependent recruitment of BRCA1 to RNA promotes DNA end resection in homologous recombination. Nature communications 12, 4126, doi:10.1038/s41467-021-24341-z (2021).

92 Chakraborty, P., Huang, J. T. J. & Hiom, K. DHX9 helicase promotes R-loop formation in cells with impaired RNA splicing. Nature communications 9, 4346, doi:10.1038/s41467-018-06677-1 (2018).

93 Cristini, A., Groh, M., Kristiansen, M. S. & Gromak, N. RNA/DNA Hybrid Interactome Identifies DXH9 as a Molecular Player in Transcriptional Termination and R-Loop-Associated DNA Damage. Cell reports 23, 1891–1905, doi:10.1016/j.celrep.2018.04.025 (2018).

94 Yuce, O. & West, S. C. Senataxin, defective in the neurodegenerative disorder ataxia with oculomotor apraxia 2, lies at the interface of transcription and the DNA damage response. Molecular and cellular biology 33, 406–417, doi:10.1128/MCB.01195-12 (2013).

95 Cohen, S. et al. A POLD3/BLM dependent pathway handles DSBs in transcribed chromatin upon excessive RNA:DNA hybrid accumulation. Nature communications 13, 2012, doi:10.1038/s41467-022-29629-2 (2022).

96 Hasanova, Z., Klapstova, V., Porrua, O., Stefl, R. & Sebesta, M. Human senataxin is a bona fide R-loop resolving enzyme and transcription termination factor. Nucleic acids research 51, 2818–2837, doi:10.1093/nar/gkad092 (2023).

97 Boleslavska, B. et al. DDX17 helicase promotes resolution of R-loop-mediated transcription-replication conflicts in human cells. Nucleic acids research 50, 12274–12290, doi:10.1093/nar/gkac1116 (2022).

98 Bardia, A. et al. Final Results From the Randomized Phase III ASCENT Clinical Trial in Metastatic Triple-Negative Breast Cancer and Association of Outcomes by Human Epidermal Growth Factor Receptor 2 and Trophoblast Cell Surface Antigen 2 Expression. J Clin Oncol 42, 1738–1744, doi:10.1200/JCO.23.01409 (2024).

99 Wahby, S. et al. FDA Approval Summary: Accelerated Approval of Sacituzumab Govitecan-hziy for Third-line Treatment of Metastatic Triple-negative Breast Cancer. Clinical cancer research : an official journal of the American Association for Cancer Research 27, 1850–1854, doi:10.1158/1078-0432.CCR-20-3119 (2021).

100 McGuinness, J. E. & Kalinsky, K. Antibody-drug conjugates in metastatic triple negative breast cancer: a spotlight on sacituzumab govitecan, ladiratuzumab vedotin, and trastuzumab deruxtecan. Expert Opin Biol Ther 21, 903–913, doi:10.1080/14712598.2021.1840547 (2021).

101 Schreiber, A. R., Andress, M. & Diamond, J. R. Tackling metastatic triple-negative breast cancer with sacituzumab govitecan. Expert Rev Anticancer Ther 21, 1303–1311, doi:10.1080/14737140.2021.1993065 (2021).

102 Bardia, A. et al. Sacituzumab Govitecan in Metastatic Triple-Negative Breast Cancer. The New England journal of medicine 384, 1529–1541, doi:10.1056/NEJMoa2028485 (2021).

103 Sidaway, P. Sacituzumab govitecan improves OS. Nat Rev Clin Oncol 18, 322, doi:10.1038/s41571-021-00516-x (2021).

104 Bardia, A. et al. Biomarker analyses in the phase III ASCENT study of sacituzumab govitecan versus chemotherapy in patients with metastatic triple-negative breast cancer. Ann Oncol 32, 1148–1156, doi:10.1016/j.annonc.2021.06.002 (2021).

105 Renaudin, X., Lee, M., Shehata, M., Surmann, E.-M. & Venkitaraman, A. R. BRCA2 deficiency reveals that oxidative stress impairs RNaseH1 function to cripple mitochondrial DNA maintenance. Cell Rep. 36, 109478 (2021).

106 Sanz, L. A. & Chedin, F. High-resolution, strand-specific R-loop mapping via S9.6-based DNA-RNA immunoprecipitation and high-throughput sequencing. Nat Protoc 14, 1734–1755, doi:10.1038/s41596-019-0159-1 (2019).

107 Langmead, B. & Salzberg, S. L. Fast gapped-read alignment with Bowtie 2. Nature Methods 9, 357–359, doi:10.1038/nmeth.1923 (2012).

108 Li, H. et al. The Sequence Alignment/Map format and SAMtools. Bioinformatics 25, 2078–2079, doi:10.1093/bioinformatics/btp352 (2009).

109 Zhang, Y. et al. Model-based Analysis of ChIP-Seq (MACS). Genome Biology 9, R137, doi:10.1186/gb-2008-9-9-r137 (2008).

110 Love, M. I., Huber, W. & Anders, S. Moderated estimation of fold change and dispersion for RNA-seq data with DESeq2. Genome Biology 15, doi:10.1186/s13059-014-0550-8 (2014).

111 Duttke, S. H., Chang, M. W., Heinz, S. & Benner, C. Identification and dynamic quantification of regulatory elements using total RNA. Genome Research 29, 1836–1846, doi:10.1101/gr.253492.119 (2019).

