## Supplementary Information for "HELZ is a RNA-DNA helicase that resolves R loops to facilitate homologous recombination repair"

**The PDF file includes:**

Figs. S1 to S10
Tables S1 to S2 (provided as excel files)

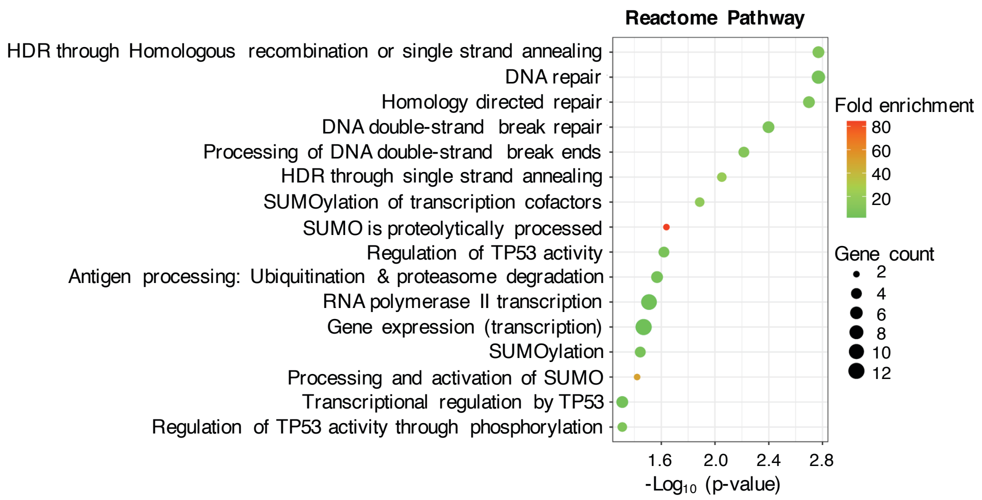


**Figure S1. Reactome pathway analysis**

**Figure S1. Reactome pathway analysis.** Reactome pathway analysis of the 62 etoposide sensitization hits highlights diverse roles in the DDR. The analysis was conducted using DAVID functional annotation bioinformatics tools. Significant pathways ($p < 0.05$) are displayed.

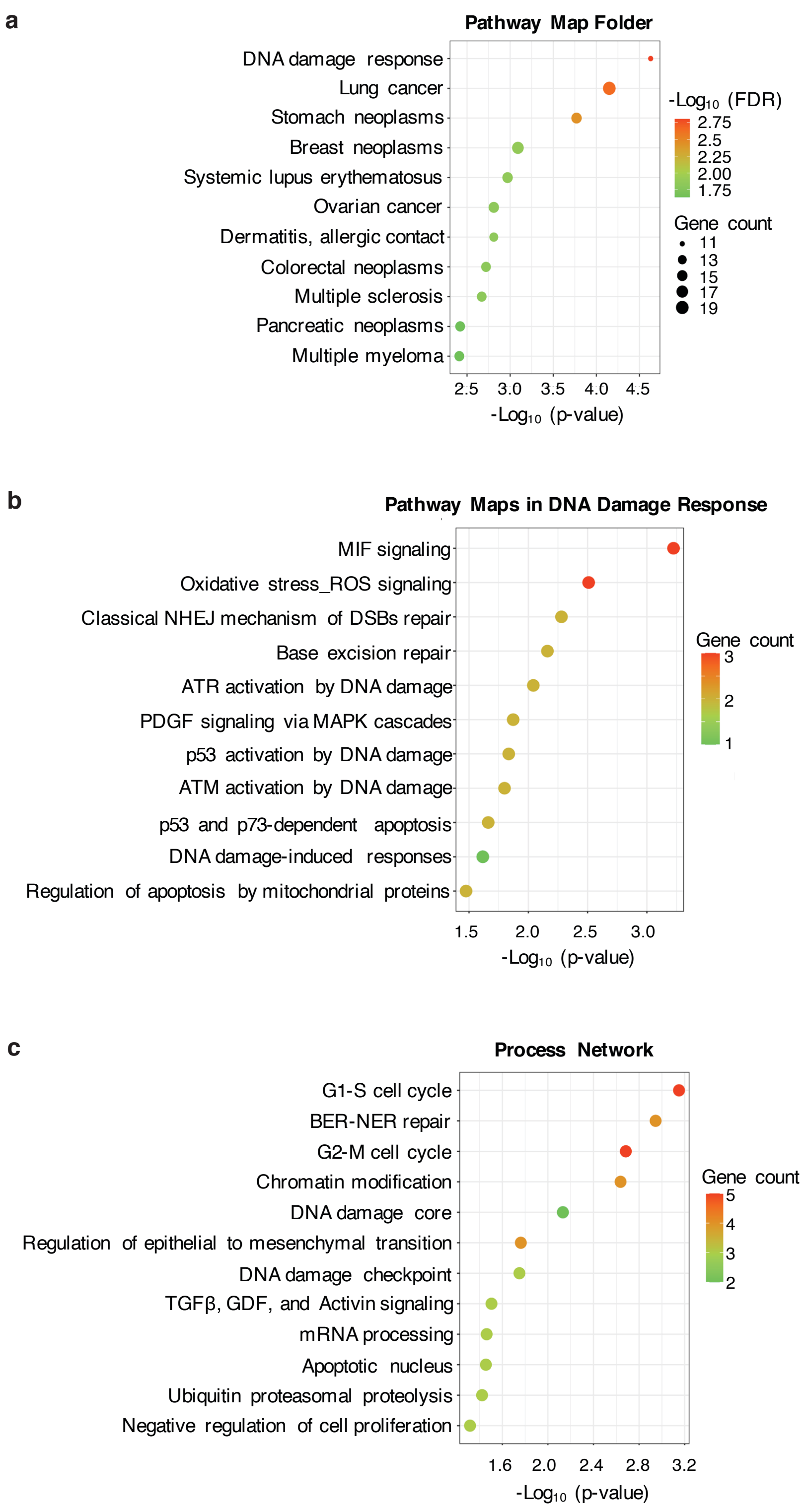


**Figure S2. Metacore pathway map and process network enrichment analysis**

**Figure S2. MetaCore pathway map and process network enrichment analysis.** MetaCore pathway map and process network enrichment analysis based on the 62 sensitization hits. **a** Pathway map folders. **b** Pathway maps focusing on the DNA damage response. **c** Process network.

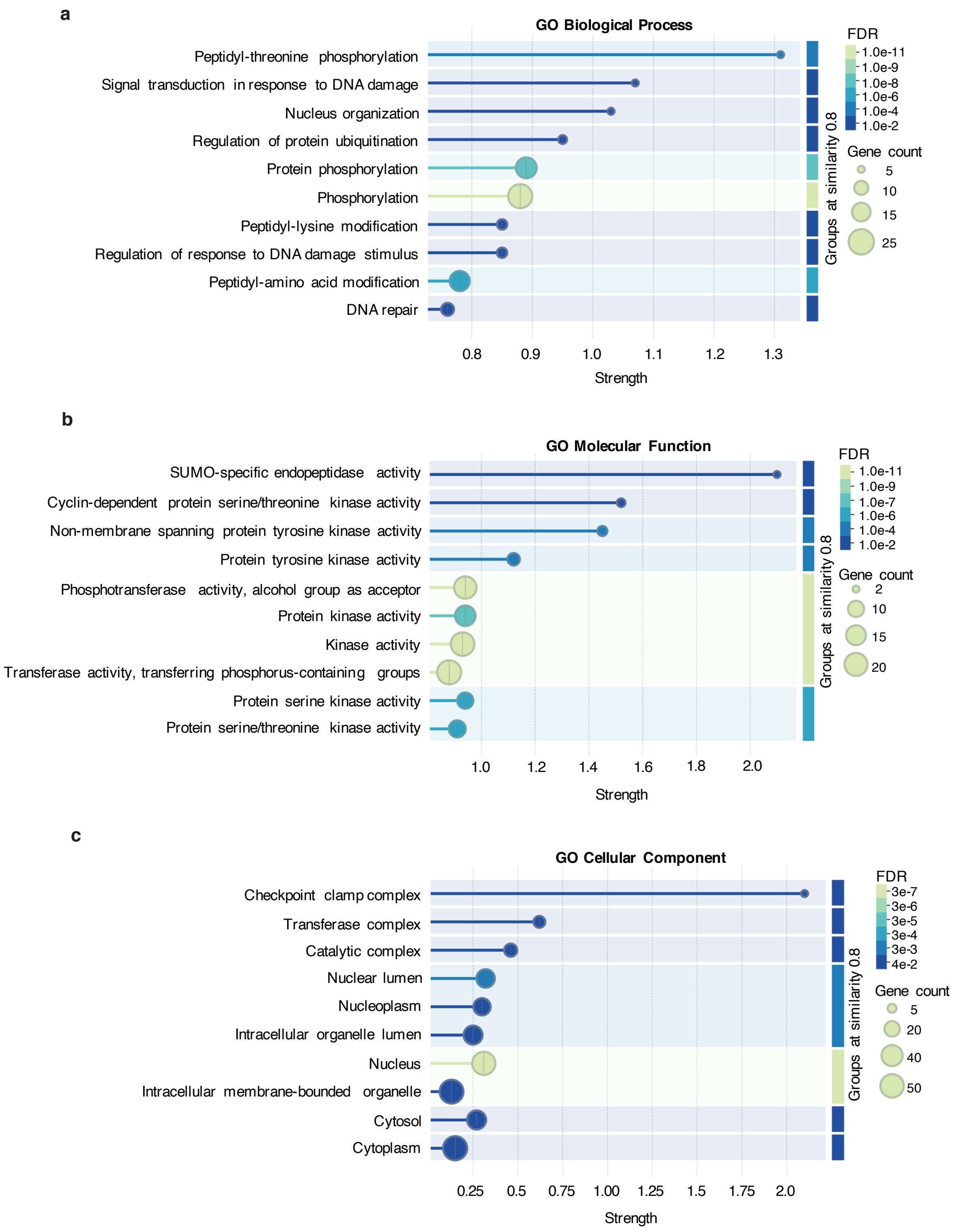


**Figure S3. Gene Ontology enrichment analysis**

**Figure S3. Gene Ontology enrichment analysis.** Gene Ontology enrichment analysis of the 62 sensitization hits performed using the STRING platform. **a** Biological processes. **b** Molecular functions. **c** Cellular components.

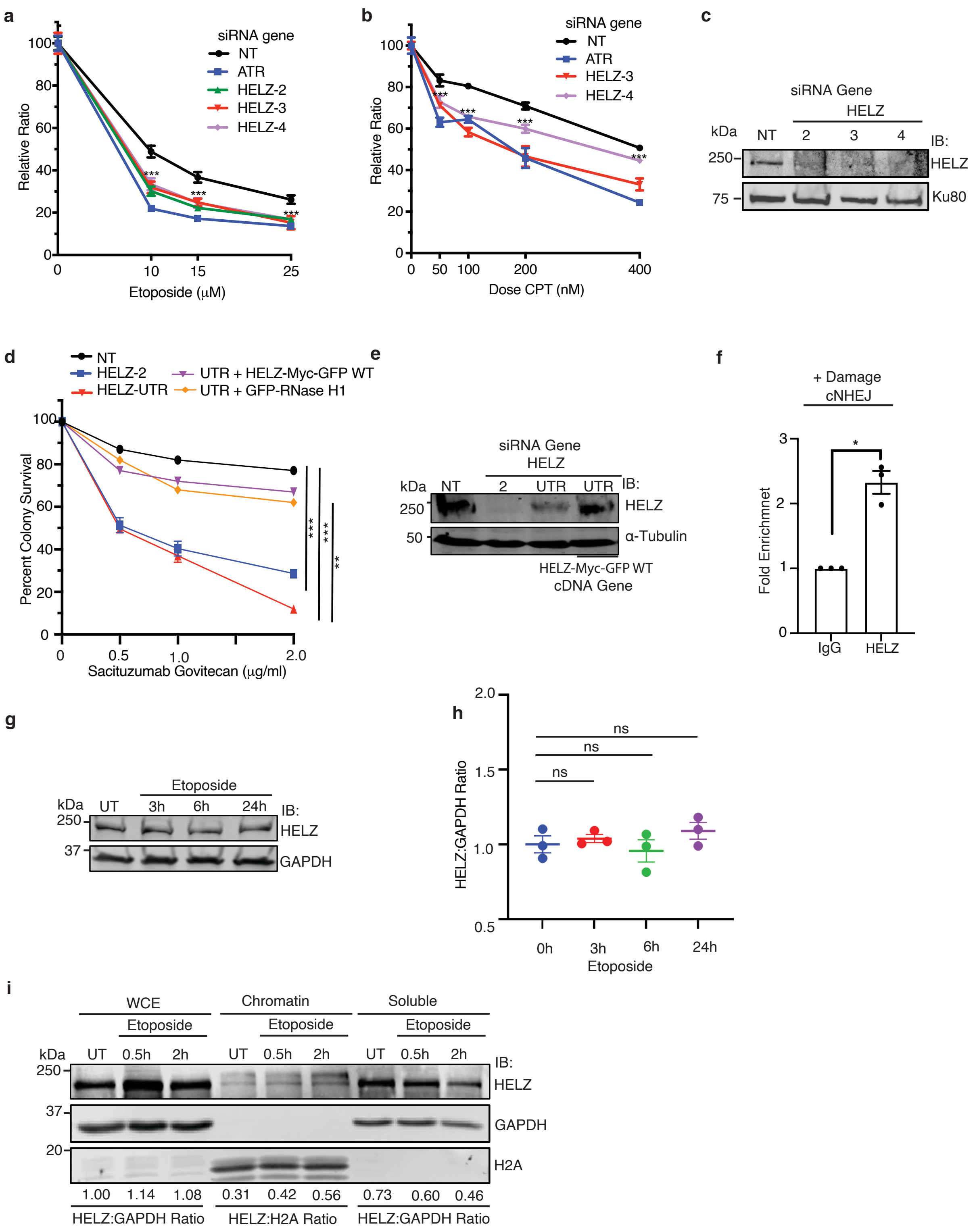


**Figure S4. HELZ depletion causes hypersensitivity to DSB-inducing agents in a R loop dependent manner**

**Figure S4. HELZ depletion causes hypersensitivity to DSB-inducing agents. a-b** HELZ depletion causes hypersensitivity to etoposide (**a**) and CPT (**b**). H128 cells were transfected with siRNA targeting HELZ, ATR, or a NT control. 72 hours after transfection, cells were treated with etoposide or CPT for 72 hours prior to measuring for metabolic activity. Mean and SD from three independent replicas is shown. *** $p < 0.001$. **c** Western blot analysis of HELZ expression in H128 cell demonstrating HELZ knockdown. **d** BT-549 cells were transfected with indicated siRNA. 72 h after transfection, cells were treated with or without indicated doses of SG continuously. Mean and SD from three independent replicas is shown. ** $p <0.01$, *** $p < 0.001$. **e** Western blot analysis showing HELZ knockdown and rescue in MDA-MB-231 cells from (**d**). **f** CHIP-qPCR analysis of endogenous HELZ enrichment at NHEJ prone DNA repair sites in DivA cells. DSBs were induced by treatment with 500 nM 4-OHT for 4 h. Mean and SEM from three independent replicas is shown. **g** Western blot analysis of HELZ expression level in U2OS cells after 20 μM etoposide treatment for indicated time points. **h.** Quantification of HELZ:GAPDH ratio from (**h**). Mean and SD three independent replicas is shown. p was ns. (**i**) U2OS biochemical fractionation after 20 μM etoposide treatment (UT: untreated).

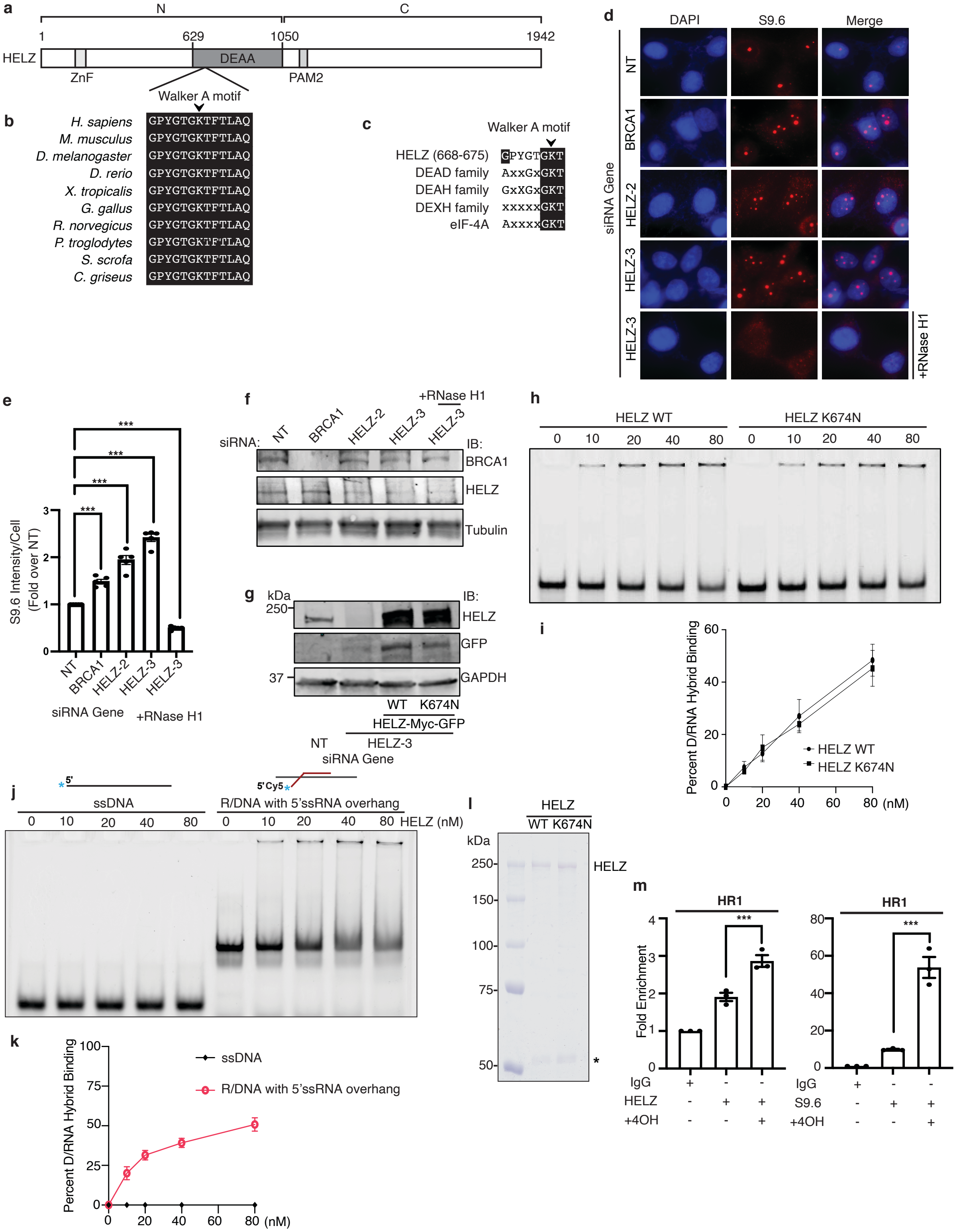


**Figure S5. HELZ prevents R loop accumulation in cells**

**Figure S5. HELZ prevents R loop accumulation in cells.** **a** Schematic representation of human HELZ indicating Zinc finger (ZnF), putative helicase (DEAA: Asp, Glu, Ala, Ala) domain, and polyA-binding protein (PABP) interacting motif 2 (PAM2). HELZ N- and C-terminal fragments are indicated above the scheme. **b** Sequence alignment of Walker A motif in different species. Strictly conserved residues are shown as white letters on black background. K674 is the residue mutated to N to abolish ATPase dependent helicase activity of HELZ. **c** Highly conserved amino acid regions in HELZ compared to DEAD, DEAH, DEXH families and eIF-4A protein. x indicates any amino acid. K in the Walker A motif is often the amino acid mutated to investigate helicase activity of the protein in these families. **d-f** MDA-MB-231 cells were transfected with HELZ siRNA and processed 72 h later for indirect immunofluorescence with fixation, RNaseH treatment for 30 min, and anti-S9.6 antibody. Representative images (**d**) and quantification (**e**) are shown. The median is indicated by a horizontal line. *** $p < 0.001$. **f** Western blot showing HELZ knockdown in MDA-MB-231 cells. **g** Western blot analysis showing HELZ knockdown and overexpression in HCT116 cells. **h-i** EMSA showing comparable binding of insect purified recombinant HELZ WT and K674N with RNA/DNA hybrid with 5' ssRNA overhang. Mean and SD from three independent replicas is shown. **j-k** EMSA showing binding of insect purified recombinant HELZ WT with RNA/DNA hybrid with 5' ssRNA overhang but not ssDNA. Mean and SD from three independent replicas is shown. **l** Coomassie blue gel of insect purified recombinant HELZ WT and K674N. **m** ChIP-qPCR analysis of endogenous HELZ and S9.6 enrichment at HR1 prone DNA repair site. DSBs were induced by treating DIvA cells with 500 nM of 4-OHT for 4 h. Mean and SD from three independent replicas is shown. *** $p < 0.001$.

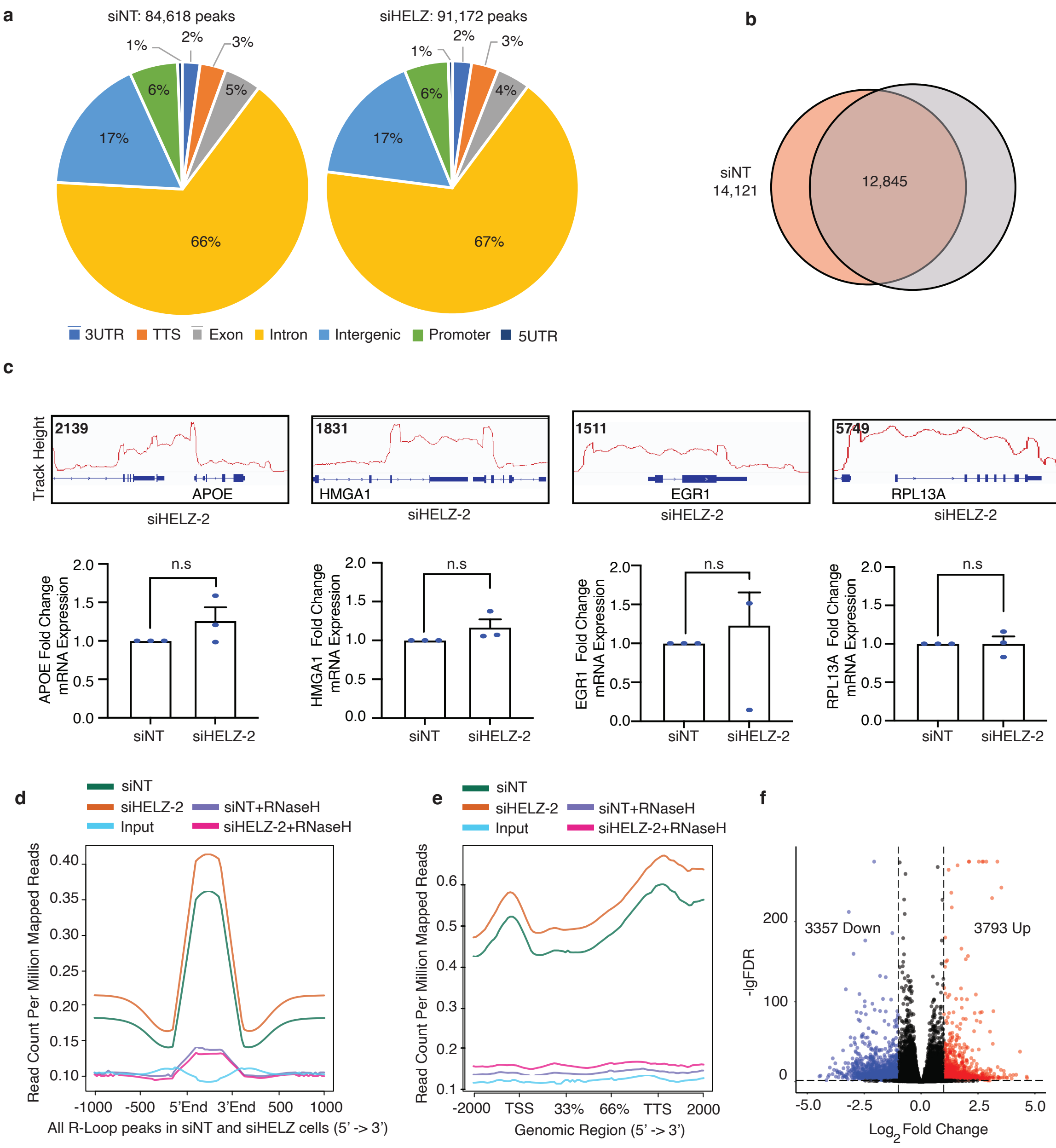


**Figure S6. HELZ depletion leads to genome-wide accumulation of R loops**

**Figure S6. HELZ depletion causes genome wide accumulation of R loops. a** Genomic distribution of R loop regions identified in siNT and siHELZ cells. **b** Venn diagram of the number of genes annotated by R loop regions in siNT and siHELZ cells. **c** R loop upregulation at selected genes with associated mRNA detected by ChIP-qPCR. **d** Ngsplot of R loop read counts across all R loop peaks identified in siNT and siHELZ cells. **e** Ngsplot of R loop read counts across defined human RefSeq (hg38) genes. **f** Volcano plot showing the number of R loop regions increased or decreased in HELZ depleted U2OS cells. The R loop regions that achieved an FDR of $<0.05$ and fold-change of $\geq 2$ are indicated in red (increased) and blue (decreased).

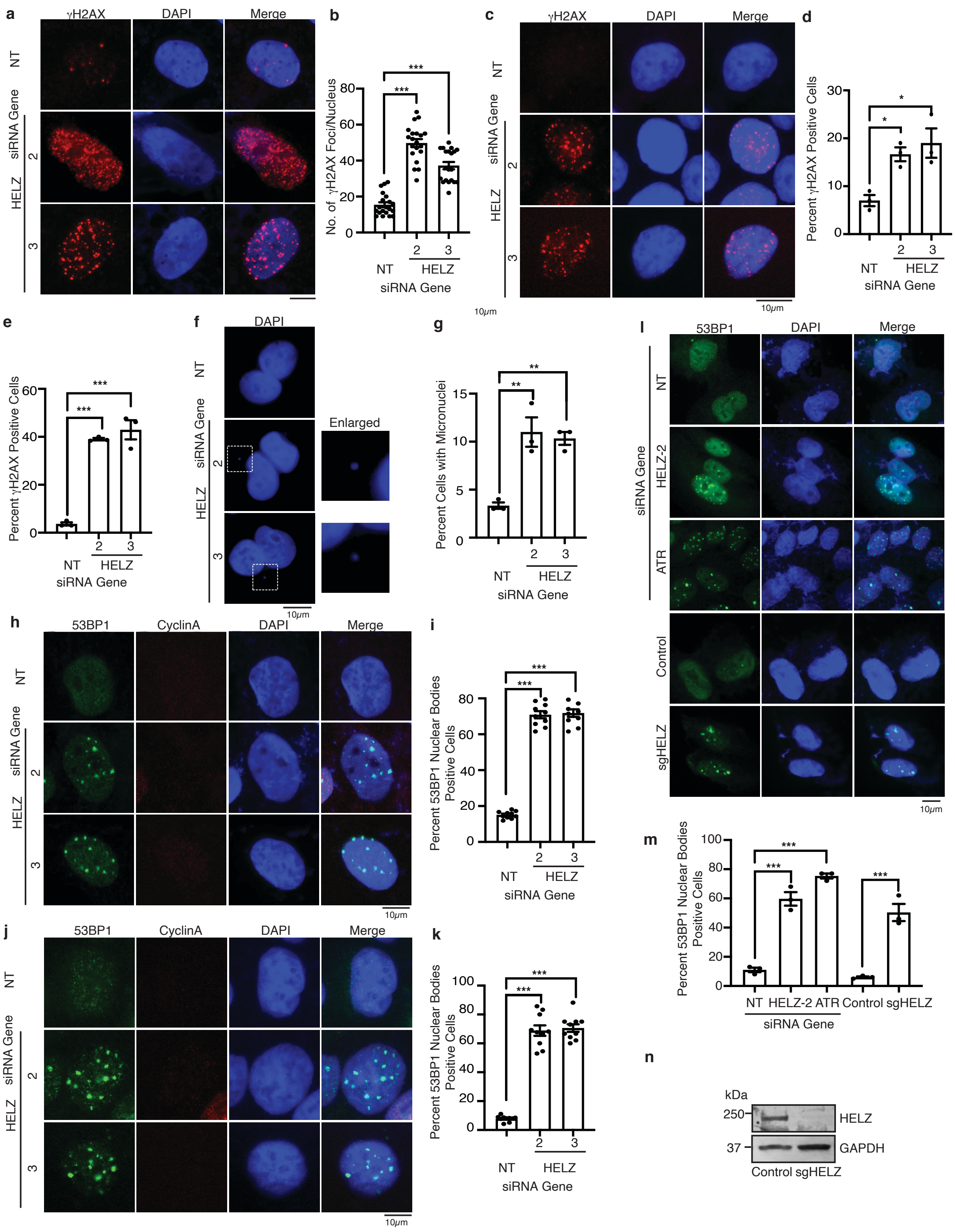


**Figure S7. HELZ deficiency causes genomic instability**

**Figure S7. HELZ deficiency causes genomic instability. a-d** Spontaneous formation of γH2AX foci after 72 h of HELZ depletion in U2OS (**a-b**), RPE-1 cells (**c-d**), and HeLa (**e**). Representative images and quantification are shown. **f-g** Micronuclei formation in U2OS cells 72 h following HELZ depletion. Cells were fixed and stained with DAPI (**f**) and quantified (**g**). **h-k** Representative images and quantification from independent replicas for 53BP1 nuclear bodies in Cyclin A negative U2OS (**h-i**) and HeLa cells (**j-k**) silenced for HELZ or a NT control. *** $p < 0.001$. **l-n** Immunofluorescence staining for spontaneous 53BP1 nuclear bodies was performed in U2OS cells depleted for HELZ or a NT control or in U2OS HELZ KO cells generated by CRISPR/Cas9. Representative images and quantification are shown. **m** Western blot showing KO of HELZ in U2OS cells. For **b**, **d**, **e**, **g**, **i**, **k**, **m**, mean and SD from three independent replicas is shown. * $p < 0.05$, ** $p < 0.01$, *** $p < 0.001$.

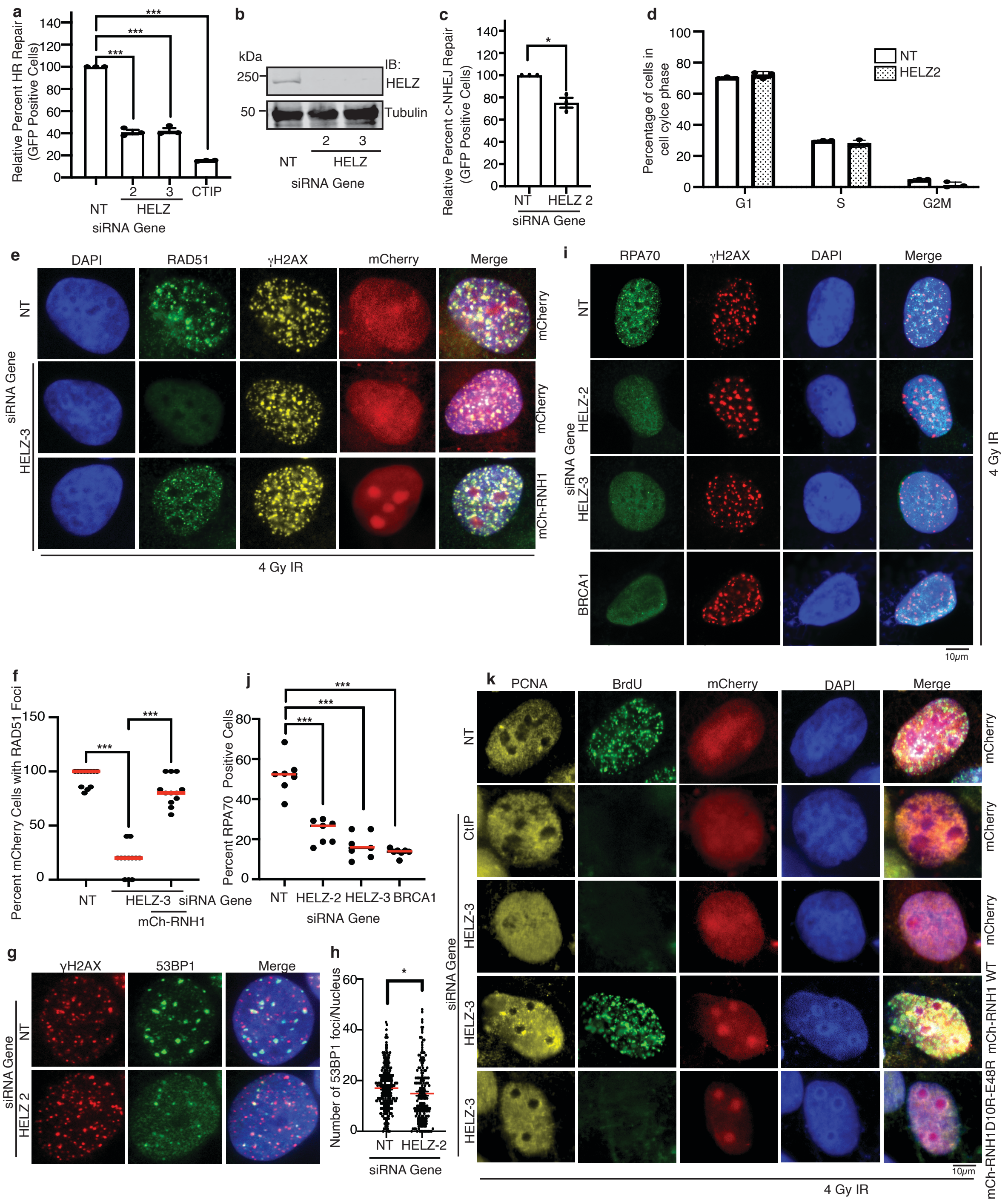


**Supplementary Figure S8. HELZ depletion impairs HR and resection**

**Figure S8. HELZ depletion impairs homologous recombination. a** U2OS cells containing an integrated DR-GFP HR reporter were silenced with indicated siRNA and transfected with I-SceI endonuclease. Live cells collected were subjected to flow cytometry. GFP positive cells were gated to assess for HR. Mean and SD of three replicas are shown. *** $p < 0.001$. **b** Western blot showing HELZ knockdown from **a**. **c** Classical non-homologous end joining (c-NHEJ) repair efficiency in HEK293 cells integrated with the EJ7-GFP reporter, transfected with indicated siRNAs and Cas9/sgRNA targeting the reporter locus. **d** Cell cycle analysis of U2OS cells following depletion of HELZ and CHK1 for 72 h and stained with propidium iodide. **e-f** U2OS cells were transfected with HELZ, CtIP or NT siRNA control for 72h and mCherry-RNaseH1 or mCherry for 48h, treated with 4 Gy IR, and processed after 4h for indirect immunofluorescence with indicated antibodies. Representative images (**e**) and quantification are shown (**f**). The median is indicated by a horizontal line. *** $p < 0.001$**. g-h** U2OS cells were silenced with HELZ or a NT control for 72h, treated with 4 Gy IR, and processed after 4h for indirect immunofluorescence with indicated antibodies. Representative images (**g**) and quantification from three independent replicas are shown (**h**). The median is indicated by a horizontal line. * $p < 0.05$. **i-j** U2OS cells were depleted for HELZ for 72h, treated with 4 Gy IR, and processed 4h later for indirect immunofluorescence with indicated antibodies. Representative images and quantitation are shown. The median is indicated by a horizontal line. *** $p < 0.001$. **k**. U2OS cells were transfected with HELZ, CtIP or NT siRNA control for 72h and mCherry-RNaseH1 WT or D10R/E48R for 48 h, treated with 4 Gy IR, and processed 4h later for indirect immunofluorescence with indicated antibodies.

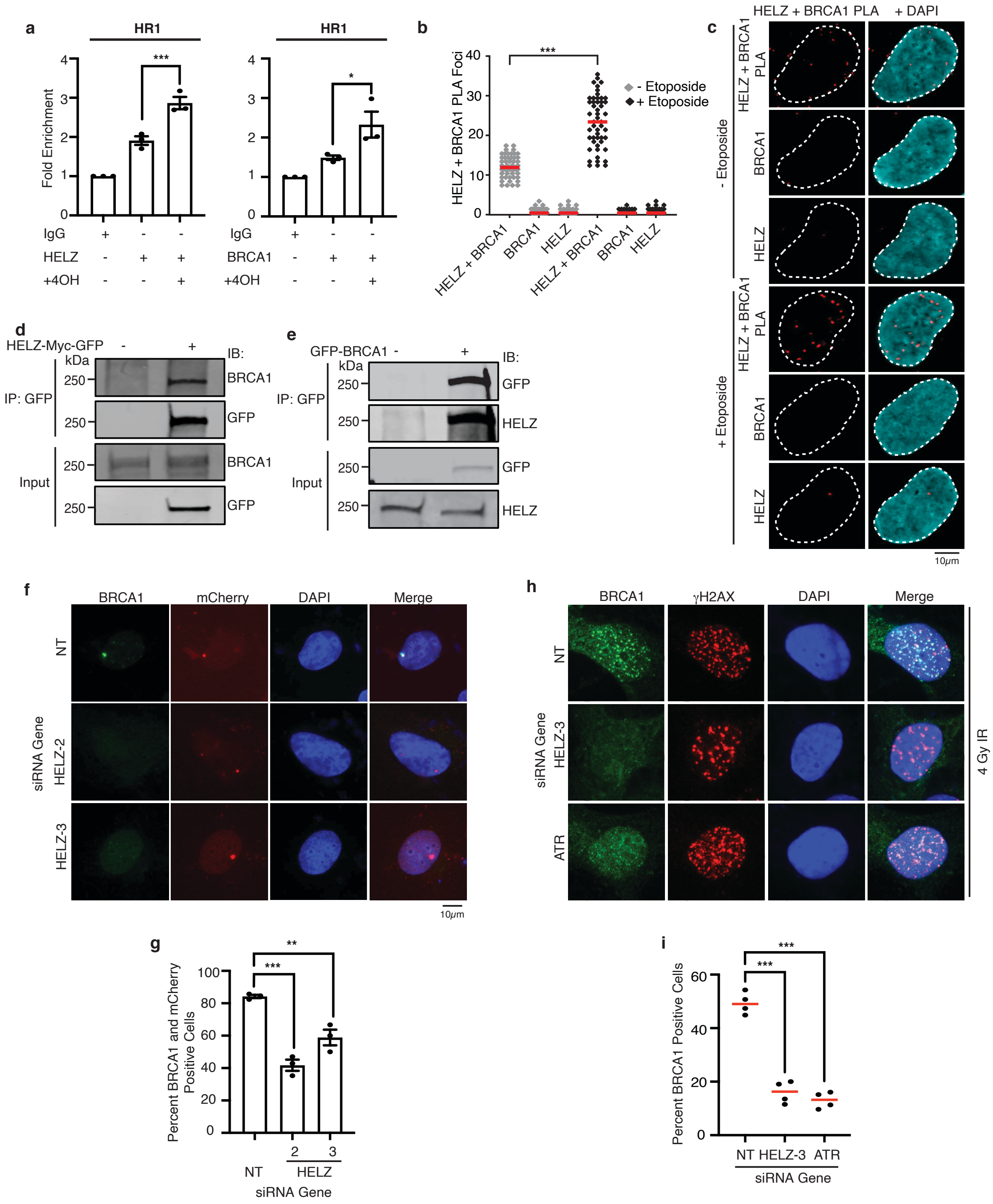


**Supplementary Figure S9. HELZ interacts with BRCA1 in a damage-regulated and RNA-dependent manner and facilitates BRCA1 recruitment to DSBs by preventing accumulation of RNA/DNA hybrids**

**Figure S9. HELZ interacts with BRCA1 in a damage regulated and RNA-dependent manner and facilitates BRCA1 recruitment to DSBs by preventing accumulation of R loops. a** CHIP-qPCR analysis of endogenous HELZ and BRCA1 enrichment at HR1 prone DNA repair site in DIvA cells. DSBs were induced by treatment with 500 nM of 4-OHT for 4 h. Mean and SEM from three replicas is shown. * $p < 0.05$, *** $p < 0.001$. **b-c**. PLA foci suggestive of an interaction between endogenous HELZ and BRCA1 in U2OS cells before and after etoposide treatment (20 μM 4h), including controls with BRCA1 and HELZ alone. Representative images and quantification are shown. The median is indicated by a horizontal line. *** $p < 0.001$. **d-e** Co-IP of HELZ-Myc-GFP or GFP-BRCA1 expressed in HEK293T cells pulls down endogenous BRCA1 or HELZ respectively. **f-g** U2OS-265 Fok1 cells were depleted for HELZ. After 72 h, DSBs were induced by Shield-1 and 4-OHT. Cells were fixed after 4 h and stained with indicated antibodies. Percentage of cells with both a single red focus and green focus of BRCA1 were quantified. Representative images and quantification are shown. Mean and SD of three replicas are shown.** $p < 0.01$, *** $p < 0.001$. **h-i** U2OS cells were transfected with HELZ, ATR, or NT siRNA, treated with 4 Gy IR, and fixed after 4 h for indirect immunoflourescence with indicated antibodies. Representative images and quantification from three independent replicas are shown. The median is indicated by a horizontal line. *** $p < 0.001$.

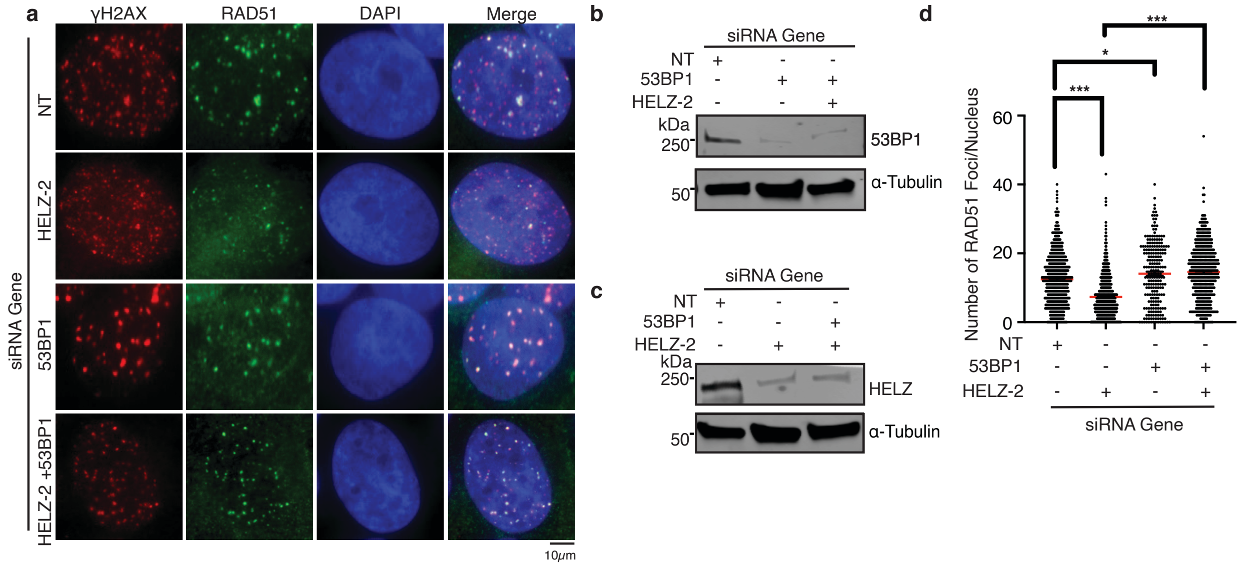


**Supplementary Figure S10. 53BP1 depletion rescues the impairment in IR-induced RAD51 foci of HELZ depletion**

**Figure S10. 53BP1 depletion rescues the impairment in IR-induced RAD51 foci of HELZ depletion. a-d** U2OS cells were silenced with HELZ, 53BP1, or a NT control for 72h, treated with 4 Gy IR, and processed after 4h for indirect immunofluorescence with indicated antibodies. Representative images (**a**) and quantification from three independent replicas are shown (**d**). The median is indicated by a horizontal line. * $p < 0.05$, *** $p < 0.001$**.** **b-c** Western blot showing HELZ and 53BP1 knockdown in U2OS cells in (**a**).

**Table S1.**
Provided as excel file

**Table S2.**
Provided as excel file

Primary Etoposide Hypersensitivity siRNA Screen

| Gene symbol | Gene ID | Average Treated/Untreated Viability (Normalized to NS = 1) (this was threshold) | Standard Deviation (n=3) | SSMD | /alue (Two-Tailed T-Te |
|---|---|---|---|---|---|
| CETN1 | 1068 | 0.273 | 0.030 | -7.433 | 0.0002 |
| LYK5 | 92335 | 0.366 | 0.073 | -5.366 | 0.0039 |
| DKFZP761P0423 | 157285 | 0.400 | 0.187 | -2.866 | 0.0308 |
| HIPK2 | 28996 | 0.437 | 0.138 | -3.385 | 0.0190 |
| USP30 | 84749 | 0.438 | 0.246 | -2.139 | 0.0580 |
| PTGS2 | 5743 | 0.446 | 0.093 | -4.205 | 0.0088 |
| CDKL3 | 51265 | 0.446 | 0.055 | -5.120 | 0.0026 |
| siATR Control | siATR Control | 0.460 | 0.013 | -5.744 | 0.0000 |
| TXNL4 | 10907 | 0.463 | 0.106 | -3.800 | 0.0123 |
| SENP2 | 59343 | 0.470 | 0.190 | -2.511 | 0.0398 |
| FBXL18 | 80028 | 0.470 | 0.078 | -4.374 | 0.0065 |
| SUMO4 | 387082 | 0.474 | 0.105 | -3.752 | 0.0125 |
| LIPI | 149998 | 0.492 | 0.063 | -4.530 | 0.0043 |
| PNKP | 11284 | 0.501 | 0.105 | -3.551 | 0.0140 |
| LMTK2 | 22853 | 0.504 | 0.065 | -4.356 | 0.0050 |
| ZMPSTE24 | 10269 | 0.506 | 0.129 | -3.097 | 0.0217 |
| EHMT2 | 10919 | 0.511 | 0.179 | -2.424 | 0.0415 |
| SUMO2 | 6613 | 0.512 | 0.067 | -4.265 | 0.0054 |
| PIAS4 | 51588 | 0.513 | 0.203 | -2.180 | 0.0530 |
| MAP4K3 | 8491 | 0.518 | 0.061 | -4.318 | 0.0046 |
| P101-PI3K | 23533 | 0.527 | 0.177 | -2.359 | 0.0435 |
| MATK | 4145 | 0.528 | 0.117 | -3.154 | 0.0194 |
| HUS1 | 3364 | 0.529 | 0.118 | -3.143 | 0.0196 |
| SENP7 | 57337 | 0.529 | 0.100 | -3.450 | 0.0140 |
| HNRPAB | 3182 | 0.530 | 0.064 | -4.155 | 0.0054 |
| HERC4 | 26091 | 0.530 | 0.119 | -3.116 | 0.0201 |
| CBX8 | 57332 | 0.530 | 0.119 | -3.105 | 0.0203 |
| CDKL4 | 344387 | 0.532 | 0.024 | -4.864 | 0.0002 |
| ACPT | 93650 | 0.546 | 0.090 | -3.498 | 0.0123 |
| RNF8 | 9025 | 0.551 | 0.074 | -3.774 | 0.0082 |
| siATRIP Control | siATRIP Contro | 0.557 | 0.114 | -3.001 | 0.0210 |
| CERS2 | 29956 | 0.560 | 0.064 | -3.881 | 0.0062 |
| LOC91461 | 91461 | 0.561 | 0.201 | -1.980 | 0.0631 |
| CA9 | 768 | 0.561 | 0.074 | -3.689 | 0.0085 |
| HDAC3 | 8841 | 0.562 | 0.141 | -2.596 | 0.0322 |
| ABL1 | 25 | 0.565 | 0.106 | -3.092 | 0.0184 |
| EZH2 | 2146 | 0.567 | 0.235 | -1.710 | 0.0858 |
| PAPOLA | 10914 | 0.567 | 0.131 | -2.689 | 0.0288 |
| HRMT1L3 | 10196 | 0.569 | 0.148 | -2.464 | 0.0367 |
| UHRF2 | 115426 | 0.577 | 0.235 | -1.673 | 0.0891 |

| | | | | | |
|---|---|---|---|---|---|
| CDK10 | 8558 | 0.589 | 0.103 | -2.964 | 0.0195 |
| SMUG1 | 23583 | 0.589 | 0.064 | -3.642 | 0.0070 |
| DDX26 | 26512 | 0.590 | 0.134 | -2.508 | 0.0335 |
| ANAPC10 | 10393 | 0.592 | 0.046 | -3.923 | 0.0032 |
| STK33 | 65975 | 0.600 | 0.071 | -3.410 | 0.0095 |
| ADAR | 103 | 0.606 | 0.083 | -3.160 | 0.0136 |
| CTDSP1 | 58190 | 0.606 | 0.075 | -3.295 | 0.0109 |
| OCRL | 4952 | 0.606 | 0.161 | -2.120 | 0.0508 |
| MGC10067 | 134510 | 0.607 | 0.267 | -1.392 | 0.1250 |
| FBXL20 | 84961 | 0.612 | 0.065 | -3.410 | 0.0083 |
| TGFBR1 | 7046 | 0.614 | 0.045 | -3.723 | 0.0034 |
| FN3K | 64122 | 0.616 | 0.018 | -4.040 | 0.0000 |
| SRPK1 | 6732 | 0.618 | 0.053 | -3.561 | 0.0053 |
| KIAA1333 | 55632 | 0.622 | 0.120 | -2.488 | 0.0314 |
| STK17A | 9263 | 0.626 | 0.155 | -2.072 | 0.0521 |
| PPP1R7 | 5510 | 0.627 | 0.113 | -2.548 | 0.0285 |
| EPM2A | 7957 | 0.628 | 0.165 | -1.968 | 0.0590 |
| ADK | 132 | 0.631 | 0.076 | -3.072 | 0.0128 |
| RIPK2 | 8767 | 0.631 | 0.118 | -2.448 | 0.0320 |
| RAD9A | 5883 | 0.632 | 0.107 | -2.596 | 0.0262 |
| UBE2T | 29089 | 0.642 | 0.238 | -1.400 | 0.1211 |
| PKM2 | 5315 | 0.644 | 0.187 | -1.704 | 0.0806 |
| SETDB2 | 83852 | 0.655 | 0.022 | -3.604 | 0.0003 |
| PUS3 | 83480 | 0.655 | 0.096 | -2.580 | 0.0239 |
| SENP1 | 29843 | 0.657 | 0.197 | -1.574 | 0.0943 |
| HELLS | 3070 | 0.657 | 0.089 | -2.661 | 0.0208 |
| EDF1 | 8721 | 0.664 | 0.167 | -1.756 | 0.0731 |
| HELZ | 9931 | 0.665 | 0.055 | -3.101 | 0.0075 |
| UCHL1 | 7345 | 0.667 | 0.197 | -1.530 | 0.0989 |
| RNASEH1 | 246243 | 0.669 | 0.107 | -2.332 | 0.0323 |
| UBE2L6 | 9246 | 0.673 | 0.152 | -1.833 | 0.0647 |
| TIF1 | 8805 | 0.673 | 0.146 | -1.890 | 0.0598 |
| UNG2 | 10309 | 0.675 | 0.039 | -3.213 | 0.0033 |
| UCHL5 | 51377 | 0.676 | 0.092 | -2.482 | 0.0246 |
| SARS | 6301 | 0.680 | 0.089 | -2.489 | 0.0237 |
| ERF | 2077 | 0.684 | 0.245 | -1.205 | 0.1550 |
| CTDP1 | 9150 | 0.684 | 0.110 | -2.199 | 0.0370 |
| UAP1 | 6675 | 0.684 | 0.133 | -1.948 | 0.0535 |
| SHPRH | 257218 | 0.685 | 0.223 | -1.299 | 0.1347 |
| URKL1 | 54963 | 0.686 | 0.098 | -2.329 | 0.0298 |
| FGR | 2268 | 0.688 | 0.040 | -3.077 | 0.0039 |
| DUSP16 | 80824 | 0.690 | 0.184 | -1.498 | 0.1004 |
| DYRK3 | 8444 | 0.694 | 0.056 | -2.808 | 0.0097 |

| | | | | | |
|---|---|---|---|---|---|
| NPEPPS | 9520 | 0.694 | 0.093 | -2.324 | 0.0284 |
| PRKCN | 23683 | 0.698 | 0.136 | -1.833 | 0.0607 |
| PLK2 | 10769 | 0.701 | 0.072 | -2.543 | 0.0174 |
| REV1L | 51455 | 0.701 | 0.089 | -2.320 | 0.0273 |
| ROCK1 | 6093 | 0.702 | 0.134 | -1.827 | 0.0606 |
| C20ORF13 | 55617 | 0.705 | 0.120 | -1.939 | 0.0505 |
| CRY2 | 1408 | 0.705 | 0.200 | -1.336 | 0.1250 |
| SGK494 | 124923 | 0.705 | 0.129 | -1.855 | 0.0574 |
| CHFR | 55743 | 0.706 | 0.060 | -2.647 | 0.0123 |
| SNARK | 81788 | 0.707 | 0.108 | -2.051 | 0.0418 |
| CDKN3 | 1033 | 0.710 | 0.169 | -1.501 | 0.0969 |
| PPP3CC | 5533 | 0.711 | 0.178 | -1.441 | 0.1059 |
| PRKWNK1 | 65125 | 0.712 | 0.198 | -1.316 | 0.1277 |
| STK32B | 55351 | 0.715 | 0.142 | -1.678 | 0.0732 |
| ERCC6 | 2074 | 0.718 | 0.291 | -0.923 | 0.2349 |
| MCM8 | 84515 | 0.721 | 0.165 | -1.472 | 0.0990 |
| C9ORF96 | 169436 | 0.722 | 0.074 | -2.331 | 0.0217 |
| UBE2E1 | 7324 | 0.722 | 0.046 | -2.674 | 0.0071 |
| FLJ35894 | 283847 | 0.725 | 0.102 | -1.988 | 0.0422 |
| FBXL3P | 26223 | 0.725 | 0.039 | -2.718 | 0.0047 |
| UBR2 | 23304 | 0.725 | 0.209 | -1.200 | 0.1505 |
| USP18 | 11274 | 0.726 | 0.297 | -0.879 | 0.2517 |
| IKBKG | 8517 | 0.727 | 0.172 | -1.398 | 0.1099 |
| G3BP2 | 9908 | 0.727 | 0.050 | -2.583 | 0.0091 |
| HRMT1L4 | 56341 | 0.728 | 0.106 | -1.928 | 0.0461 |
| UHMK1 | 127933 | 0.728 | 0.134 | -1.668 | 0.0714 |
| TREX1 | 11277 | 0.728 | 0.145 | -1.582 | 0.0820 |
| CHD4 | 1108 | 0.730 | 0.139 | -1.614 | 0.0774 |
| FBXL2 | 25827 | 0.732 | 0.267 | -0.950 | 0.2233 |
| FEM1B | 10116 | 0.733 | 0.096 | -2.003 | 0.0390 |
| SENP8 | 123228 | 0.734 | 0.130 | -1.667 | 0.0701 |
| HIF1AN | 55662 | 0.736 | 0.126 | -1.685 | 0.0673 |
| SENP6 | 26054 | 0.738 | 0.089 | -2.040 | 0.0348 |
| TAF11 | 6882 | 0.741 | 0.181 | -1.272 | 0.1312 |
| PIAS3 | 10401 | 0.743 | 0.091 | -1.978 | 0.0378 |
| TDP1 | 55775 | 0.743 | 0.411 | -0.609 | 0.3923 |
| MAPKAPK3 | 7867 | 0.743 | 0.065 | -2.255 | 0.0193 |
| PPM1E | 22843 | 0.744 | 0.366 | -0.677 | 0.3498 |
| USP42 | 84132 | 0.745 | 0.184 | -1.238 | 0.1378 |
| BRD2 | 6046 | 0.747 | 0.070 | -2.171 | 0.0230 |
| RIOK3 | 8780 | 0.748 | 0.104 | -1.807 | 0.0511 |
| PME-1 | 51400 | 0.750 | 0.078 | -2.053 | 0.0298 |
| SENP5 | 205564 | 0.751 | 0.141 | -1.478 | 0.0910 |

| | | | | | |
|---|---|---|---|---|---|
| BMP2K | 55589 | 0.753 | 0.089 | -1.920 | 0.0392 |
| APC | 324 | 0.754 | 0.063 | -2.187 | 0.0192 |
| RAD50 | 10111 | 0.754 | 0.244 | -0.939 | 0.2236 |
| NUCKS | 64710 | 0.755 | 0.135 | -1.489 | 0.0880 |
| USP13 | 8975 | 0.755 | 0.070 | -2.103 | 0.0243 |
| POLD1 | 5424 | 0.760 | 0.130 | -1.506 | 0.0841 |
| NEIL2 | 252969 | 0.760 | 0.070 | -2.061 | 0.0256 |
| GSK3A | 2931 | 0.761 | 0.051 | -2.253 | 0.0127 |
| REV3L | 5980 | 0.761 | 0.127 | -1.513 | 0.0824 |
| ZDHHC2 | 51201 | 0.764 | 0.127 | -1.498 | 0.0838 |
| CDK5 | 1020 | 0.765 | 0.023 | -2.446 | 0.0008 |
| MRE11A | 4361 | 0.766 | 0.061 | -2.108 | 0.0197 |
| TAF1L | 138474 | 0.768 | 0.170 | -1.195 | 0.1417 |
| CMAS | 55907 | 0.769 | 0.078 | -1.901 | 0.0345 |
| POLE3 | 54107 | 0.769 | 0.094 | -1.742 | 0.0501 |
| SIAH1 | 6477 | 0.773 | 0.147 | -1.307 | 0.1148 |
| TAF1 | 6872 | 0.773 | 0.101 | -1.648 | 0.0595 |
| FBXO22 | 26263 | 0.774 | 0.190 | -1.065 | 0.1759 |
| CDY1 | 9085 | 0.779 | 0.065 | -1.953 | 0.0252 |
| PLSCR2 | 57047 | 0.779 | 0.052 | -2.073 | 0.0154 |
| DDB2 | 1643 | 0.780 | 0.152 | -1.233 | 0.1287 |
| CLK2 | 1196 | 0.781 | 0.026 | -2.263 | 0.0019 |
| FKBP4 | 2288 | 0.782 | 0.086 | -1.717 | 0.0468 |
| CHD8 | 57680 | 0.783 | 0.034 | -2.188 | 0.0054 |
| TBL1XR1 | 79718 | 0.783 | 0.232 | -0.868 | 0.2465 |
| PIM1 | 5292 | 0.783 | 0.143 | -1.270 | 0.1192 |
| FN3KRP | 79672 | 0.783 | 0.040 | -2.136 | 0.0085 |
| CDY2A | 9426 | 0.786 | 0.036 | -2.144 | 0.0065 |
| PTPN18 | 26469 | 0.786 | 0.015 | -2.270 | 0.0000 |
| VRK2 | 7444 | 0.788 | 0.107 | -1.495 | 0.0742 |
| MYCBP2 | 23077 | 0.788 | 0.487 | -0.427 | 0.5299 |
| CHRAC1 | 54108 | 0.789 | 0.077 | -1.742 | 0.0403 |
| SMG1 | 23049 | 0.790 | 0.069 | -1.815 | 0.0321 |
| PRKAG1 | 5571 | 0.790 | 0.089 | -1.627 | 0.0537 |
| HAT1 | 8520 | 0.791 | 0.080 | -1.701 | 0.0442 |
| CCNK | 8812 | 0.791 | 0.200 | -0.947 | 0.2120 |
| SPAG5 | 10615 | 0.792 | 0.137 | -1.258 | 0.1184 |
| EXO1 | 9156 | 0.793 | 0.027 | -2.129 | 0.0027 |
| IRAK4 | 51135 | 0.794 | 0.129 | -1.291 | 0.1096 |
| CDK5R1 | 8851 | 0.796 | 0.077 | -1.692 | 0.0422 |
| UBE2A | 7319 | 0.796 | 0.115 | -1.378 | 0.0908 |
| BRD3 | 8019 | 0.798 | 0.036 | -2.029 | 0.0071 |
| DYRK4 | 8798 | 0.802 | 0.061 | -1.778 | 0.0276 |

| | | | | | |
|---|---|---|---|---|---|
| CHD3 | 1107 | 0.803 | 0.142 | -1.164 | 0.1367 |
| PPP3CB | 5532 | 0.803 | 0.125 | -1.259 | 0.1121 |
| NOP5/NOP58 | 51602 | 0.804 | 0.111 | -1.352 | 0.0912 |
| CTBP1 | 1487 | 0.807 | 0.075 | -1.614 | 0.0451 |
| ERCC2 | 2068 | 0.807 | 0.098 | -1.427 | 0.0749 |
| ATP5A1 | 498 | 0.807 | 0.132 | -1.190 | 0.1272 |
| CARM1 | 10498 | 0.809 | 0.075 | -1.599 | 0.0457 |
| ERCC5 | 2073 | 0.809 | 0.075 | -1.591 | 0.0465 |
| PCAF | 8850 | 0.811 | 0.087 | -1.486 | 0.0622 |
| FLJ32752 | 144132 | 0.815 | 0.151 | -1.040 | 0.1680 |
| INPP4B | 8821 | 0.817 | 0.044 | -1.771 | 0.0158 |
| TOP1 | 7150 | 0.817 | 0.038 | -1.817 | 0.0103 |
| RRM1 | 6240 | 0.818 | 0.225 | -0.747 | 0.2961 |
| MGMT | 4255 | 0.818 | 0.032 | -1.843 | 0.0067 |
| ADCK2 | 90956 | 0.819 | 0.125 | -1.159 | 0.1287 |
| MDM2 | 4193 | 0.821 | 0.307 | -0.559 | 0.4184 |
| USP11 | 8237 | 0.822 | 0.113 | -1.217 | 0.1107 |
| HEL308 | 113510 | 0.822 | 0.172 | -0.909 | 0.2148 |
| PIN1 | 5300 | 0.823 | 0.101 | -1.286 | 0.0930 |
| KIAA0377 | 9677 | 0.823 | 0.163 | -0.943 | 0.2003 |
| SPHK1 | 8877 | 0.825 | 0.219 | -0.736 | 0.3001 |
| MTMR6 | 9107 | 0.826 | 0.021 | -1.823 | 0.0013 |
| LOC340156 | 340156 | 0.826 | 0.095 | -1.308 | 0.0850 |
| PTPLA | 9200 | 0.827 | 0.086 | -1.362 | 0.0720 |
| CHD2 | 1106 | 0.828 | 0.032 | -1.744 | 0.0076 |
| USP40 | 55230 | 0.832 | 0.159 | -0.912 | 0.2084 |
| UBE2W | 55284 | 0.833 | 0.171 | -0.857 | 0.2327 |
| POLL | 27343 | 0.833 | 0.138 | -1.003 | 0.1703 |
| ATM | 472 | 0.834 | 0.126 | -1.057 | 0.1501 |
| XRCC2 | 7516 | 0.835 | 0.113 | -1.127 | 0.1265 |
| LTK | 4058 | 0.837 | 0.122 | -1.060 | 0.1465 |
| FLJ22405 | 64419 | 0.839 | 0.124 | -1.039 | 0.1523 |
| APEX1 | 328 | 0.846 | 0.050 | -1.454 | 0.0303 |
| C18orf56 | 494514 | 0.848 | 0.050 | -1.444 | 0.0299 |
| SETDB1 | 9869 | 0.849 | 0.173 | -0.770 | 0.2687 |
| PTEN | 5728 | 0.849 | 0.063 | -1.342 | 0.0509 |
| PPP5C | 5536 | 0.852 | 0.103 | -1.069 | 0.1284 |
| PPFIA1 | 8500 | 0.853 | 0.107 | -1.033 | 0.1404 |
| ELAC1 | 55520 | 0.853 | 0.146 | -0.846 | 0.2238 |
| ATP6V0A2 | 23545 | 0.855 | 0.094 | -1.100 | 0.1137 |
| GSG2 | 83903 | 0.856 | 0.103 | -1.038 | 0.1352 |
| MPHOSPH1 | 9585 | 0.859 | 0.144 | -0.819 | 0.2327 |
| POLDIP3 | 84271 | 0.860 | 0.090 | -1.078 | 0.1139 |

| | | | | | |
|---|---|---|---|---|---|
| HIPK3 | 10114 | 0.860 | 0.123 | -0.906 | 0.1873 |
| PPM1D | 8493 | 0.861 | 0.016 | -1.474 | 0.0005 |
| HDAC9 | 9734 | 0.861 | 0.105 | -0.986 | 0.1491 |
| RFC5 | 5985 | 0.862 | 0.213 | -0.594 | 0.3778 |
| RECQL | 5965 | 0.863 | 0.168 | -0.716 | 0.2915 |
| NUDT15 | 55270 | 0.863 | 0.164 | -0.726 | 0.2847 |
| SET7 | 80854 | 0.865 | 0.115 | -0.912 | 0.1787 |
| NCOA3 | 8202 | 0.865 | 0.104 | -0.962 | 0.1542 |
| AASDHPPT | 60496 | 0.867 | 0.046 | -1.279 | 0.0339 |
| UBE2Q1 | 55585 | 0.867 | 0.256 | -0.488 | 0.4631 |
| MAST4 | 375449 | 0.870 | 0.041 | -1.278 | 0.0263 |
| FEN1 | 2237 | 0.872 | 0.169 | -0.662 | 0.3206 |
| MK-STYX | 51657 | 0.872 | 0.083 | -1.025 | 0.1147 |
| CDY2B | 203611 | 0.874 | 0.084 | -1.005 | 0.1202 |
| MMP9 | 4318 | 0.875 | 0.002 | -1.337 | 0.0000 |
| SUV420H1 | 51111 | 0.877 | 0.327 | -0.363 | 0.5808 |
| Non-Transfect Co | Non-Transfect ( | 0.879 | 0.030 | -1.235 | 0.0130 |
| KUB3 | 91419 | 0.879 | 0.079 | -0.987 | 0.1162 |
| TAF15 | 8148 | 0.880 | 0.186 | -0.578 | 0.3792 |
| PSKH2 | 85481 | 0.880 | 0.032 | -1.218 | 0.0161 |
| CDC25C | 995 | 0.880 | 0.130 | -0.747 | 0.2523 |
| LAK | 80216 | 0.881 | 0.216 | -0.505 | 0.4413 |
| UBE2D4 | 51619 | 0.887 | 0.050 | -1.068 | 0.0551 |
| PPM1A | 5494 | 0.887 | 0.158 | -0.612 | 0.3435 |
| FLJ25449 | 151649 | 0.887 | 0.069 | -0.973 | 0.1021 |
| UFD1L | 7353 | 0.888 | 0.113 | -0.763 | 0.2282 |
| USP19 | 10869 | 0.889 | 0.150 | -0.630 | 0.3273 |
| RNF148 | 378925 | 0.890 | 0.230 | -0.445 | 0.4931 |
| FLJ13052 | 65220 | 0.890 | 0.101 | -0.799 | 0.1998 |
| PCOLN3 | 5119 | 0.893 | 0.037 | -1.068 | 0.0311 |
| BRIP1 | 83990 | 0.894 | 0.138 | -0.637 | 0.3141 |
| FBXO32 | 114907 | 0.896 | 0.167 | -0.542 | 0.3949 |
| EYA3 | 2140 | 0.896 | 0.046 | -0.995 | 0.0552 |
| TAF1C | 9013 | 0.897 | 0.065 | -0.904 | 0.1083 |
| LASS6 | 253782 | 0.898 | 0.085 | -0.811 | 0.1710 |
| DNMT2 | 1787 | 0.900 | 0.191 | -0.469 | 0.4617 |
| PPP2R5D | 5528 | 0.901 | 0.037 | -0.987 | 0.0363 |
| ASK | 10926 | 0.902 | 0.076 | -0.814 | 0.1533 |
| ENC1 | 8507 | 0.905 | 0.172 | -0.487 | 0.4388 |
| MSH2 | 4436 | 0.907 | 0.108 | -0.652 | 0.2744 |
| UBE2R2 | 54926 | 0.907 | 0.080 | -0.755 | 0.1813 |
| CUL5 | 8065 | 0.908 | 0.141 | -0.541 | 0.3785 |
| POP4 | 10775 | 0.910 | 0.127 | -0.573 | 0.3439 |

| | | | | | |
|---|---|---|---|---|---|
| UBE3A | 7337 | 0.911 | 0.134 | -0.545 | 0.3691 |
| TP53RK | 112858 | 0.912 | 0.124 | -0.570 | 0.3425 |
| PLK3 | 1263 | 0.912 | 0.231 | -0.355 | 0.5760 |
| USP27X | 389856 | 0.914 | 0.141 | -0.507 | 0.4026 |
| TP53 | 7157 | 0.915 | 0.274 | -0.295 | 0.6432 |
| CBLL1 | 79872 | 0.915 | 0.179 | -0.419 | 0.4990 |
| PSEN2 | 5664 | 0.916 | 0.063 | -0.750 | 0.1441 |
| CDC14A | 8556 | 0.917 | 0.079 | -0.683 | 0.2073 |
| FBXL5 | 26234 | 0.917 | 0.209 | -0.365 | 0.5608 |
| OSR1 | 9943 | 0.917 | 0.197 | -0.381 | 0.5414 |
| INPP5E | 56623 | 0.918 | 0.102 | -0.594 | 0.2978 |
| FLJ11200 | 55325 | 0.922 | 0.179 | -0.385 | 0.5310 |
| UBE2H | 7328 | 0.922 | 0.281 | -0.262 | 0.6794 |
| UBE2I | 7329 | 0.924 | 0.226 | -0.313 | 0.6174 |
| INOC1 | 54617 | 0.924 | 0.147 | -0.438 | 0.4641 |
| ATR | 545 | 0.924 | 0.171 | -0.391 | 0.5211 |
| SPO11 | 23626 | 0.925 | 0.137 | -0.451 | 0.4451 |
| CYLD | 1540 | 0.926 | 0.093 | -0.567 | 0.2977 |
| MAD2L2 | 10459 | 0.927 | 0.063 | -0.649 | 0.1808 |
| STAMBP | 10617 | 0.928 | 0.101 | -0.526 | 0.3405 |
| LIG1 | 3978 | 0.929 | 0.059 | -0.646 | 0.1701 |
| PRKAG2 | 51422 | 0.930 | 0.138 | -0.423 | 0.4706 |
| ADARB2 | 105 | 0.930 | 0.027 | -0.722 | 0.0347 |
| SIRT7 | 51547 | 0.931 | 0.144 | -0.401 | 0.4950 |
| CASP8 | 841 | 0.931 | 0.023 | -0.713 | 0.0228 |
| PBEF1 | 10135 | 0.932 | 0.160 | -0.367 | 0.5385 |
| SMARCAD1 | 56916 | 0.933 | 0.169 | -0.346 | 0.5643 |
| MYST3 | 7994 | 0.934 | 0.126 | -0.423 | 0.4591 |
| RAD51C | 5889 | 0.936 | 0.148 | -0.363 | 0.5349 |
| PPP2R2D | 55844 | 0.937 | 0.082 | -0.507 | 0.3145 |
| IPO8 | 10526 | 0.939 | 0.147 | -0.349 | 0.5490 |
| ANKIB1 | 54467 | 0.940 | 0.009 | -0.640 | 0.0001 |
| RAD51 | 5888 | 0.942 | 0.230 | -0.235 | 0.7042 |
| CHAT | 1103 | 0.944 | 0.054 | -0.524 | 0.2114 |
| NLK | 51701 | 0.947 | 0.143 | -0.312 | 0.5847 |
| POLN | 353497 | 0.947 | 0.104 | -0.381 | 0.4697 |
| TAZ | 6901 | 0.948 | 0.097 | -0.389 | 0.4492 |
| PPP2CZ | 333926 | 0.949 | 0.072 | -0.432 | 0.3440 |
| LPIN3 | 64900 | 0.950 | 0.061 | -0.448 | 0.2901 |
| DTYMK | 1841 | 0.951 | 0.059 | -0.443 | 0.2852 |
| DBR1 | 51163 | 0.952 | 0.131 | -0.299 | 0.5898 |
| BAP1 | 8314 | 0.953 | 0.138 | -0.284 | 0.6137 |
| MAST3 | 23031 | 0.953 | 0.156 | -0.260 | 0.6523 |

| | | | | | |
|---|---|---|---|---|---|
| SMARCE1 | 6605 | 0.957 | 0.164 | -0.225 | 0.6978 |
| RAD54B | 25788 | 0.958 | 0.150 | -0.240 | 0.6732 |
| RTCD1 | 8634 | 0.958 | 0.015 | -0.444 | 0.0147 |
| FLJ23074 | 80122 | 0.959 | 0.022 | -0.430 | 0.0673 |
| ROCK2 | 9475 | 0.959 | 0.334 | -0.119 | 0.8510 |
| USP44 | 84101 | 0.959 | 0.040 | -0.404 | 0.2136 |
| FBXO11 | 80204 | 0.961 | 0.107 | -0.273 | 0.5947 |
| DLG7 | 9787 | 0.962 | 0.251 | -0.142 | 0.8175 |
| RNF2 | 6045 | 0.965 | 0.203 | -0.155 | 0.7957 |
| CDY1B | 253175 | 0.966 | 0.169 | -0.175 | 0.7629 |
| HDAC5 | 10014 | 0.967 | 0.023 | -0.348 | 0.2622 |
| NCOA1 | 8648 | 0.967 | 0.024 | -0.344 | 0.1230 |
| MUTYH | 4595 | 0.968 | 0.082 | -0.256 | 0.5729 |
| AEBP1 | 165 | 0.969 | 0.075 | -0.260 | 0.5495 |
| CTH | 1491 | 0.969 | 0.039 | -0.303 | 0.3080 |
| THNSL1 | 79896 | 0.970 | 0.106 | -0.212 | 0.6740 |
| ACF | 29974 | 0.971 | 0.103 | -0.212 | 0.6700 |
| MYST2 | 11143 | 0.971 | 0.350 | -0.080 | 0.8995 |
| CLK3 | 1198 | 0.971 | 0.057 | -0.263 | 0.4780 |
| SUV39H2 | 79723 | 0.971 | 0.135 | -0.174 | 0.7490 |
| ASNA1 | 439 | 0.974 | 0.093 | -0.200 | 0.6720 |
| UMP-CMPK | 51727 | 0.975 | 0.182 | -0.121 | 0.8359 |
| KIAA1804 | 84451 | 0.976 | 0.108 | -0.169 | 0.7369 |
| USP9Y | 8287 | 0.976 | 0.140 | -0.141 | 0.7976 |
| STK24 | 8428 | 0.977 | 0.145 | -0.132 | 0.8112 |
| SHFM1 | 7979 | 0.977 | 0.150 | -0.128 | 0.8193 |
| USP25 | 29761 | 0.979 | 0.270 | -0.075 | 0.9030 |
| DCK | 1633 | 0.979 | 0.312 | -0.063 | 0.9197 |
| MGC42105 | 167359 | 0.980 | 0.199 | -0.092 | 0.8760 |
| PRKDC | 5591 | 0.981 | 0.132 | -0.117 | 0.8274 |
| ARL4A | 10124 | 0.983 | 0.122 | -0.111 | 0.8315 |
| UBE2O | 63893 | 0.984 | 0.050 | -0.154 | 0.6315 |
| RUVBL1 | 8607 | 0.985 | 0.207 | -0.068 | 0.9098 |
| SAE1 | 10055 | 0.985 | 0.127 | -0.095 | 0.8580 |
| NTAN1 | 123803 | 0.986 | 0.150 | -0.081 | 0.8837 |
| UBE2B | 7320 | 0.987 | 0.071 | -0.113 | 0.7782 |
| ATP2A3 | 489 | 0.988 | 0.089 | -0.093 | 0.8376 |
| USP3 | 9960 | 0.988 | 0.217 | -0.049 | 0.9346 |
| NSD1 | 64324 | 0.989 | 0.054 | -0.100 | 0.7661 |
| SUMO2P7 | 100131823 | 0.992 | 0.139 | -0.048 | 0.9296 |
| PRKWNK2 | 65268 | 0.992 | 0.068 | -0.065 | 0.8667 |
| NAT8 | 9027 | 0.996 | 0.186 | -0.019 | 0.9742 |
| CHEK2 | 11200 | 0.996 | 0.149 | -0.021 | 0.9701 |

| | | | | | |
|---|---|---|---|---|---|
| FBXO25 | 26260 | 0.998 | 0.043 | -0.024 | 0.9320 |
| TESK2 | 10420 | 0.998 | 0.229 | -0.008 | 0.9896 |
| VRK1 | 7443 | 0.999 | 0.185 | -0.004 | 0.9943 |
| siNS Control | siNS Control | 1.000 | 0.093 | 0.000 | 1.0000 |
| PTGS1 | 5742 | 1.000 | 0.105 | 0.001 | 0.9990 |
| PIAS2 | 9063 | 1.002 | 0.147 | 0.009 | 0.9874 |
| STK17B | 9262 | 1.004 | 0.191 | 0.020 | 0.9723 |
| PRDM9 | 56979 | 1.005 | 0.132 | 0.030 | 0.9553 |
| PPEF1 | 5475 | 1.006 | 0.071 | 0.049 | 0.9027 |
| NEK3 | 4752 | 1.008 | 0.201 | 0.034 | 0.9536 |
| LHPP | 64077 | 1.010 | 0.020 | 0.104 | 0.4890 |
| BCOR | 54880 | 1.010 | 0.064 | 0.092 | 0.8048 |
| DUSP14 | 11072 | 1.011 | 0.147 | 0.061 | 0.9125 |
| NXN | 64359 | 1.011 | 0.184 | 0.054 | 0.9267 |
| POLH | 5429 | 1.011 | 0.054 | 0.105 | 0.7526 |
| DUSP1 | 1843 | 1.012 | 0.143 | 0.069 | 0.8995 |
| SMARCA1 | 6594 | 1.013 | 0.050 | 0.120 | 0.7063 |
| LATS2 | 26524 | 1.013 | 0.029 | 0.133 | 0.5279 |
| TNKS | 8658 | 1.015 | 0.233 | 0.062 | 0.9190 |
| XRCC5 | 7520 | 1.019 | 0.112 | 0.127 | 0.8016 |
| NDRG1 | 10397 | 1.022 | 0.119 | 0.145 | 0.7811 |
| USP54 | 159195 | 1.025 | 0.085 | 0.196 | 0.6639 |
| YPEL1 | 29799 | 1.026 | 0.163 | 0.138 | 0.8088 |
| MSRB | 22921 | 1.026 | 0.055 | 0.244 | 0.4942 |
| NME7 | 29922 | 1.028 | 0.110 | 0.194 | 0.7032 |
| CDADC1 | 81602 | 1.029 | 0.063 | 0.256 | 0.5114 |
| FMR1 | 2332 | 1.030 | 0.126 | 0.193 | 0.7183 |
| STK19 | 8859 | 1.032 | 0.128 | 0.205 | 0.7036 |
| NEIL3 | 55247 | 1.033 | 0.138 | 0.196 | 0.7218 |
| PPP2R3A | 5523 | 1.033 | 0.095 | 0.248 | 0.6088 |
| TRIM33 | 51592 | 1.033 | 0.209 | 0.144 | 0.8105 |
| PRPS2 | 5634 | 1.034 | 0.097 | 0.252 | 0.6079 |
| UBE3C | 9690 | 1.034 | 0.130 | 0.214 | 0.6932 |
| ENPP7 | 339221 | 1.037 | 0.062 | 0.326 | 0.4143 |
| RNF6 | 6049 | 1.038 | 0.106 | 0.266 | 0.6022 |
| KHSRP | 8570 | 1.039 | 0.077 | 0.324 | 0.4723 |
| PARP4 | 143 | 1.040 | 0.130 | 0.247 | 0.6511 |
| CCRK | 23552 | 1.041 | 0.094 | 0.307 | 0.5328 |
| UBR1 | 197131 | 1.041 | 0.118 | 0.276 | 0.6051 |
| USP2 | 9099 | 1.042 | 0.256 | 0.154 | 0.8038 |
| CDK7 | 1022 | 1.042 | 0.079 | 0.346 | 0.4519 |
| FBXO21 | 23014 | 1.045 | 0.074 | 0.377 | 0.4058 |
| POLE2 | 5427 | 1.045 | 0.136 | 0.275 | 0.6216 |

| | | | | | |
|---|---|---|---|---|---|
| CBLC | 23624 | 1.046 | 0.163 | 0.245 | 0.6737 |
| PRIMA1 | 145270 | 1.046 | 0.104 | 0.330 | 0.5239 |
| Mock Control | Mock Control | 1.047 | 0.021 | 0.493 | 0.0410 |
| RNF12 | 51132 | 1.048 | 0.104 | 0.344 | 0.5087 |
| USP15 | 9958 | 1.050 | 0.118 | 0.334 | 0.5388 |
| FBXO4 | 26272 | 1.051 | 0.183 | 0.246 | 0.6801 |
| ISG20L2 | 81875 | 1.051 | 0.060 | 0.462 | 0.2740 |
| JIK | 51347 | 1.052 | 0.219 | 0.220 | 0.7190 |
| PTPN11 | 5781 | 1.053 | 0.142 | 0.312 | 0.5846 |
| UBA2 | 10054 | 1.053 | 0.054 | 0.497 | 0.2246 |
| STK29 | 9024 | 1.054 | 0.161 | 0.289 | 0.6219 |
| HTCD37 | 58497 | 1.056 | 0.096 | 0.414 | 0.4233 |
| FLJ10458 | 54475 | 1.058 | 0.154 | 0.320 | 0.5841 |
| UCHL3 | 7347 | 1.058 | 0.266 | 0.205 | 0.7429 |
| UBE3B | 89910 | 1.058 | 0.227 | 0.238 | 0.6999 |
| AFMID | 125061 | 1.059 | 0.089 | 0.461 | 0.3663 |
| CDKL1 | 8814 | 1.060 | 0.107 | 0.422 | 0.4348 |
| NTRK2 | 4915 | 1.060 | 0.013 | 0.638 | 0.0023 |
| PPM1G | 5496 | 1.061 | 0.036 | 0.608 | 0.0920 |
| MUS81 | 80198 | 1.063 | 0.152 | 0.351 | 0.5501 |
| WRNIP1 | 56897 | 1.064 | 0.164 | 0.337 | 0.5708 |
| WRN | 7486 | 1.065 | 0.245 | 0.247 | 0.6922 |
| PPP4R1 | 9989 | 1.065 | 0.091 | 0.497 | 0.3437 |
| G3BP | 10146 | 1.065 | 0.008 | 0.696 | 0.0000 |
| MCM6 | 4175 | 1.065 | 0.187 | 0.312 | 0.6076 |
| ERBB2IP | 55914 | 1.067 | 0.202 | 0.301 | 0.6236 |
| PPIH | 10465 | 1.067 | 0.135 | 0.409 | 0.4796 |
| PAPOLG | 64895 | 1.069 | 0.306 | 0.215 | 0.7345 |
| AKT3 | 10000 | 1.070 | 0.150 | 0.394 | 0.5061 |
| PR48 | 28227 | 1.071 | 0.045 | 0.686 | 0.1050 |
| CDC23 | 8697 | 1.071 | 0.073 | 0.602 | 0.2324 |
| FLJ23751 | 92370 | 1.072 | 0.114 | 0.489 | 0.3875 |
| RECQL4 | 9401 | 1.074 | 0.161 | 0.398 | 0.5092 |
| GMPS | 8833 | 1.075 | 0.094 | 0.563 | 0.3029 |
| USP32 | 84669 | 1.077 | 0.131 | 0.478 | 0.4164 |
| MBD4 | 8930 | 1.077 | 0.099 | 0.571 | 0.3061 |
| NEK5 | 341676 | 1.078 | 0.092 | 0.592 | 0.2824 |
| AK7 | 122481 | 1.083 | 0.196 | 0.381 | 0.5411 |
| HNF4A | 3172 | 1.083 | 0.113 | 0.566 | 0.3322 |
| TESK1 | 7016 | 1.085 | 0.270 | 0.298 | 0.6395 |
| UBE2E3 | 10477 | 1.086 | 0.129 | 0.541 | 0.3669 |
| SUV420H2 | 84787 | 1.087 | 0.350 | 0.240 | 0.7088 |
| HECTD1 | 25831 | 1.087 | 0.049 | 0.826 | 0.0877 |

| | | | | | |
|---|---|---|---|---|---|
| NEK11 | 79858 | 1.088 | 0.180 | 0.433 | 0.4876 |
| TLE2 | 7089 | 1.088 | 0.157 | 0.484 | 0.4330 |
| NR2C1 | 7181 | 1.089 | 0.159 | 0.483 | 0.4345 |
| MYST4 | 23522 | 1.089 | 0.212 | 0.384 | 0.5433 |
| DNMT3B | 1789 | 1.090 | 0.188 | 0.430 | 0.4932 |
| PARP1 | 142 | 1.091 | 0.059 | 0.821 | 0.1136 |
| POLE | 5426 | 1.091 | 0.142 | 0.536 | 0.3825 |
| USP7 | 7874 | 1.092 | 0.089 | 0.717 | 0.2135 |
| PRKAB2 | 5565 | 1.093 | 0.168 | 0.486 | 0.4379 |
| MOV10L1 | 54456 | 1.094 | 0.332 | 0.272 | 0.6733 |
| RPS6KL1 | 83694 | 1.095 | 0.259 | 0.344 | 0.5917 |
| LENG5 | 79042 | 1.095 | 0.126 | 0.606 | 0.3219 |
| SMARCA2 | 6595 | 1.098 | 0.121 | 0.644 | 0.2939 |
| LEPREL2 | 10536 | 1.099 | 0.111 | 0.686 | 0.2600 |
| CAMK4 | 814 | 1.100 | 0.075 | 0.836 | 0.1451 |
| UBE2U | 148581 | 1.101 | 0.211 | 0.437 | 0.4948 |
| TENR | 132612 | 1.102 | 0.124 | 0.657 | 0.2900 |
| FLJ31952 | 146857 | 1.102 | 0.161 | 0.550 | 0.3862 |
| PSKH1 | 5681 | 1.106 | 0.120 | 0.693 | 0.2677 |
| CRSP9 | 9443 | 1.107 | 0.700 | 0.151 | 0.8161 |
| RTEL1 | 51750 | 1.109 | 0.025 | 1.125 | 0.0100 |
| NEK1 | 4750 | 1.110 | 0.261 | 0.397 | 0.5414 |
| UBE2G1 | 7326 | 1.111 | 0.157 | 0.607 | 0.3459 |
| SLFN5 | 342615 | 1.111 | 0.191 | 0.522 | 0.4201 |
| AURKB | 9212 | 1.113 | 0.063 | 1.004 | 0.0872 |
| HERC6 | 55008 | 1.113 | 0.135 | 0.691 | 0.2827 |
| FBXO5 | 26271 | 1.114 | 0.098 | 0.838 | 0.1824 |
| ADARB1 | 104 | 1.114 | 0.187 | 0.547 | 0.4007 |
| FLT1 | 2321 | 1.118 | 0.064 | 1.044 | 0.0823 |
| MASTL | 84930 | 1.119 | 0.084 | 0.946 | 0.1330 |
| DNTT | 1791 | 1.120 | 0.236 | 0.472 | 0.4722 |
| DUSP9 | 1852 | 1.121 | 0.225 | 0.494 | 0.4522 |
| HAK | 115701 | 1.121 | 0.090 | 0.936 | 0.1429 |
| ARK5 | 9891 | 1.122 | 0.082 | 0.978 | 0.1230 |
| DLP | 54957 | 1.122 | 0.131 | 0.758 | 0.2486 |
| ERCC4 | 2072 | 1.122 | 0.227 | 0.498 | 0.4494 |
| RERG | 85004 | 1.123 | 0.087 | 0.960 | 0.1342 |
| PARN | 5073 | 1.125 | 0.031 | 1.268 | 0.0142 |
| PIM2 | 11040 | 1.125 | 0.237 | 0.491 | 0.4574 |
| CKN1 | 1161 | 1.125 | 0.365 | 0.332 | 0.6127 |
| UBE1 | 7317 | 1.126 | 0.131 | 0.783 | 0.2374 |
| DPYSL4 | 10570 | 1.127 | 0.090 | 0.980 | 0.1335 |
| FLJ23356 | 84197 | 1.128 | 0.078 | 1.054 | 0.1027 |

| | | | | | |
|---|---|---|---|---|---|
| AOF2 | 23028 | 1.129 | 0.164 | 0.681 | 0.3078 |
| JAK1 | 3716 | 1.129 | 0.085 | 1.020 | 0.1184 |
| SRPK2 | 6733 | 1.130 | 0.227 | 0.529 | 0.4261 |
| SOD2 | 6648 | 1.130 | 0.122 | 0.848 | 0.2056 |
| USP28 | 57646 | 1.130 | 0.150 | 0.738 | 0.2711 |
| RNF130 | 55819 | 1.131 | 0.159 | 0.714 | 0.2877 |
| RP6-213H19.1 | 51765 | 1.132 | 0.151 | 0.746 | 0.2681 |
| TEP1 | 7011 | 1.133 | 0.147 | 0.765 | 0.2565 |
| FBXO3 | 26273 | 1.133 | 0.121 | 0.873 | 0.1960 |
| ZCCHC11 | 23318 | 1.133 | 0.027 | 1.374 | 0.0080 |
| SRP72 | 6731 | 1.134 | 0.194 | 0.621 | 0.3554 |
| ANKRD3 | 54101 | 1.136 | 0.113 | 0.929 | 0.1713 |
| RAD54L | 8438 | 1.137 | 0.082 | 1.099 | 0.1009 |
| FNBP1 | 23048 | 1.137 | 0.113 | 0.938 | 0.1689 |
| MCM7 | 4176 | 1.137 | 0.151 | 0.774 | 0.2557 |
| RAG1 | 5896 | 1.138 | 0.039 | 1.370 | 0.0208 |
| POLD4 | 57804 | 1.142 | 0.226 | 0.582 | 0.3891 |
| NME2 | 4831 | 1.143 | 0.214 | 0.616 | 0.3647 |
| MLCK | 91807 | 1.146 | 0.041 | 1.435 | 0.0204 |
| TTF2 | 8458 | 1.147 | 0.049 | 1.399 | 0.0309 |
| NEIL1 | 79661 | 1.147 | 0.223 | 0.609 | 0.3713 |
| MET | 4233 | 1.148 | 0.191 | 0.696 | 0.3117 |
| FLJ34389 | 197259 | 1.149 | 0.158 | 0.813 | 0.2434 |
| UBE4A | 9354 | 1.153 | 0.096 | 1.148 | 0.1078 |
| PDIK1L | 149420 | 1.154 | 0.076 | 1.281 | 0.0701 |
| RRM2B | 50484 | 1.154 | 0.051 | 1.452 | 0.0311 |
| PARK2 | 5071 | 1.155 | 0.054 | 1.438 | 0.0349 |
| KDR | 3791 | 1.155 | 0.108 | 1.090 | 0.1290 |
| SMARCA4 | 6597 | 1.155 | 0.114 | 1.057 | 0.1404 |
| EIF2AK4 | 440275 | 1.158 | 0.052 | 1.481 | 0.0311 |
| CHKB | 1120 | 1.159 | 0.128 | 1.007 | 0.1629 |
| PARP2 | 10038 | 1.161 | 0.105 | 1.143 | 0.1168 |
| FBXL3A | 26224 | 1.161 | 0.094 | 1.218 | 0.0955 |
| MSH6 | 2956 | 1.161 | 0.172 | 0.826 | 0.2449 |
| AURKC | 6795 | 1.162 | 0.130 | 1.016 | 0.1620 |
| PTPN20A | 653129 | 1.164 | 0.201 | 0.742 | 0.2919 |
| COIL | 8161 | 1.165 | 0.082 | 1.334 | 0.0710 |
| PPIE | 10450 | 1.166 | 0.081 | 1.344 | 0.0689 |
| TAF6L | 10629 | 1.166 | 0.179 | 0.826 | 0.2475 |
| ERCC1 | 2067 | 1.166 | 0.184 | 0.806 | 0.2582 |
| RPS6KA5 | 9252 | 1.170 | 0.542 | 0.309 | 0.6413 |
| USP26 | 83844 | 1.173 | 0.098 | 1.281 | 0.0908 |
| CDK4 | 1019 | 1.174 | 0.038 | 1.724 | 0.0120 |

| | | | | | |
|---|---|---|---|---|---|
| USP46 | 64854 | 1.174 | 0.044 | 1.689 | 0.0170 |
| UBE2Q2 | 92912 | 1.177 | 0.117 | 1.183 | 0.1191 |
| RNASEH2A | 10535 | 1.178 | 0.071 | 1.516 | 0.0473 |
| CBX4 | 8535 | 1.178 | 0.072 | 1.511 | 0.0485 |
| POP5 | 51367 | 1.178 | 0.081 | 1.441 | 0.0611 |
| PPP6C | 5537 | 1.181 | 0.192 | 0.849 | 0.2432 |
| NME1 | 4830 | 1.182 | 0.154 | 1.009 | 0.1774 |
| PPM1F | 9647 | 1.183 | 0.067 | 1.596 | 0.0396 |
| XYLB | 9942 | 1.184 | 0.183 | 0.897 | 0.2229 |
| POLM | 27434 | 1.184 | 0.090 | 1.421 | 0.0700 |
| FBXO7 | 25793 | 1.185 | 0.186 | 0.889 | 0.2268 |
| TNRC11 | 9968 | 1.186 | 0.136 | 1.128 | 0.1410 |
| SAMHD1 | 25939 | 1.188 | 0.154 | 1.043 | 0.1686 |
| NEK9 | 91754 | 1.188 | 0.119 | 1.248 | 0.1100 |
| UBE2L3 | 7332 | 1.189 | 0.211 | 0.818 | 0.2612 |
| NEK8 | 284086 | 1.191 | 0.245 | 0.729 | 0.3093 |
| UBOX5 | 22888 | 1.191 | 0.166 | 1.007 | 0.1828 |
| NMNAT1 | 64802 | 1.196 | 0.544 | 0.354 | 0.5969 |
| NYD-SP25 | 89882 | 1.196 | 0.109 | 1.373 | 0.0875 |
| ELP3 | 55140 | 1.197 | 0.124 | 1.266 | 0.1106 |
| PSMC5 | 5705 | 1.197 | 0.386 | 0.498 | 0.4688 |
| MYO3B | 140469 | 1.200 | 0.148 | 1.145 | 0.1431 |
| PPP2R1B | 5519 | 1.200 | 0.026 | 2.067 | 0.0024 |
| CDC25B | 994 | 1.202 | 0.093 | 1.539 | 0.0620 |
| CASP3 | 836 | 1.206 | 0.044 | 1.997 | 0.0124 |
| TDG | 6996 | 1.207 | 0.052 | 1.938 | 0.0178 |
| UBE2N | 7334 | 1.207 | 0.107 | 1.464 | 0.0770 |
| LPIN1 | 23175 | 1.207 | 0.102 | 1.500 | 0.0710 |
| RBBP4 | 5928 | 1.208 | 0.072 | 1.759 | 0.0365 |
| CCNB1IP1 | 57820 | 1.208 | 0.132 | 1.287 | 0.1113 |
| CENTG1 | 116986 | 1.210 | 0.130 | 1.312 | 0.1068 |
| FLJ20321 | 54897 | 1.210 | 0.130 | 1.312 | 0.1070 |
| USP4 | 7375 | 1.211 | 0.101 | 1.541 | 0.0668 |
| USP5 | 8078 | 1.212 | 0.120 | 1.397 | 0.0913 |
| FLJ32685 | 152110 | 1.216 | 0.064 | 1.910 | 0.0261 |
| DKFZP434C212 | 26130 | 1.216 | 0.018 | 2.279 | 0.0002 |
| RIOK1 | 83732 | 1.217 | 0.126 | 1.383 | 0.0959 |
| RNGTT | 8732 | 1.220 | 0.103 | 1.590 | 0.0641 |
| PPAPDC1 | 196051 | 1.222 | 0.222 | 0.921 | 0.2256 |
| PPP2R5C | 5527 | 1.222 | 0.067 | 1.938 | 0.0271 |
| ZNF650 | 130507 | 1.223 | 0.107 | 1.565 | 0.0685 |
| USP29 | 57663 | 1.223 | 0.125 | 1.432 | 0.0897 |
| TRIB3 | 57761 | 1.225 | 0.339 | 0.639 | 0.3700 |

| | | | | | |
|---|---|---|---|---|---|
| SPAST | 6683 | 1.226 | 0.250 | 0.846 | 0.2584 |
| KIAA0073 | 23398 | 1.227 | 0.141 | 1.339 | 0.1082 |
| MAP4K4 | 9448 | 1.227 | 0.095 | 1.704 | 0.0527 |
| NEK6 | 10783 | 1.229 | 0.082 | 1.839 | 0.0391 |
| FLJ10761 | 55224 | 1.229 | 0.417 | 0.536 | 0.4415 |
| FBXW2 | 26190 | 1.229 | 0.094 | 1.732 | 0.0505 |
| ESPL1 | 9700 | 1.229 | 0.509 | 0.444 | 0.5163 |
| USP39 | 10713 | 1.232 | 0.215 | 0.990 | 0.2022 |
| USP21 | 27005 | 1.233 | 0.200 | 1.059 | 0.1801 |
| PPP2R2C | 5522 | 1.234 | 0.089 | 1.814 | 0.0439 |
| RFC3 | 5983 | 1.235 | 0.080 | 1.913 | 0.0350 |
| CUL2 | 8453 | 1.235 | 0.497 | 0.465 | 0.4989 |
| C14ORF20 | 283629 | 1.236 | 0.122 | 1.539 | 0.0776 |
| POLQ | 10721 | 1.236 | 0.121 | 1.547 | 0.0766 |
| PRKACG | 5568 | 1.238 | 0.280 | 0.805 | 0.2795 |
| HDAC2 | 3066 | 1.238 | 0.141 | 1.408 | 0.0993 |
| ACIN1 | 22985 | 1.238 | 0.176 | 1.197 | 0.1432 |
| PPFIBP2 | 8495 | 1.240 | 0.127 | 1.529 | 0.0806 |
| PSPHL | 8781 | 1.241 | 0.020 | 2.526 | 0.0004 |
| HDAC7A | 51564 | 1.241 | 0.005 | 2.581 | 0.0000 |
| ANKK1 | 255239 | 1.243 | 0.059 | 2.200 | 0.0172 |
| RRAGB | 10325 | 1.244 | 0.380 | 0.625 | 0.3808 |
| siERCC1 Control | siERCC1 Contr | 1.246 | 1.243 | 2.633 | 0.0000 |
| TOP3B | 8940 | 1.247 | 0.156 | 1.362 | 0.1101 |
| JJAZ1 | 23512 | 1.248 | 0.257 | 0.906 | 0.2370 |
| ATP11C | 286410 | 1.250 | 0.090 | 1.930 | 0.0395 |
| BRCA2 | 675 | 1.250 | 0.098 | 1.848 | 0.0467 |
| UBE2E2 | 7325 | 1.251 | 0.044 | 2.433 | 0.0079 |
| BMI1 | 648 | 1.251 | 0.106 | 1.774 | 0.0540 |
| TYMS | 7298 | 1.251 | 0.169 | 1.300 | 0.1235 |
| DYRK1B | 9149 | 1.251 | 0.089 | 1.952 | 0.0380 |
| DNMT1 | 1786 | 1.252 | 0.065 | 2.216 | 0.0197 |
| CDC7 | 8317 | 1.254 | 0.065 | 2.235 | 0.0194 |
| PPP1CC | 5501 | 1.254 | 0.152 | 1.425 | 0.1010 |
| DCPS | 28960 | 1.254 | 0.189 | 1.205 | 0.1452 |
| RAD1 | 5810 | 1.255 | 0.139 | 1.525 | 0.0857 |
| SUV39H1 | 6839 | 1.258 | 0.324 | 0.764 | 0.3023 |
| CDC25A | 993 | 1.258 | 0.148 | 1.473 | 0.0942 |
| USP34 | 9736 | 1.258 | 0.067 | 2.248 | 0.0202 |
| CKS2 | 1164 | 1.261 | 0.032 | 2.647 | 0.0027 |
| TOP3A | 7156 | 1.261 | 0.247 | 0.988 | 0.2088 |
| SIRT6 | 51548 | 1.264 | 0.103 | 1.901 | 0.0460 |
| PRKAB1 | 5564 | 1.270 | 0.466 | 0.568 | 0.4214 |

| | | | | | |
|---|---|---|---|---|---|
| C14ORF130 | 55148 | 1.271 | 0.074 | 2.268 | 0.0229 |
| siCHEK1 Control | siCHEK1 Contr | 1.271 | 0.159 | 1.468 | 0.0980 |
| SLK | 9748 | 1.271 | 0.093 | 2.057 | 0.0361 |
| CBL | 867 | 1.273 | 0.190 | 1.286 | 0.1310 |
| RNF14 | 9604 | 1.275 | 0.106 | 1.944 | 0.0454 |
| NME5 | 8382 | 1.277 | 0.047 | 2.663 | 0.0074 |
| FLJ10260 | 55106 | 1.278 | 0.262 | 1.001 | 0.2070 |
| CRK7 | 51755 | 1.279 | 0.069 | 2.411 | 0.0181 |
| PNPT1 | 87178 | 1.279 | 0.200 | 1.267 | 0.1362 |
| FBXL13 | 222235 | 1.280 | 0.136 | 1.695 | 0.0701 |
| ANAPC11 | 51529 | 1.281 | 0.241 | 1.087 | 0.1809 |
| MCM5 | 4174 | 1.283 | 0.085 | 2.238 | 0.0279 |
| UBE2D3 | 7323 | 1.283 | 0.195 | 1.309 | 0.1283 |
| PCK2 | 5106 | 1.284 | 0.007 | 3.034 | 0.0000 |
| HRMT1L6 | 55170 | 1.284 | 0.125 | 1.825 | 0.0579 |
| CDK3 | 1018 | 1.285 | 0.071 | 2.434 | 0.0186 |
| UBE2D2 | 7322 | 1.288 | 0.020 | 3.020 | 0.0002 |
| PRKCD | 5580 | 1.290 | 0.121 | 1.900 | 0.0525 |
| DKC1 | 1736 | 1.290 | 0.105 | 2.063 | 0.0403 |
| PAK2 | 5062 | 1.290 | 0.212 | 1.252 | 0.1411 |
| CUL1 | 8454 | 1.296 | 0.153 | 1.652 | 0.0782 |
| SRC | 6714 | 1.296 | 0.108 | 2.079 | 0.0406 |
| MGC19764 | 162394 | 1.297 | 0.231 | 1.193 | 0.1555 |
| KIAA2002 | 79834 | 1.298 | 0.185 | 1.439 | 0.1076 |
| FBXL6 | 26233 | 1.299 | 0.115 | 2.025 | 0.0448 |
| CETN2 | 1069 | 1.300 | 0.118 | 1.997 | 0.0471 |
| HUWE1 | 10075 | 1.302 | 0.170 | 1.562 | 0.0905 |
| RING1 | 6015 | 1.303 | 0.088 | 2.355 | 0.0263 |
| TREX2 | 11219 | 1.304 | 0.152 | 1.703 | 0.0739 |
| PPP1R12C | 54776 | 1.304 | 0.135 | 1.851 | 0.0595 |
| TA-PP2C | 160760 | 1.304 | 0.090 | 2.349 | 0.0269 |
| TOPORS | 10210 | 1.305 | 0.197 | 1.397 | 0.1155 |
| ERBB2 | 2064 | 1.306 | 0.295 | 0.987 | 0.2148 |
| CHD7 | 55636 | 1.306 | 0.162 | 1.638 | 0.0815 |
| TNKS2 | 80351 | 1.306 | 0.173 | 1.562 | 0.0911 |
| CP | 1356 | 1.308 | 0.222 | 1.279 | 0.1380 |
| NRBP | 29959 | 1.308 | 0.187 | 1.474 | 0.1038 |
| DEPC-1 | 221120 | 1.309 | 0.071 | 2.633 | 0.0161 |
| DKFZP566K0524 | 26095 | 1.313 | 0.111 | 2.162 | 0.0385 |
| WBSCR27 | 155368 | 1.314 | 0.103 | 2.258 | 0.0333 |
| DNASE1L3 | 1776 | 1.315 | 0.402 | 0.763 | 0.3076 |
| FUK | 197258 | 1.315 | 0.186 | 1.519 | 0.0982 |
| ALOX15B | 247 | 1.317 | 0.044 | 3.073 | 0.0049 |

| | | | | | |
|---|---|---|---|---|---|
| RAD51L1 | 5890 | 1.318 | 0.571 | 0.549 | 0.4370 |
| PSMD4 | 5710 | 1.319 | 0.220 | 1.335 | 0.1284 |
| CDC2L1 | 984 | 1.319 | 0.075 | 2.669 | 0.0168 |
| POLA | 5422 | 1.320 | 0.122 | 2.081 | 0.0447 |
| IBRDC2 | 255488 | 1.320 | 0.180 | 1.581 | 0.0906 |
| ERCC3 | 2071 | 1.320 | 0.049 | 3.047 | 0.0061 |
| BRDT | 676 | 1.321 | 0.101 | 2.337 | 0.0306 |
| TMX2 | 51075 | 1.321 | 0.124 | 2.077 | 0.0452 |
| PINK1 | 65018 | 1.322 | 0.166 | 1.688 | 0.0782 |
| LMTK3 | 114783 | 1.325 | 0.118 | 2.158 | 0.0408 |
| SMARCA5 | 8467 | 1.326 | 0.117 | 2.181 | 0.0395 |
| HRMT1L1 | 3275 | 1.326 | 0.154 | 1.811 | 0.0665 |
| NOSIP | 51070 | 1.328 | 0.072 | 2.792 | 0.0143 |
| HINT1 | 3094 | 1.328 | 0.128 | 2.071 | 0.0466 |
| SIAH2 | 6478 | 1.331 | 0.228 | 1.346 | 0.1279 |
| RRM2 | 6241 | 1.331 | 0.140 | 1.974 | 0.0538 |
| USP53 | 54532 | 1.331 | 0.167 | 1.733 | 0.0748 |
| PTK2 | 5747 | 1.335 | 0.286 | 1.114 | 0.1794 |
| USP31 | 57478 | 1.335 | 0.216 | 1.421 | 0.1153 |
| CDKL2 | 8999 | 1.335 | 0.036 | 3.353 | 0.0024 |
| SMC1L1 | 8243 | 1.336 | 0.132 | 2.083 | 0.0469 |
| UBE2D1 | 7321 | 1.336 | 0.097 | 2.501 | 0.0257 |
| LOC283989 | 283989 | 1.336 | 0.215 | 1.435 | 0.1132 |
| DUSP4 | 1846 | 1.337 | 0.221 | 1.403 | 0.1185 |
| ATE1 | 11101 | 1.338 | 0.192 | 1.581 | 0.0928 |
| DUSP11 | 8446 | 1.340 | 0.333 | 0.983 | 0.2190 |
| HDAC10 | 83933 | 1.341 | 0.090 | 2.632 | 0.0215 |
| ZDHHC1 | 29800 | 1.344 | 0.291 | 1.125 | 0.1772 |
| DUSP5 | 1847 | 1.344 | 0.143 | 2.016 | 0.0525 |
| SFRS9 | 8683 | 1.344 | 0.091 | 2.639 | 0.0217 |
| CREBBP | 1387 | 1.346 | 0.290 | 1.136 | 0.1745 |
| UBL4 | 8266 | 1.349 | 0.293 | 1.135 | 0.1751 |
| MTAP | 4507 | 1.350 | 0.211 | 1.516 | 0.1027 |
| RKHD3 | 84206 | 1.351 | 0.018 | 3.697 | 0.0001 |
| MIB2 | 142678 | 1.351 | 0.179 | 1.741 | 0.0763 |
| POP1 | 10940 | 1.352 | 0.294 | 1.141 | 0.1736 |
| TTBK2 | 146057 | 1.352 | 0.088 | 2.748 | 0.0193 |
| POLI | 11201 | 1.358 | 0.163 | 1.900 | 0.0627 |
| PTBP1 | 5725 | 1.361 | 0.091 | 2.765 | 0.0198 |
| FLJ10006 | 55677 | 1.362 | 0.132 | 2.242 | 0.0409 |
| NRK | 203447 | 1.363 | 0.154 | 2.015 | 0.0548 |
| CDK2 | 1017 | 1.373 | 0.044 | 3.622 | 0.0033 |
| POLD3 | 10714 | 1.375 | 0.187 | 1.796 | 0.0734 |

| | | | | | |
|---|---|---|---|---|---|
| HACE1 | 57531 | 1.377 | 0.143 | 2.210 | 0.0442 |
| USP37 | 57695 | 1.378 | 0.291 | 1.237 | 0.1533 |
| PIM3 | 415116 | 1.378 | 0.174 | 1.919 | 0.0633 |
| HEMK1 | 51409 | 1.380 | 0.044 | 3.686 | 0.0033 |
| FLJ22054 | 79612 | 1.380 | 0.187 | 1.818 | 0.0718 |
| DKFZP761G058 | 152926 | 1.380 | 0.212 | 1.643 | 0.0895 |
| TRIM28 | 10155 | 1.380 | 0.154 | 2.112 | 0.0501 |
| USP24 | 23358 | 1.383 | 0.088 | 2.990 | 0.0162 |
| FEM1A | 55527 | 1.383 | 0.210 | 1.667 | 0.0870 |
| FRAP1 | 2475 | 1.385 | 0.014 | 4.077 | 0.0000 |
| FKBP5 | 2289 | 1.388 | 0.142 | 2.283 | 0.0415 |
| PCBD | 5092 | 1.389 | 0.166 | 2.046 | 0.0551 |
| EHMT1 | 79813 | 1.390 | 0.108 | 2.739 | 0.0238 |
| TRIM23 | 373 | 1.390 | 0.216 | 1.655 | 0.0889 |
| HRMT1L2 | 3276 | 1.392 | 0.200 | 1.775 | 0.0767 |
| CLK4 | 57396 | 1.394 | 0.070 | 3.373 | 0.0095 |
| MAPKAPK2 | 9261 | 1.394 | 0.102 | 2.857 | 0.0208 |
| DUSP12 | 11266 | 1.395 | 0.243 | 1.519 | 0.1061 |
| RAD51L3 | 5892 | 1.397 | 0.145 | 2.301 | 0.0413 |
| PLK4 | 10733 | 1.397 | 0.068 | 3.440 | 0.0087 |
| CDK11 | 23097 | 1.400 | 0.171 | 2.054 | 0.0555 |
| OGG1 | 4968 | 1.403 | 0.166 | 2.116 | 0.0518 |
| UGDH | 7358 | 1.404 | 0.013 | 4.294 | 0.0000 |
| TARS | 6897 | 1.405 | 0.191 | 1.908 | 0.0663 |
| ULK4 | 54986 | 1.410 | 0.034 | 4.133 | 0.0013 |
| ATP5B | 506 | 1.410 | 0.124 | 2.651 | 0.0284 |
| FBXO9 | 26268 | 1.411 | 0.041 | 4.026 | 0.0023 |
| PAPD5 | 64282 | 1.413 | 0.197 | 1.894 | 0.0679 |
| PRKCE | 5581 | 1.416 | 0.120 | 2.741 | 0.0260 |
| APTX | 54840 | 1.417 | 0.035 | 4.197 | 0.0012 |
| HTATIP | 10524 | 1.417 | 0.188 | 1.986 | 0.0614 |
| GARNL1 | 253959 | 1.417 | 0.091 | 3.205 | 0.0147 |
| RNF111 | 54778 | 1.420 | 0.164 | 2.222 | 0.0471 |
| PUS1 | 80324 | 1.420 | 0.122 | 2.734 | 0.0265 |
| USP1 | 7398 | 1.422 | 0.085 | 3.344 | 0.0125 |
| G22P1 | 2547 | 1.423 | 0.191 | 1.992 | 0.0613 |
| CDC2L5 | 8621 | 1.423 | 0.173 | 2.156 | 0.0510 |
| RPS6KA2 | 6196 | 1.423 | 0.453 | 0.915 | 0.2470 |
| C6ORF11 | 9277 | 1.423 | 0.193 | 1.976 | 0.0624 |
| FLJ10774 | 55226 | 1.424 | 0.223 | 1.753 | 0.0811 |
| TERT | 7015 | 1.427 | 0.107 | 3.014 | 0.0196 |
| USP45 | 85015 | 1.427 | 0.170 | 2.203 | 0.0486 |
| STK22C | 81629 | 1.429 | 0.199 | 1.949 | 0.0648 |

| | | | | | |
|---|---|---|---|---|---|
| NEK4 | 6787 | 1.429 | 0.179 | 2.124 | 0.0532 |
| MDA5 | 64135 | 1.432 | 0.113 | 2.957 | 0.0213 |
| SIRT2 | 22933 | 1.433 | 0.174 | 2.198 | 0.0493 |
| POP7 | 10248 | 1.435 | 0.007 | 4.652 | 0.0000 |
| LBR | 3930 | 1.435 | 0.144 | 2.532 | 0.0344 |
| CDA | 978 | 1.437 | 0.081 | 3.545 | 0.0104 |
| ARL4D | 379 | 1.439 | 0.150 | 2.480 | 0.0366 |
| ADCK1 | 57143 | 1.440 | 0.147 | 2.530 | 0.0347 |
| BRD4 | 23476 | 1.441 | 0.321 | 1.319 | 0.1403 |
| HDAC1 | 3065 | 1.442 | 0.013 | 4.701 | 0.0000 |
| POLS | 11044 | 1.445 | 0.171 | 2.290 | 0.0453 |
| PSMC1 | 5700 | 1.446 | 0.066 | 3.907 | 0.0064 |
| LASS3 | 204219 | 1.448 | 0.211 | 1.941 | 0.0664 |
| DUB3 | 377630 | 1.448 | 0.200 | 2.031 | 0.0601 |
| PRPS1L1 | 221823 | 1.454 | 0.109 | 3.175 | 0.0179 |
| USP17 | 391627 | 1.459 | 0.275 | 1.585 | 0.1011 |
| SPEG | 10290 | 1.460 | 0.130 | 2.872 | 0.0252 |
| MSH3 | 4437 | 1.461 | 0.195 | 2.137 | 0.0543 |
| CTPS | 1503 | 1.464 | 0.233 | 1.848 | 0.0746 |
| NEDD8 | 4738 | 1.465 | 0.151 | 2.620 | 0.0330 |
| DOT1L | 84444 | 1.467 | 0.042 | 4.558 | 0.0019 |
| USP35 | 57558 | 1.468 | 0.323 | 1.391 | 0.1289 |
| PRIM2A | 5558 | 1.469 | 0.081 | 3.798 | 0.0091 |
| PTP4A3 | 11156 | 1.470 | 0.073 | 3.977 | 0.0072 |
| CHD5 | 26038 | 1.471 | 0.130 | 2.945 | 0.0240 |
| HDAC11 | 79885 | 1.475 | 0.230 | 1.916 | 0.0696 |
| PCGF2 | 7703 | 1.475 | 0.175 | 2.399 | 0.0419 |
| ACP5 | 54 | 1.475 | 0.054 | 4.419 | 0.0034 |
| ATRX | 546 | 1.480 | 0.117 | 3.202 | 0.0188 |
| HECW2 | 57520 | 1.481 | 0.172 | 2.459 | 0.0397 |
| C10ORF89 | 118672 | 1.481 | 0.149 | 2.733 | 0.0302 |
| XRCC3 | 7517 | 1.481 | 0.048 | 4.587 | 0.0025 |
| PIAS1 | 8554 | 1.481 | 0.150 | 2.722 | 0.0306 |
| LIG3 | 3980 | 1.483 | 0.073 | 4.089 | 0.0068 |
| STK32C | 282974 | 1.484 | 0.083 | 3.880 | 0.0090 |
| SET | 6418 | 1.485 | 0.149 | 2.761 | 0.0296 |
| MTA2 | 9219 | 1.488 | 0.490 | 0.979 | 0.2266 |
| ADCK5 | 203054 | 1.489 | 0.146 | 2.823 | 0.0280 |
| DHPS | 1725 | 1.489 | 0.214 | 2.090 | 0.0583 |
| KIAA1008 | 22894 | 1.489 | 0.232 | 1.953 | 0.0675 |
| PHYHD1 | 254295 | 1.489 | 0.156 | 2.692 | 0.0319 |
| FRK | 2444 | 1.489 | 0.045 | 4.721 | 0.0021 |
| DYRK1A | 1859 | 1.491 | 0.051 | 4.620 | 0.0028 |

| | | | | | |
|---|---|---|---|---|---|
| PPM1H | 57460 | 1.492 | 0.174 | 2.495 | 0.0388 |
| RNMT | 8731 | 1.493 | 0.097 | 3.659 | 0.0122 |
| POLA2 | 23649 | 1.494 | 0.166 | 2.588 | 0.0355 |
| BRAP | 8315 | 1.495 | 0.095 | 3.722 | 0.0115 |
| PRIM1 | 5557 | 1.496 | 0.097 | 3.695 | 0.0119 |
| FTSJ2 | 29960 | 1.497 | 0.103 | 3.570 | 0.0136 |
| PAPOLB | 56903 | 1.502 | 0.146 | 2.896 | 0.0267 |
| ISG20 | 3669 | 1.502 | 0.133 | 3.087 | 0.0223 |
| SUMO3 | 6612 | 1.505 | 0.185 | 2.440 | 0.0415 |
| HBXAP | 51773 | 1.506 | 0.124 | 3.270 | 0.0188 |
| DCLRE1C | 64421 | 1.509 | 0.119 | 3.357 | 0.0174 |
| CDYL | 9425 | 1.511 | 0.090 | 3.934 | 0.0097 |
| RECQL5 | 9400 | 1.511 | 0.252 | 1.905 | 0.0719 |
| IBRDC1 | 154214 | 1.511 | 0.051 | 4.808 | 0.0026 |
| NEK2 | 4751 | 1.512 | 0.180 | 2.525 | 0.0386 |
| LATS1 | 9113 | 1.514 | 0.099 | 3.773 | 0.0117 |
| FLJ34922 | 91607 | 1.516 | 0.190 | 2.434 | 0.0422 |
| ART4 | 420 | 1.516 | 0.073 | 4.364 | 0.0059 |
| UFC1 | 51506 | 1.517 | 0.080 | 4.204 | 0.0073 |
| AGTPBP1 | 23287 | 1.519 | 0.007 | 5.550 | 0.0000 |
| POLB | 5423 | 1.523 | 0.120 | 3.445 | 0.0166 |
| RANBP2 | 5903 | 1.524 | 0.115 | 3.540 | 0.0152 |
| UBE2NL | 389898 | 1.525 | 0.347 | 1.462 | 0.1198 |
| UBE4B | 10277 | 1.526 | 0.248 | 1.987 | 0.0665 |
| PSPH | 5723 | 1.527 | 0.138 | 3.159 | 0.0218 |
| CDC34 | 997 | 1.534 | 0.019 | 5.612 | 0.0000 |
| TENS1 | 64759 | 1.535 | 0.204 | 2.384 | 0.0450 |
| CUL3 | 8452 | 1.536 | 0.171 | 2.754 | 0.0319 |
| LPIN2 | 9663 | 1.539 | 0.236 | 2.128 | 0.0580 |
| VRK3 | 51231 | 1.540 | 0.159 | 2.924 | 0.0275 |
| CDK8 | 1024 | 1.541 | 0.129 | 3.398 | 0.0181 |
| TRIM63 | 84676 | 1.544 | 0.271 | 1.901 | 0.0733 |
| RNASEN | 29102 | 1.547 | 0.270 | 1.917 | 0.0722 |
| USP51 | 158880 | 1.552 | 0.080 | 4.489 | 0.0064 |
| WEE1 | 7465 | 1.553 | 0.153 | 3.095 | 0.0241 |
| PKN3 | 29941 | 1.559 | 0.104 | 4.009 | 0.0108 |
| CHD6 | 84181 | 1.559 | 0.117 | 3.735 | 0.0139 |
| HDAC8 | 55869 | 1.560 | 0.092 | 4.278 | 0.0083 |
| UBE2V2 | 7336 | 1.560 | 0.277 | 1.919 | 0.0724 |
| POLD2 | 5425 | 1.562 | 0.305 | 1.762 | 0.0856 |
| WHSC1L1 | 54904 | 1.564 | 0.278 | 1.922 | 0.0723 |
| NRF1 | 4899 | 1.567 | 0.094 | 4.287 | 0.0085 |
| POLK | 51426 | 1.569 | 0.081 | 4.615 | 0.0061 |

| | | | | | |
|---|---|---|---|---|---|
| KIAA1596 | 57697 | 1.578 | 0.425 | 1.329 | 0.1425 |
| PIP5KL1 | 138429 | 1.579 | 0.057 | 5.300 | 0.0026 |
| MEN1 | 4221 | 1.579 | 0.148 | 3.312 | 0.0207 |
| SUMO1 | 7341 | 1.581 | 0.203 | 2.601 | 0.0381 |
| UBE1L | 7318 | 1.581 | 0.255 | 2.140 | 0.0584 |
| RAG2 | 5897 | 1.582 | 0.254 | 2.152 | 0.0578 |
| ASB2 | 51676 | 1.583 | 0.186 | 2.809 | 0.0319 |
| FBXO2 | 26232 | 1.584 | 0.226 | 2.391 | 0.0462 |
| KIAA0317 | 9870 | 1.585 | 0.222 | 2.430 | 0.0446 |
| MIDORI | 57538 | 1.588 | 0.026 | 6.073 | 0.0002 |
| ADCK4 | 79934 | 1.589 | 0.050 | 5.557 | 0.0018 |
| TRRAP | 8295 | 1.591 | 0.296 | 1.902 | 0.0744 |
| HIPK1 | 204851 | 1.594 | 0.263 | 2.127 | 0.0595 |
| U5-116KD | 9343 | 1.596 | 0.941 | 0.630 | 0.3870 |
| CDC14B | 8555 | 1.603 | 0.095 | 4.519 | 0.0078 |
| SIRT1 | 23411 | 1.603 | 0.058 | 5.508 | 0.0025 |
| DHRS2 | 10202 | 1.605 | 0.056 | 5.558 | 0.0023 |
| C9ORF67 | 84814 | 1.605 | 0.181 | 2.967 | 0.0284 |
| CDK6 | 1021 | 1.607 | 0.171 | 3.111 | 0.0253 |
| DUT | 1854 | 1.608 | 0.108 | 4.260 | 0.0100 |
| BUB1B | 701 | 1.611 | 0.259 | 2.217 | 0.0550 |
| UBE2S | 27338 | 1.611 | 0.154 | 3.393 | 0.0203 |
| DKFZP434C131 | 25989 | 1.614 | 0.120 | 4.044 | 0.0121 |
| FBXL7 | 23194 | 1.614 | 0.136 | 3.723 | 0.0157 |
| MCM3 | 4172 | 1.615 | 0.064 | 5.443 | 0.0030 |
| ALKBH2 | 121642 | 1.615 | 0.137 | 3.705 | 0.0159 |
| ADPRTL3 | 10039 | 1.620 | 0.166 | 3.254 | 0.0228 |
| FBXO6 | 26270 | 1.624 | 0.145 | 3.614 | 0.0173 |
| SSTK | 83983 | 1.625 | 0.201 | 2.819 | 0.0326 |
| USP36 | 57602 | 1.626 | 0.184 | 3.031 | 0.0274 |
| RBKS | 64080 | 1.631 | 0.084 | 5.036 | 0.0054 |
| PPP3R2 | 5535 | 1.633 | 0.056 | 5.831 | 0.0020 |
| CLK1 | 1195 | 1.634 | 0.210 | 2.757 | 0.0345 |
| PSMC6 | 5706 | 1.636 | 0.261 | 2.293 | 0.0517 |
| RNF150 | 57484 | 1.636 | 0.050 | 6.021 | 0.0015 |
| IGHMBP2 | 3508 | 1.638 | 0.092 | 4.875 | 0.0064 |
| FTSJ3 | 117246 | 1.639 | 0.218 | 2.690 | 0.0366 |
| FLJ37970 | 283234 | 1.639 | 0.083 | 5.113 | 0.0052 |
| SSB | 6741 | 1.644 | 0.445 | 1.417 | 0.1289 |
| FBXO10 | 26267 | 1.644 | 0.125 | 4.121 | 0.0121 |
| XRN2 | 22803 | 1.645 | 0.132 | 3.985 | 0.0134 |
| NRBP2 | 340371 | 1.648 | 0.340 | 1.836 | 0.0808 |
| FLT4 | 2324 | 1.648 | 0.079 | 5.300 | 0.0045 |

| | | | | | |
|---|---|---|---|---|---|
| MTMR8 | 55613 | 1.650 | 0.259 | 2.359 | 0.0490 |
| PTPN14 | 5784 | 1.650 | 0.344 | 1.825 | 0.0819 |
| EDD1 | 51366 | 1.652 | 0.213 | 2.809 | 0.0334 |
| BPGM | 669 | 1.654 | 0.184 | 3.169 | 0.0252 |
| ARL4C | 10123 | 1.655 | 0.247 | 2.479 | 0.0442 |
| PTPDC1 | 138639 | 1.655 | 0.082 | 5.294 | 0.0047 |
| C1orf48 | 25936 | 1.658 | 0.259 | 2.389 | 0.0479 |
| MX1 | 4599 | 1.659 | 0.169 | 3.409 | 0.0211 |
| GRK7 | 131890 | 1.660 | 0.014 | 7.008 | 0.0000 |
| PPM1L | 151742 | 1.661 | 0.160 | 3.573 | 0.0187 |
| KIAA0625 | 23064 | 1.662 | 0.380 | 1.689 | 0.0947 |
| CDKL5 | 6792 | 1.664 | 0.122 | 4.316 | 0.0108 |
| FBXL19 | 54620 | 1.665 | 0.036 | 6.670 | 0.0005 |
| ADP-GK | 83440 | 1.668 | 0.091 | 5.144 | 0.0057 |
| BLM | 641 | 1.671 | 0.144 | 3.919 | 0.0146 |
| TTBK1 | 84630 | 1.672 | 0.230 | 2.711 | 0.0366 |
| ANAPC2 | 29882 | 1.673 | 0.059 | 6.117 | 0.0020 |
| BPNT1 | 10380 | 1.674 | 0.074 | 5.660 | 0.0036 |
| USP49 | 25862 | 1.674 | 0.128 | 4.256 | 0.0115 |
| HELB | 92797 | 1.678 | 0.344 | 1.905 | 0.0758 |
| METTL3 | 56339 | 1.682 | 0.337 | 1.951 | 0.0725 |
| USP47 | 55031 | 1.684 | 0.183 | 3.331 | 0.0228 |
| PRKY | 5616 | 1.684 | 0.208 | 2.996 | 0.0294 |
| EYA2 | 2139 | 1.684 | 0.141 | 4.050 | 0.0136 |
| PPIL2 | 23759 | 1.687 | 0.181 | 3.377 | 0.0221 |
| FLJ32440 | 286053 | 1.687 | 0.509 | 1.328 | 0.1443 |
| EYA1 | 2138 | 1.690 | 0.098 | 5.110 | 0.0063 |
| MPG | 4350 | 1.690 | 0.468 | 1.447 | 0.1250 |
| USP22 | 23326 | 1.690 | 0.589 | 1.157 | 0.1796 |
| ARD1 | 8260 | 1.691 | 0.222 | 2.867 | 0.0326 |
| MAP3K6 | 9064 | 1.691 | 0.229 | 2.798 | 0.0344 |
| FLJ32332 | 132160 | 1.691 | 0.246 | 2.629 | 0.0395 |
| UNG | 7374 | 1.692 | 0.084 | 5.507 | 0.0045 |
| EP300 | 2033 | 1.697 | 0.158 | 3.796 | 0.0165 |
| POGZ | 23126 | 1.704 | 0.207 | 3.099 | 0.0275 |
| NP | 4860 | 1.709 | 0.148 | 4.060 | 0.0139 |
| MOV10 | 4343 | 1.710 | 0.187 | 3.398 | 0.0221 |
| LOC402682 | 402682 | 1.715 | 0.435 | 1.607 | 0.1044 |
| KIT | 3815 | 1.717 | 0.094 | 5.408 | 0.0054 |
| BUB1 | 699 | 1.720 | 0.225 | 2.962 | 0.0307 |
| POLE4 | 56655 | 1.726 | 0.446 | 1.592 | 0.1062 |
| NCB5OR | 51167 | 1.728 | 0.445 | 1.601 | 0.1052 |
| DUSP10 | 11221 | 1.728 | 0.163 | 3.885 | 0.0160 |

| | | | | | |
|---|---|---|---|---|---|
| MCM4 | 4173 | 1.729 | 0.134 | 4.464 | 0.0108 |
| TOP2B | 7155 | 1.730 | 0.280 | 2.474 | 0.0456 |
| TBDN100 | 80155 | 1.730 | 0.254 | 2.694 | 0.0380 |
| UBE2C | 11065 | 1.730 | 0.021 | 7.638 | 0.0000 |
| SBK1 | 388228 | 1.730 | 0.143 | 4.279 | 0.0123 |
| DUSP8 | 1850 | 1.732 | 0.282 | 2.463 | 0.0460 |
| PANK1 | 53354 | 1.733 | 0.161 | 3.938 | 0.0155 |
| PSMC2 | 5701 | 1.738 | 0.241 | 2.854 | 0.0337 |
| DBC1 | 1620 | 1.742 | 0.087 | 5.817 | 0.0042 |
| ZBED1 | 9189 | 1.744 | 0.042 | 7.276 | 0.0007 |
| WBSCR22 | 114049 | 1.744 | 0.355 | 2.028 | 0.0681 |
| LOC81691 | 81691 | 1.751 | 0.042 | 7.346 | 0.0006 |
| MCM2 | 4171 | 1.753 | 0.098 | 5.560 | 0.0053 |
| GTPBP4 | 23560 | 1.753 | 0.130 | 4.698 | 0.0096 |
| SMARCAL1 | 50485 | 1.757 | 0.471 | 1.578 | 0.1082 |
| PRPF4B | 8899 | 1.760 | 0.235 | 3.005 | 0.0303 |
| BRCA1 | 672 | 1.773 | 0.109 | 5.405 | 0.0062 |
| HUMAUANTIG | 29889 | 1.774 | 0.050 | 7.319 | 0.0010 |
| PLK1 | 5347 | 1.775 | 0.205 | 3.436 | 0.0224 |
| FKBP6 | 8468 | 1.776 | 0.041 | 7.618 | 0.0006 |
| HELIC1 | 10973 | 1.780 | 0.109 | 5.442 | 0.0061 |
| PFTK1 | 5218 | 1.784 | 0.063 | 6.958 | 0.0018 |
| PTPN23 | 25930 | 1.786 | 0.168 | 4.086 | 0.0147 |
| PRODH2 | 58510 | 1.789 | 0.355 | 2.148 | 0.0613 |
| NOS3 | 4846 | 1.793 | 0.417 | 1.854 | 0.0812 |
| DUSP2 | 1844 | 1.795 | 0.174 | 4.031 | 0.0154 |
| PHKG2 | 5261 | 1.802 | 0.258 | 2.924 | 0.0327 |
| CHD1 | 1105 | 1.805 | 0.267 | 2.846 | 0.0346 |
| PSMC4 | 5704 | 1.810 | 0.413 | 1.914 | 0.0766 |
| USP38 | 84640 | 1.812 | 0.285 | 2.711 | 0.0385 |
| MYST1 | 84148 | 1.821 | 0.102 | 5.931 | 0.0049 |
| ITCH | 83737 | 1.827 | 0.134 | 5.068 | 0.0084 |
| PEG3 | 5178 | 1.829 | 0.105 | 5.922 | 0.0050 |
| USP43 | 124739 | 1.838 | 0.162 | 4.486 | 0.0120 |
| AKIP | 54998 | 1.840 | 0.137 | 5.065 | 0.0085 |
| LIG4 | 3981 | 1.848 | 0.892 | 0.946 | 0.2411 |
| PRPS1 | 5631 | 1.852 | 0.217 | 3.603 | 0.0209 |
| MX2 | 4600 | 1.855 | 0.118 | 5.699 | 0.0060 |
| PIN4 | 5303 | 1.863 | 0.300 | 2.748 | 0.0379 |
| PRPF19 | 27339 | 1.881 | 0.305 | 2.765 | 0.0375 |
| PGAM4 | 441531 | 1.890 | 0.141 | 5.255 | 0.0081 |
| MGC8407 | 79012 | 1.890 | 0.669 | 1.318 | 0.1477 |
| NARF | 26502 | 1.893 | 0.069 | 7.709 | 0.0017 |

| | | | | | |
|---|---|---|---|---|---|
| ALS2CR7 | 65061 | 1.894 | 0.187 | 4.279 | 0.0141 |
| HECTD3 | 79654 | 1.899 | 0.137 | 5.426 | 0.0074 |
| HDAC6 | 10013 | 1.900 | 0.082 | 7.240 | 0.0396 |
| LRRC29 | 26231 | 1.911 | 0.126 | 5.807 | 0.0061 |
| LEPRE1 | 64175 | 1.915 | 0.180 | 4.506 | 0.0125 |
| GAK | 2580 | 1.920 | 0.167 | 4.803 | 0.0107 |
| CDK9 | 1025 | 1.921 | 0.071 | 7.881 | 0.0017 |
| U5-200KD | 23020 | 1.934 | 0.502 | 1.829 | 0.0843 |
| PPP1R8 | 5511 | 1.944 | 0.110 | 6.550 | 0.0043 |
| APEX2 | 27301 | 1.952 | 0.123 | 6.184 | 0.0053 |
| SENP3 | 26168 | 1.955 | 0.154 | 5.296 | 0.0084 |
| RPP30 | 10556 | 1.960 | 0.631 | 1.505 | 0.1189 |
| RPP14 | 11102 | 1.963 | 0.533 | 1.780 | 0.0886 |
| STK32A | 202374 | 1.968 | 0.423 | 2.233 | 0.0582 |
| RIOK2 | 55781 | 1.972 | 0.211 | 4.210 | 0.0152 |
| FKBP3 | 2287 | 1.976 | 0.172 | 4.991 | 0.0100 |
| RRP22 | 10633 | 1.978 | 0.114 | 6.642 | 0.0043 |
| RPP21 | 79897 | 1.995 | 0.442 | 2.201 | 0.0599 |
| CASP10 | 843 | 2.003 | 0.220 | 4.202 | 0.0155 |
| RPP40 | 10799 | 2.004 | 0.094 | 7.581 | 0.0027 |
| PPAPDC2 | 403313 | 2.005 | 0.211 | 4.365 | 0.0142 |
| PPIL4 | 85313 | 2.006 | 0.170 | 5.193 | 0.0092 |
| UCK1 | 83549 | 2.007 | 0.304 | 3.166 | 0.0290 |
| PPM1B | 5495 | 2.007 | 0.071 | 8.590 | 0.0014 |
| AURKA | 6790 | 2.011 | 0.072 | 8.569 | 0.0015 |
| ITPK1 | 3705 | 2.018 | 0.247 | 3.859 | 0.0189 |
| PRDM1 | 639 | 2.023 | 0.270 | 3.580 | 0.0224 |
| ANAPC5 | 51433 | 2.034 | 0.148 | 5.907 | 0.0066 |
| NTHL1 | 4913 | 2.034 | 0.171 | 5.318 | 0.0088 |
| PPIG | 9360 | 2.043 | 0.165 | 5.491 | 0.0081 |
| NEK7 | 140609 | 2.047 | 0.130 | 6.555 | 0.0049 |
| TOP2A | 7153 | 2.048 | 0.361 | 2.815 | 0.0372 |
| PPP1R3D | 5509 | 2.087 | 0.269 | 3.814 | 0.0198 |
| ELL | 8178 | 2.088 | 0.339 | 3.094 | 0.0308 |
| RCHY1 | 25898 | 2.112 | 0.217 | 4.713 | 0.0123 |
| FBXO24 | 26261 | 2.118 | 0.243 | 4.304 | 0.0152 |
| CHEK1 | 1111 | 2.127 | 0.210 | 4.907 | 0.0113 |
| DFFA | 1676 | 2.136 | 0.320 | 3.407 | 0.0254 |
| CDC2 | 983 | 2.140 | 0.033 | 11.525 | 0.0001 |
| GSK3B | 2932 | 2.142 | 0.083 | 9.147 | 0.0016 |
| APOBEC3G | 60489 | 2.151 | 0.146 | 6.656 | 0.0052 |
| ELAC2 | 60528 | 2.165 | 0.371 | 3.047 | 0.0321 |
| SIK3 | 23387 | 2.172 | 0.099 | 8.642 | 0.0022 |

| | | | | | |
|---|---|---|---|---|---|
| PAK7 | 57144 | 2.173 | 0.109 | 8.198 | 0.0027 |
| FBXW11 | 23291 | 2.183 | 0.385 | 2.986 | 0.0335 |
| G6PC3 | 92579 | 2.203 | 0.142 | 7.076 | 0.0045 |
| TTK | 7272 | 2.205 | 0.214 | 5.165 | 0.0102 |
| IPMK | 253430 | 2.226 | 0.252 | 4.563 | 0.0137 |
| CDC2L2 | 728642 | 2.250 | 0.059 | 11.345 | 0.0006 |
| SIPL | 55256 | 2.254 | 0.343 | 3.531 | 0.0239 |
| RPP38 | 10557 | 2.255 | 0.331 | 3.645 | 0.0224 |
| DFFB | 1677 | 2.275 | 0.129 | 7.992 | 0.0033 |
| PTP4A1 | 7803 | 2.292 | 0.182 | 6.328 | 0.0064 |
| MLL3 | 58508 | 2.296 | 0.295 | 4.187 | 0.0168 |
| RCL1 | 10171 | 2.297 | 0.270 | 4.537 | 0.0141 |
| CRY1 | 1407 | 2.301 | 0.290 | 4.272 | 0.0161 |
| SLU7 | 10569 | 2.304 | 0.375 | 3.374 | 0.0264 |
| SND1 | 27044 | 2.314 | 0.499 | 2.586 | 0.0449 |
| DDX48 | 9775 | 2.333 | 0.337 | 3.810 | 0.0206 |
| WBP11 | 51729 | 2.344 | 0.132 | 8.302 | 0.0031 |
| DNAJA2 | 10294 | 2.361 | 0.810 | 1.671 | 0.1004 |
| RUVBL2 | 10856 | 2.365 | 0.083 | 10.944 | 0.0011 |
| TBCK | 93627 | 2.431 | 0.144 | 8.357 | 0.0032 |
| RAN | 5901 | 2.434 | 0.753 | 1.891 | 0.0808 |
| PPP1R15B | 84919 | 2.483 | 0.284 | 4.956 | 0.0120 |
| DMC1 | 11144 | 2.536 | 0.544 | 2.783 | 0.0393 |
| PRKAG3 | 53632 | 2.551 | 0.291 | 5.070 | 0.0115 |
| TAF5 | 6877 | 2.557 | 0.586 | 2.622 | 0.0441 |
| CIB2 | 10518 | 2.623 | 0.125 | 10.424 | 0.0019 |
| GPS1 | 2873 | 2.725 | 0.124 | 11.131 | 0.0016 |

est)

Positive Hits from Primary Etoposide Hypersensitivity siRNA Screen (62 hits)

| Gene symbol | Gene ID | Average Treated/Untreated Viability (Normalized to NS = 1) (this was threshold) | Standard Deviation (n=3) | SSMD | P-Value (Two-Tailed T-Test) |
|---|---|---|---|---|---|
| FN3K | 64122 | 0.616 | 0.018 | -4.040 | 0.0000 |
| CETN1 | 1068 | 0.273 | 0.030 | -7.433 | 0.0002 |
| CDKL4 | 344387 | 0.532 | 0.024 | -4.864 | 0.0002 |
| SETDB2 | 83852 | 0.655 | 0.022 | -3.604 | 0.0003 |
| CDKL3 | 51265 | 0.446 | 0.055 | -5.120 | 0.0026 |
| ANAPC10 | 10393 | 0.592 | 0.046 | -3.923 | 0.0032 |
| UNG2 | 10309 | 0.675 | 0.039 | -3.213 | 0.0033 |
| TGFBR1 | 7046 | 0.614 | 0.045 | -3.723 | 0.0034 |
| FGR | 2268 | 0.688 | 0.040 | -3.077 | 0.0039 |
| LYK5 | 92335 | 0.366 | 0.073 | -5.366 | 0.0039 |
| LIPI | 149998 | 0.492 | 0.063 | -4.530 | 0.0043 |
| MAP4K3 | 8491 | 0.518 | 0.061 | -4.318 | 0.0046 |
| LMTK2 | 22853 | 0.504 | 0.065 | -4.356 | 0.0050 |
| SRPK1 | 6732 | 0.618 | 0.053 | -3.561 | 0.0053 |
| HNRPAB | 3182 | 0.530 | 0.064 | -4.155 | 0.0054 |
| SUMO2 | 6613 | 0.512 | 0.067 | -4.265 | 0.0054 |
| CERS2 | 29956 | 0.560 | 0.064 | -3.881 | 0.0062 |
| FBXL18 | 80028 | 0.470 | 0.078 | -4.374 | 0.0065 |
| SMUG1 | 23583 | 0.589 | 0.064 | -3.642 | 0.0070 |
| HELZ | 9931 | 0.665 | 0.055 | -3.101 | 0.0075 |
| RNF8 | 9025 | 0.551 | 0.074 | -3.774 | 0.0082 |
| FBXL20 | 84961 | 0.612 | 0.065 | -3.410 | 0.0083 |
| CA9 | 768 | 0.561 | 0.074 | -3.689 | 0.0085 |
| PTGS2 | 5743 | 0.446 | 0.093 | -4.205 | 0.0088 |
| STK33 | 65975 | 0.600 | 0.071 | -3.410 | 0.0095 |
| DYRK3 | 8444 | 0.694 | 0.056 | -2.808 | 0.0097 |
| CTDSP1 | 58190 | 0.606 | 0.075 | -3.295 | 0.0109 |
| ACPT | 93650 | 0.546 | 0.090 | -3.498 | 0.0123 |
| TXNL4 | 10907 | 0.463 | 0.106 | -3.800 | 0.0123 |
| SUMO4 | 387082 | 0.474 | 0.105 | -3.752 | 0.0125 |
| ADK | 132 | 0.631 | 0.076 | -3.072 | 0.0128 |
| ADAR | 103 | 0.606 | 0.083 | -3.160 | 0.0136 |
| PNKP | 11284 | 0.501 | 0.105 | -3.551 | 0.0140 |
| SENP7 | 57337 | 0.529 | 0.100 | -3.450 | 0.0140 |
| ABL1 | 25 | 0.565 | 0.106 | -3.092 | 0.0184 |
| HIPK2 | 28996 | 0.437 | 0.138 | -3.385 | 0.0190 |
| MATK | 4145 | 0.528 | 0.117 | -3.154 | 0.0194 |
| CDK10 | 8558 | 0.589 | 0.103 | -2.964 | 0.0195 |
| HUS1 | 3364 | 0.529 | 0.118 | -3.143 | 0.0196 |
| HERC4 | 26091 | 0.530 | 0.119 | -3.116 | 0.0201 |

| | | | | | |
|---|---|---|---|---|---|
| CBX8 | 57332 | 0.530 | 0.119 | -3.105 | 0.0203 |
| HELLS | 3070 | 0.657 | 0.089 | -2.661 | 0.0208 |
| ZMPSTE24 | 10269 | 0.506 | 0.129 | -3.097 | 0.0217 |
| SARS | 6301 | 0.680 | 0.089 | -2.489 | 0.0237 |
| PUS3 | 83480 | 0.655 | 0.096 | -2.580 | 0.0239 |
| UCHL5 | 51377 | 0.676 | 0.092 | -2.482 | 0.0246 |
| RAD9A | 5883 | 0.632 | 0.107 | -2.596 | 0.0262 |
| NPEPPS | 9520 | 0.694 | 0.093 | -2.324 | 0.0284 |
| PPP1R7 | 5510 | 0.627 | 0.113 | -2.548 | 0.0285 |
| PAPOLA | 10914 | 0.567 | 0.131 | -2.689 | 0.0288 |
| URKL1 | 54963 | 0.686 | 0.098 | -2.329 | 0.0298 |
| DKFZP761P04 | 157285 | 0.400 | 0.187 | -2.866 | 0.0308 |
| KIAA1333 | 55632 | 0.622 | 0.120 | -2.488 | 0.0314 |
| RIPK2 | 8767 | 0.631 | 0.118 | -2.448 | 0.0320 |
| HDAC3 | 8841 | 0.562 | 0.141 | -2.596 | 0.0322 |
| RNASEH1 | 246243 | 0.669 | 0.107 | -2.332 | 0.0323 |
| DDX26 | 26512 | 0.590 | 0.134 | -2.508 | 0.0335 |
| HRMT1L3 | 10196 | 0.569 | 0.148 | -2.464 | 0.0367 |
| CTDP1 | 9150 | 0.684 | 0.110 | -2.199 | 0.0370 |
| SENP2 | 59343 | 0.470 | 0.190 | -2.511 | 0.0398 |
| EHMT2 | 10919 | 0.511 | 0.179 | -2.424 | 0.0415 |
| P101-PI3K | 23533 | 0.527 | 0.177 | -2.359 | 0.0435 |
